# Causal Evidence for Left DLPFC Contributions to Working Memory, Attention, and Cognitive Control in Ageing: A cTBS–ERP Study

**DOI:** 10.64898/2026.01.15.699654

**Authors:** Esteban León-Correa, Alex Balani, Adam Qureshi, Dorothy Tse, Stergios Makris

**Affiliations:** Department of Psychology, Edge Hill University, Ormskirk, UK; Department of Theoretical and Applied Sciences, eCampus University, Novedrate, Italy; IRCCS Centro Neurolesi Bonino-Pulejo, Messina, Italy

**Author notes:** Corresponding author: Prof. Stergios Makris.

**Keywords:** Ageing, DLPFC, cTBS, ERP, working memory, N-back

## Abstract

Ageing is characterised by progressive neurodegeneration and marked declines in working memory (WM), a domain critically dependent on the integrity of the dorsolateral prefrontal cortex (DLPFC). Although non-invasive brain stimulation is widely used to target this region, the causal contribution of the DLPFC to WM performance in older adults remains insufficiently understood. Here, we applied continuous theta burst stimulation (cTBS) to transiently inhibit the left DLPFC in twenty healthy older adults and assessed the behavioural and neurophysiological consequences during verbal and visuospatial N-back tasks. Behaviourally, moderate learning effects emerged exclusively in the verbal task following vertex stimulation, whereas learning after DLPFC stimulation remained inconclusive. Performance in the visuospatial task showed no reliable evidence of learning across conditions, consistent with greater interindividual variability and age-related decline in spatial WM. Neurophysiologically, ERP amplitudes were stable across conditions; however, marked latency perturbations were observed. In the verbal task, P200 and N200 latencies robustly predicted performance, with P200 latency significantly prolonged after DLPFC inhibition, indicating disrupted early attentional gating. N200 and P300 latency modulations further suggested load-dependent compensatory strategies, reflecting adaptive slowing of cognitive control processes. In contrast, visuospatial performance was not reliably predicted by P200 latency, although N200 and P300 latencies remained informative, indicating domain-specific differences in DLPFC involvement. Taken together, these findings provide causal evidence that left DLPFC disruption selectively interferes with the temporal coordination of verbal WM, emphasising the central role of processing speed and compensatory dynamics in cognitive ageing. ERP latency markers emerge as sensitive indices of subtle stimulation-induced perturbations in older adults.

## INTRODUCTION

Ageing is accompanied by numerous challenges, including neurodegeneration and declines across multiple cognitive domains (Funahashi, 2017; Grady, 2012; Salthouse, 2011; Wyss-Coray, 2016; Ziaei et al., 2017). One of the most affected functions is working memory (WM), defined as the capacity to encode, retain, and manipulate information temporarily in support of goal-directed behaviour (Baddeley, 1998, 2003). The dorsolateral prefrontal cortex (DLPFC) plays a pivotal role in WM, particularly in executive control, information integration, maintenance and manipulation, updating, and decision-making (Jimura et al., 2018; Kim et al., 2015; Moore et al., 2013; Murty et al., 2011; Osaka et al., 2003; Rodriguez Merzagora et al., 2014; Vartanian et al., 2013). However, ageing is associated with both structural and functional decline in the DLPFC—such as reductions in white matter integrity, cortical thickness, and grey matter volume—which are closely linked to impairments in WM performance (Golestani et al., 2014; Lemaitre et al., 2012; Liu et al., 2017; Nissim et al., 2017; Salat et al., 1999, 2001, 2004; Storsve et al., 2014; Wyss-Coray, 2016).

Despite these declines, evidence indicates that older adults can recruit compensatory mechanisms to offset neurodegenerative changes and preserve cognitive performance (Cabeza et al., 2018; Festini et al., 2018; Reuter-Lorenz & Cappell, 2008; Reuter-Lorenz & Park, 2014). Prominent compensatory patterns include bilateral frontal activation (HAROLD model; Cabeza, 2002; Cabeza et al., 2002; Tucker & Stern, 2011), a posterior-to-anterior shift in activation (PASA; Davis et al., 2008; Dennis & Cabeza, 2011), and increased frontal recruitment under higher task demands (CRUNCH; Festini et al., 2018; Reuter-Lorenz & Cappell, 2008). This dynamic interplay between neurodegeneration and compensation raises an important question: does the DLPFC continue to support WM in older adults to the same extent as it does in younger individuals?

This question is especially relevant given that many non-invasive brain stimulation (NIBS) studies have targeted the DLPFC to enhance WM in ageing. Yet, findings from these studies have been inconsistent, with several reporting limited or highly variable effects (Goldsworthy et al., 2021; Goldthorpe et al., 2020; Indahlastari et al., 2021; Siegert et al., 2021). Such variability likely reflects age-related alterations in DLPFC functioning and its integration within broader neural networks, underscoring the need for approaches capable of directly probing its causal contribution to WM.

Continuous Theta Burst Stimulation (cTBS) provides such an approach. As a form of non-invasive brain stimulation, cTBS transiently modulates cortical excitability and allows causal examination of specific brain regions and their cognitive functions. It consists of bursts of magnetic pulses that mimic theta–gamma oscillatory patterns (Huang et al., 2005), inducing inhibitory effects on cortical excitability and effectively generating a transient “virtual lesion” (Di Lazzaro et al., 2005). cTBS is particularly advantageous because it can be administered in under a minute (20–40 seconds) and its effects persist for up to 50 minutes (Huang et al., 2005; Rounis & Huang, 2020; Wischnewski & Schutter, 2015).

Previous studies in younger adults indicate that the behavioural effects of cTBS over the DLPFC on WM are highly variable and dependent on task difficulty. For example, Schicktanz et al. (2015) and Ngetich et al. (2021) found impairments specifically in the 2-back task, whereas Viejo-Sobera et al. (2017) reported impairments only in the 3-back. Vékony et al. (2018) observed no direct impairments but noted a suppression of typical practice-related improvements in both 2-back and 3-back tasks. Evidence in older adults is even more limited: Debarnot et al. (2015) reported no behavioural effect of cTBS during a prospective memory task. Reaction-time results have also been inconsistent, with some studies reporting no effects (Ngetich et al., 2021; Vékony et al., 2018), and others demonstrating disruption of learning-related RT improvements (Viejo-Sobera et al., 2017). Overall, these findings suggest that cTBS outcomes are not uniform and may depend on task difficulty, cognitive load, and interindividual variation in neural responsiveness.

Importantly, none of these previous investigations has combined cTBS with neurophysiological recording to evaluate its impact on prefrontal functioning within a WM context. Although some studies have used EEG, these were outside WM paradigms and yielded mixed results: for example, Chung et al. (2017) reported reductions in theta-band power without ERP changes, whereas Grossheinrich et al. (2009) found no effects on resting-state EEG.

To our knowledge, no prior study has integrated cTBS with neurophysiological measures to investigate the causal role of the DLPFC in WM among older adults. The present study addresses this gap by examining whether inducing a virtual lesion in the left DLPFC through cTBS affects WM performance in healthy older adults. We employed EEG to assess neurophysiological after-effects during a well-established N-back paradigm across two cognitive loads (2-back and 3-back) and two modalities (verbal-auditory and visuospatial), enabling investigation of potential hemispheric lateralisation (Ngetich et al., 2020).

## METHODS

### Participants

Twenty-nine healthy older adults (16 women; M = 71.61, SD = 4.62, range = 64–80 years) were recruited from the local community through public advertisements. Exclusion criteria were left-handedness; metal implants in the head or skull; implanted electronic devices (e.g., pacemakers); psychiatric or neurological conditions (including epilepsy or seizures); a history of alcohol or drug abuse; prior surgical procedures involving the head or spinal cord; and medications that might impair cognitive function (e.g., those affecting attention, memory, or causing fatigue). Eligibility was initially assessed via telephone screening. Suitable participants were invited to the School of Psychology, Edge Hill University, where they completed all testing sessions.

### Experimental procedure

The study consisted of three sessions over three weeks, spaced at least seven days apart to minimise practice and carry-over effects. In Session 1, participants underwent further eligibility assessments, including the Montreal Cognitive Assessment (MoCA), the Geriatric Depression Scale (GDS), and a medical questionnaire. Vision and hearing were also assessed. Participants scoring below 26 on the MoCA, above 5 on the GDS, or exhibiting medical or sensory issues relevant to the exclusion criteria were not included (two exclusions).

Eligible participants provided written informed consent and completed baseline EEG recordings while performing the verbal-auditory and visuospatial N-back tasks (2-back and 3-back). Task order was counterbalanced, and extensive training ensured comprehension. No stimulation was applied in this session.

In Sessions 2 and 3, participants repeated the N-back tasks following continuous theta burst stimulation (cTBS). Stimulation (40 s) was delivered either over the vertex (Cz; 10–10 system) or the left DLPFC (F3), counterbalanced across sessions and participants. Individual motor thresholds were established in Session 2 and verified in Session 3. EEG was recorded during all post-stimulation tasks.

Before each session, participants completed a brief questionnaire on sleep, medication, caffeine, alcohol, and fatigue. One participant withdrew due to discomfort and three for personal reasons. Ethical approval was granted by the Edge Hill University Research Ethics Committee.

### N-back

Two versions of the N-back were administered: a verbal-auditory task and a visuospatial task. Both were implemented in PsychoPy with minor adaptations. Task order was counterbalanced.

In the verbal task, participants heard spoken consonants (C, F, H, J, L, N, K, P, Q, R, V, X) while fixating on a central cross, responding “a” for targets and “l” for non-targets. In the visuospatial task, a white square was presented for 50 ms in one of eight screen locations; responses followed the same target/non-target rule.

Each task comprised 16 blocks of 20 trials (30% targets), with eight 2-back blocks followed by eight 3-back blocks. Participants completed practice blocks (1-, 2-, and 3-back) with feedback; no feedback was provided during experimental trials.

### TMS protocol

TMS was delivered using a MagStim Rapid 2 stimulator. Stimulation intensity was based on each participant’ s motor threshold (MT), determined from single pulses applied at ≤0.5 Hz to the left primary motor cortex (C3). MT was defined as the minimum intensity eliciting motor-evoked potentials in the abductor pollicis brevis muscle in at least five out of ten trials. This ensured stimulation was physiologically calibrated to individual variability.

Stimulation sites were located using the 10–10 EEG system. The vertex (Cz) served as the control site, as stimulation here does not significantly alter functional connectivity. The experimental site was the left DLPFC (F3).

cTBS was delivered at 80% of MT, consisting of 600 pulses in 50 Hz triplet bursts every 200 ms for 40 seconds. This protocol reliably induces inhibitory effects lasting up to 50 minutes, covering the duration of the post-stimulation tasks.

### EEG data acquisition

EEG was recorded using a 64-channel actiCAP system and actiCHamp amplifier (Brain Products GmbH), sampled at 1000 Hz. Electrodes were positioned according to the 10–10 system and referenced online to Cz. Impedance was maintained below 8 kΩ. Continuous EEG was acquired with BrainVision Recorder during all tasks.

### Behavioural data analysis

Two primary behavioural metrics were extracted for each task: accuracy and reaction times (RTs). RTs were calculated as the median response time for correct trials, separately for each stimulus type (target and non-target) and each difficulty level (2-back and 3-back). Accuracy was assessed using the sensitivity index d′, which provides a robust measure of participants’ ability to discriminate between target and non-target trials (Haatveit et al., 2010).

To compute d′, the following raw scores were first obtained: the number of hits (correct responses to target stimuli), misses (incorrect non-responses to targets), correct rejections (correct responses to non-targets), and false alarms (incorrect responses to non-targets). The d′ score was then calculated using the formula: d′ = z (Hit Rate) – z (False Alarm Rate) (Macmillan & Creelman, 1990), where z denotes the inverse of the standard normal cumulative distribution function. The Hit Rate was calculated as hits / (hits + misses), and the False Alarm Rate as false alarms / (false alarms + correct rejections). For extreme values (i.e., perfect scores or zero values), adjusted rates were used as follows:

- For perfect Hit Rates: 1 – 1 / [2 × (hits + misses)]
- For perfect Correct Rejections: 1 – 1 / [2 × (false alarms + correct rejections)]
- For zero Misses or False Alarms: 1 / [2 × total relevant trials]

Response bias was calculated using the criterion β, defined as: β = –½ × [z (Hit Rate) + z (False Alarm Rate)] (Lynn & Barrett, 2014). Higher values indicate a more conservative response style (i.e., fewer hits and fewer false alarms), while lower values suggest a more liberal response tendency (i.e., more hits and more false alarms).

### EEG data pre-processing

EEG data were pre-processed using the EEGLAB toolbox (Delorme & Makeig, 2004) in MATLAB (R2023b, The MathWorks, USA). Data were resampled at 512 Hz, band-pass filtered between 0.5–30 Hz, and a 20 Hz artefact was removed using the *cleanline* plugin. Poor-quality segments and bad channels were identified via visual inspection and the *pop_clean_rawdata* function if they met any of the following criteria: 1) flat signal for ≥ 5 seconds; 2) correlation < 0.8 with neighbouring channels; or 3) amplitudes exceeding ±4 standard deviations from the mean.

Artefacts from eye blinks and muscle activity were corrected using independent component analysis (ICA). Prior to ICA, data were downsampled to 256 Hz and high pass filtered at 1 Hz. ICA components were removed based on the output of the *pop_iclabel* function if they met one of the following criteria: 1) > 80% muscular or ocular activity; or 2) < 10% brain activity.

Rejected channels were then interpolated, and data were re-referenced to average. Epochs (−200ms to 1000ms relative to stimulus onset) were extracted separately for target and non-target trials, baseline-corrected using the 200ms pre-stimulus period, and visually inspected for residual artefacts.

### ERP analysis

ERP analyses were conducted using the Fieldtrip toolbox (Oostenveld et al., 2011). Participants with fewer than 15 valid trials per stimulus type at any difficulty level were excluded (n = 2), yielding a final sample of 20.

Time windows for ERP components were identified by visual inspection of grand-average waveforms from midline electrodes (Fz, Cz, Pz). For the verbal N-back, analyses focused on Fz and Cz (see Figures 3 and 4); for the visuospatial N-back, the P200 and N200 were examined at Fz and Cz, and the P300 at Cz and Pz (see Figures 5 and 6). This data-driven approach ensured that selected windows reflected true ERP morphology, minimising interindividual variability.

**Figure 1.**
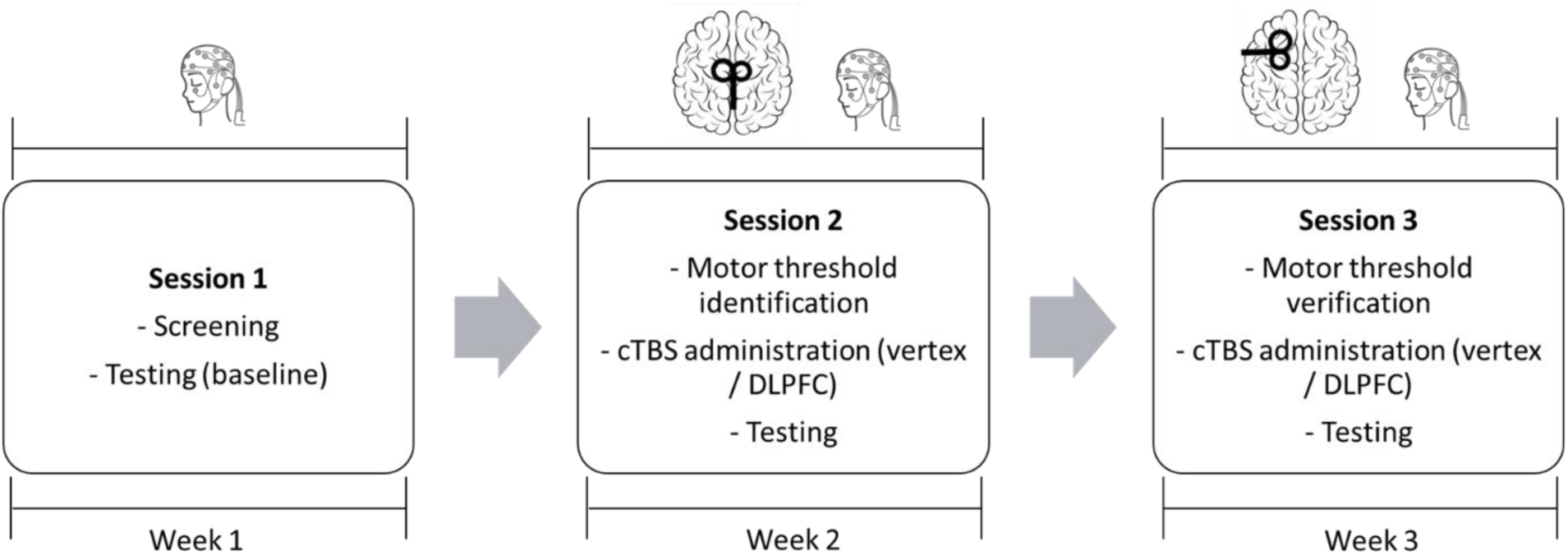
Study Design. Participants completed three sessions across three weeks: baseline, cTBS-vertex and cTBS-DLPFC. In all sessions EEG data were recorded during testing. Stimulation site order was counterbalanced across participants. In this example, it can be observed in the graphs that the coil was placed over the vertex in session 2 and over the left DLPFC in session 3.

**Figure 2.**
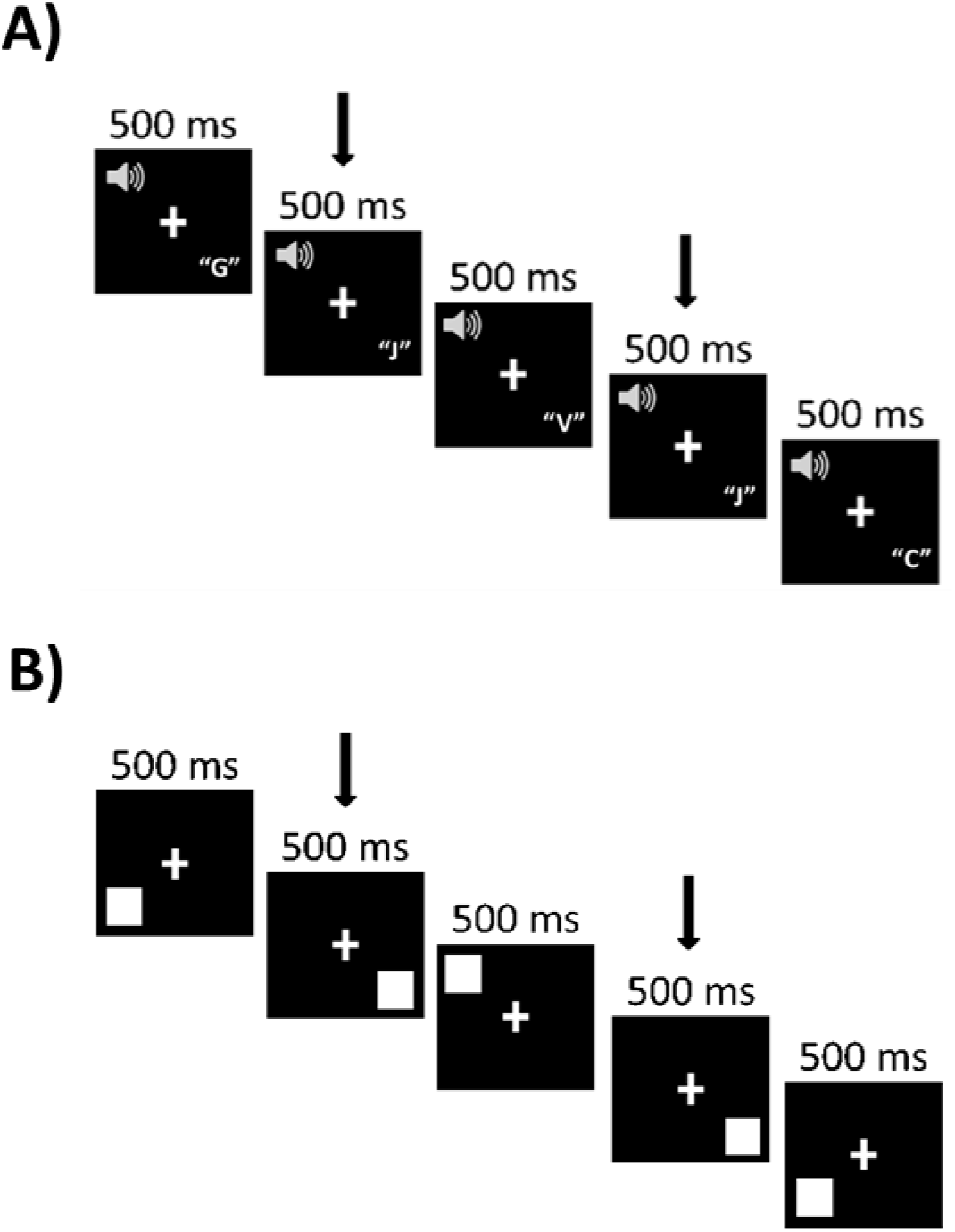
Example of 2-back trials. The arrows show the stimuli that must be compared. A) In this case, the participant would have to press “a” because the letter “J” in the fourth trial was the same as two trials ago. B) In this case, the participant would have to press “a” because the fourth square appeared in the same location as two trials ago.

**Figure 3.**
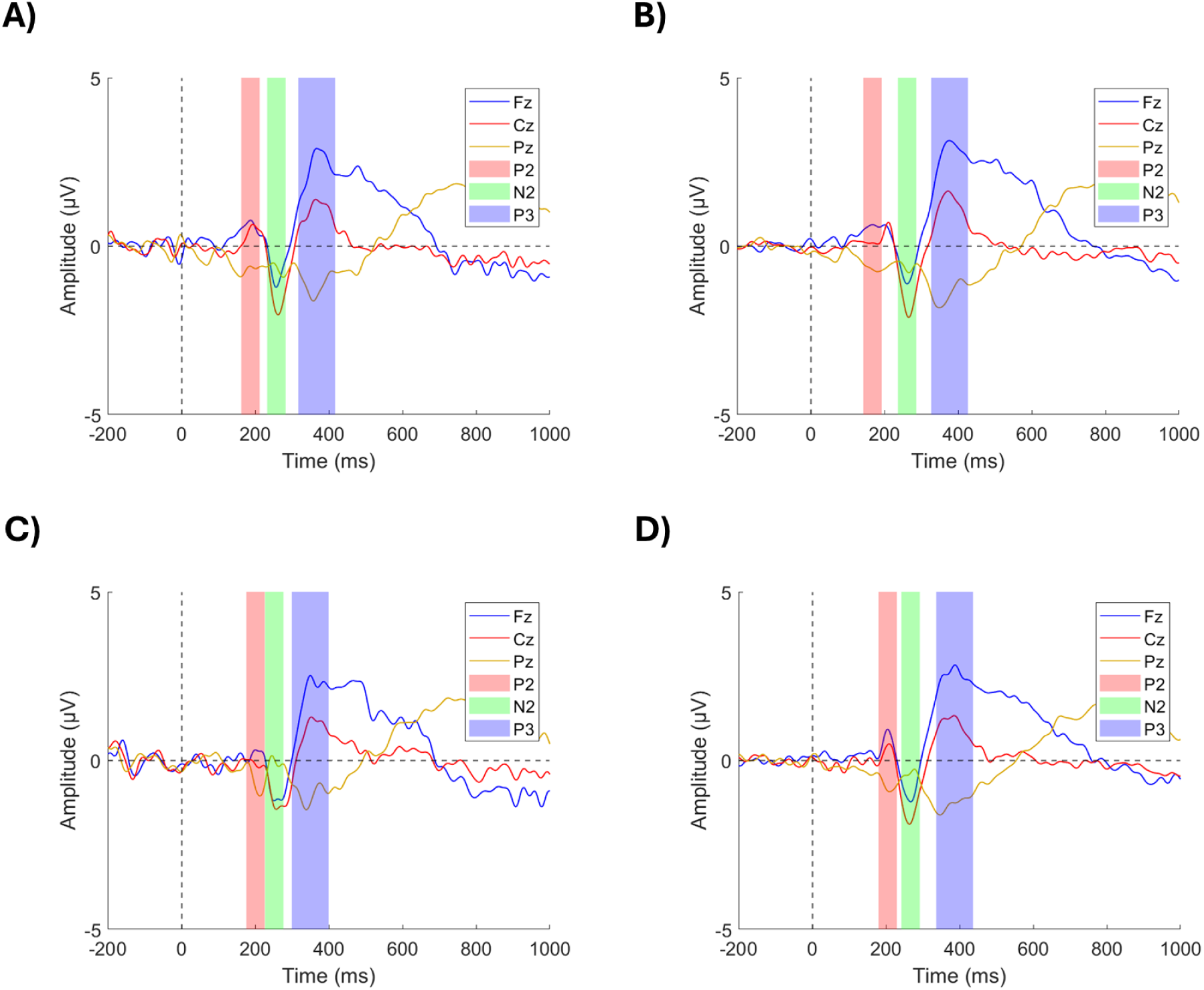
ERPs of the grand average of channels Fz and Cz used to identify the time windows for averaging during the cTBS-vertex condition of the verbal N-back. A) 2-back target trials. B) 2-back non-target trials. C) 3-back target trials. D) 3-back non-target trials.

**Figure 4.**
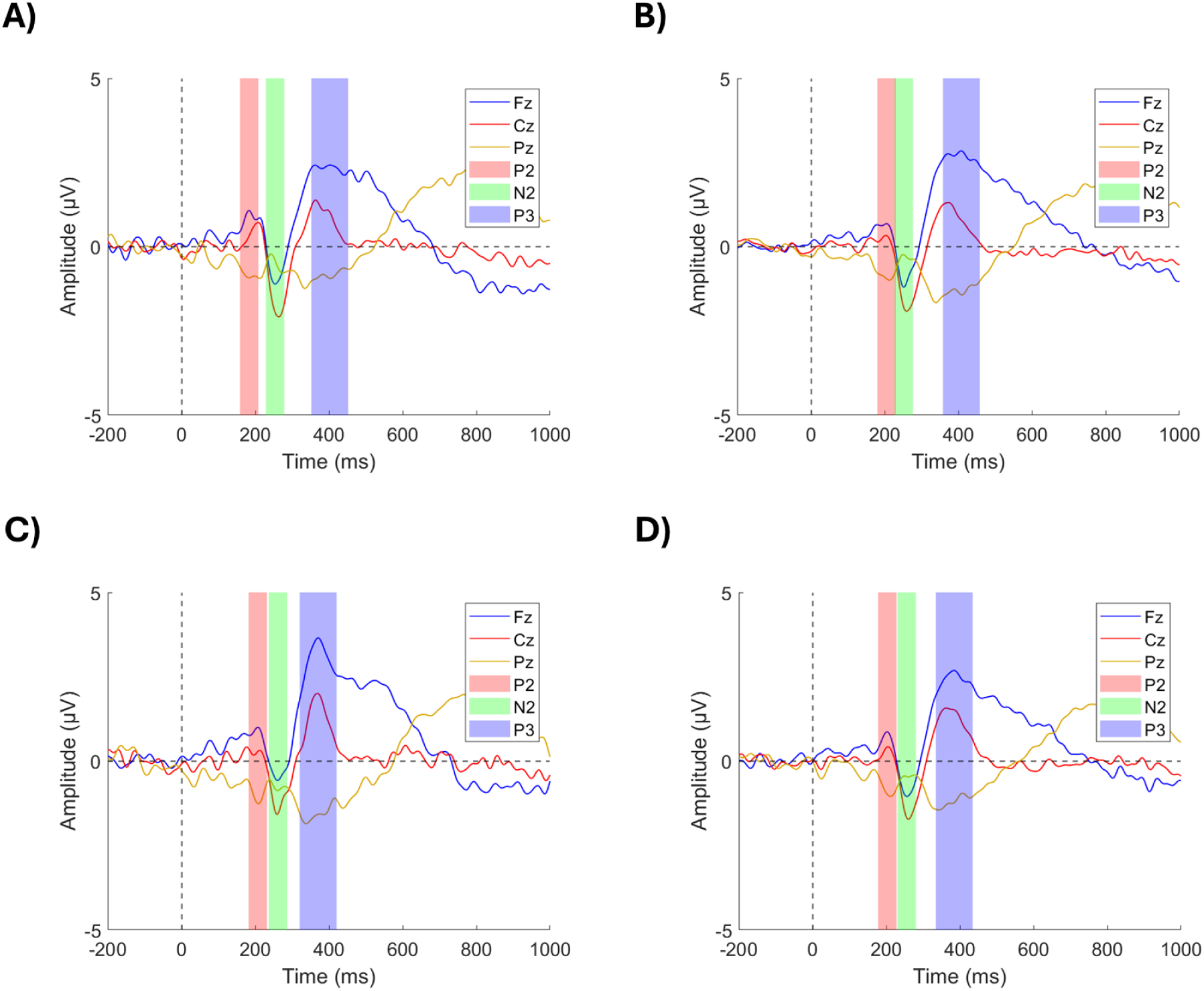
ERPs of the grand average of channels Fz and Cz used to identify the time windows for averaging during the cTBS-DLPFC condition of the verbal N-back. A) 2-back target trials. B) 2-back non-target trials. C) 3-back target trials. D) 3-back non-target trials.

**Figure 5.**
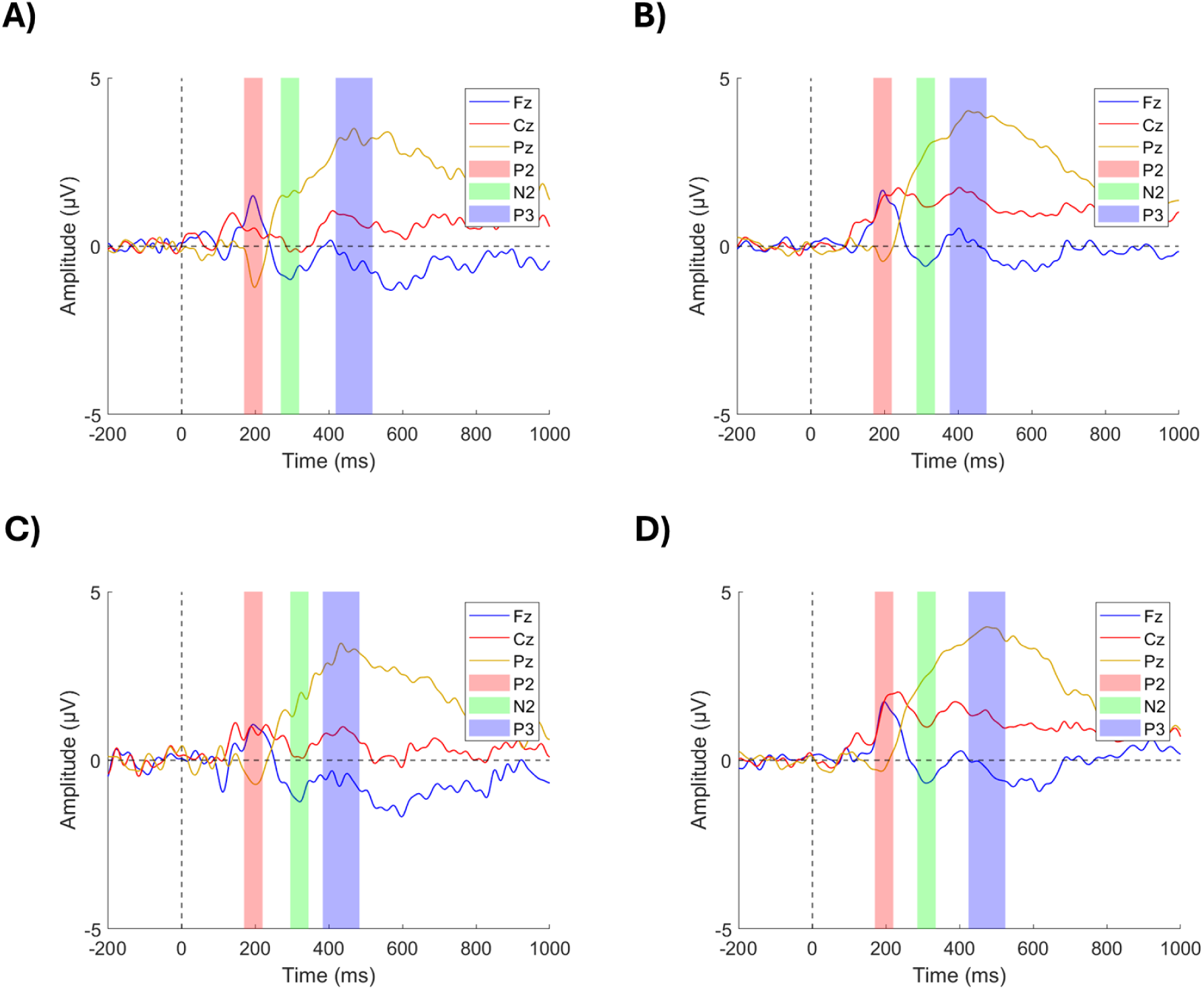
ERPs of the grand average of channels Fz, Cz and Pz used to identify the time windows for averaging during the cTBS-vertex condition of the visuospatial N-back. Channels Fz and Cz were used for the P200 and N200, and channels Cz and Pz were used for the P300. A) 2-back target trials. B) 2-back non-target trials. C) 3-back target trials. D) 3-back non-target trials.

**Figure 6.**
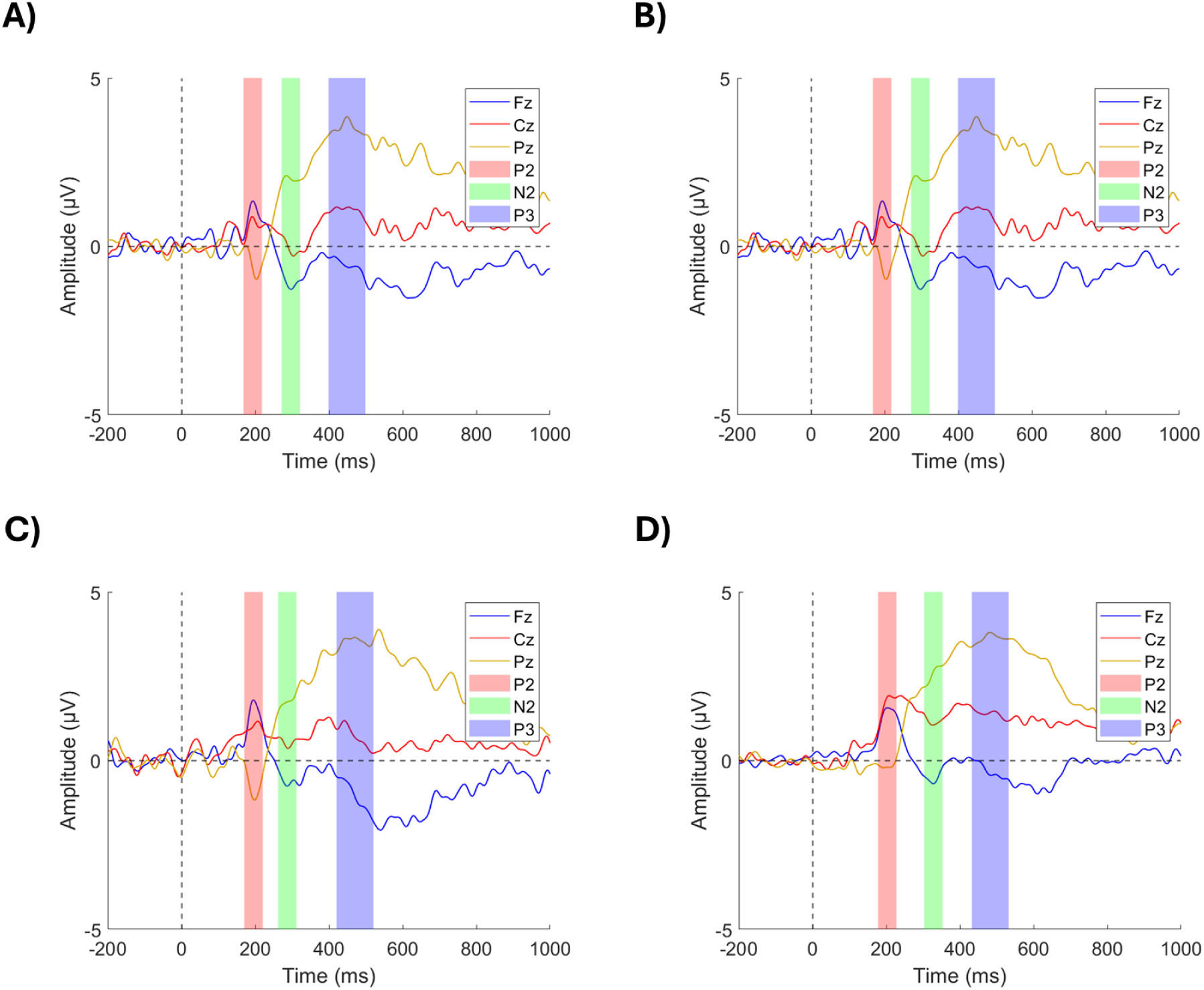
ERPs of the grand average of channels Fz, Cz and Pz used to identify the time windows for averaging during the cTBS-DLPFC condition of the visuospatial N-back. Channels Fz and Cz were used for the P200 and N200, and channels Cz and Pz were used for the P300. A) 2-back target trials. B) 2-back non-target trials. C) 3-back target trials. D) 3-back non-target trials.

Peak detection ranges were set as follows: P200 = 150–275ms (Bourisly & Shuaib, 2018), N200 = 200–350ms (Folstein & Van Petten, 2008), and P300 = 300–500ms (Johnson Jr., 1993; Polich, 2007; Scharinger et al., 2017). Mean amplitudes were extracted from ±25ms around the P200 and N200 peaks and ±50ms around the P300 peak (Ren et al., 2023). ERP latencies were defined as the peak time within each window. Electrodes analysed included Fz, F3, F4, Cz, C3, C4, Pz, P3, and P4.

### Statistical analysis

All statistical analyses were conducted in RStudio (2023). Bayesian mixed-effects models (BMEMs) were used to estimate stimulation effects, offering probabilistic inference and more nuanced interpretation, particularly with modest sample sizes (Dienes, 2011, 2014, 2016). Accuracy data were modelled using a 3 (stimulation condition) × 2 (cognitive load) design, and reaction times (RTs) with a 3 × 2 × 2 (stimulus type) design, including participant as a random effect.

Models were fitted using Stan via the *brms* package (Sorensen & Vasishth, 2016) using four chains of 4,000 iterations (1,000 iteration warmup period). A Student-t likelihood with three degrees of freedom and neutral priors centred at zero (scale = 0.5 for coefficients; scale = 1 for intercepts and variances) was adopted for robustness to outliers and heavy-tailed distributions (Jylanki et al., 2011). Convergence was confirmed by R^ < 1.01 and effective sample sizes > 1,000 for all parameters.

For ERP data, linear mixed-effects models (LMEMs) were run separately for each component (amplitude and latency) using a 3 (stimulation) × 2 (load) × 2 (stimulus type) × 8 (electrode) design, with participant as a random factor. Post hoc comparisons employed Tukey correction. Frequentist LMEMs were chosen to align with standard EEG analysis practices.

Finally, exploratory multiple regression models examined the relationship between ERP components and WM performance (accuracy and/or RTs), guided by BMEM and LMEM outcomes. Separate models were fitted for each N-back task, with the dependent variable defined as the change in performance between baseline and each stimulation condition. Stimulus type and cognitive load were excluded if non-significant or collinear; when retained, their interactions with ERP components were modelled. Optimal models were selected based on *p* ≤ 0.05, *Adj* R², and ANOVA comparisons of nested models. If no significant interactions emerged, simpler additive models were preferred following the principle of parsimony. ERP predictors were mean-centred and standardised to reduce multicollinearity. Analyses focused on frontal electrodes (Fz, F3, F4) for verbal WM and frontal and parietal electrodes (Pz, P3, P4) for visuospatial WM. This focus was theoretically motivated, given the established involvement of frontal regions, particularly the DLPFC, in WM processes, and aligned with the stimulation site (left DLPFC) employed in the study.

## RESULTS

### Verbal N-back – Accuracy

The Bayesian analysis provided moderate evidence for an increase in accuracy in the cTBS-vertex condition compared to baseline (B = 0.41, SE = 0.16, 95% CI = [0.10, 0.72], PP = 90%) suggesting learning effects. In contrast, for the cTBS-DLPFC condition, the model showed inconclusive evidence (B = 0.26, SE = 0.15, 95% CI = [-0.04, 0.55], PP = 57%), indicating that the data were consistent with both the presence and absence of learning effects (see Figure 7A).

**Figure 7.**
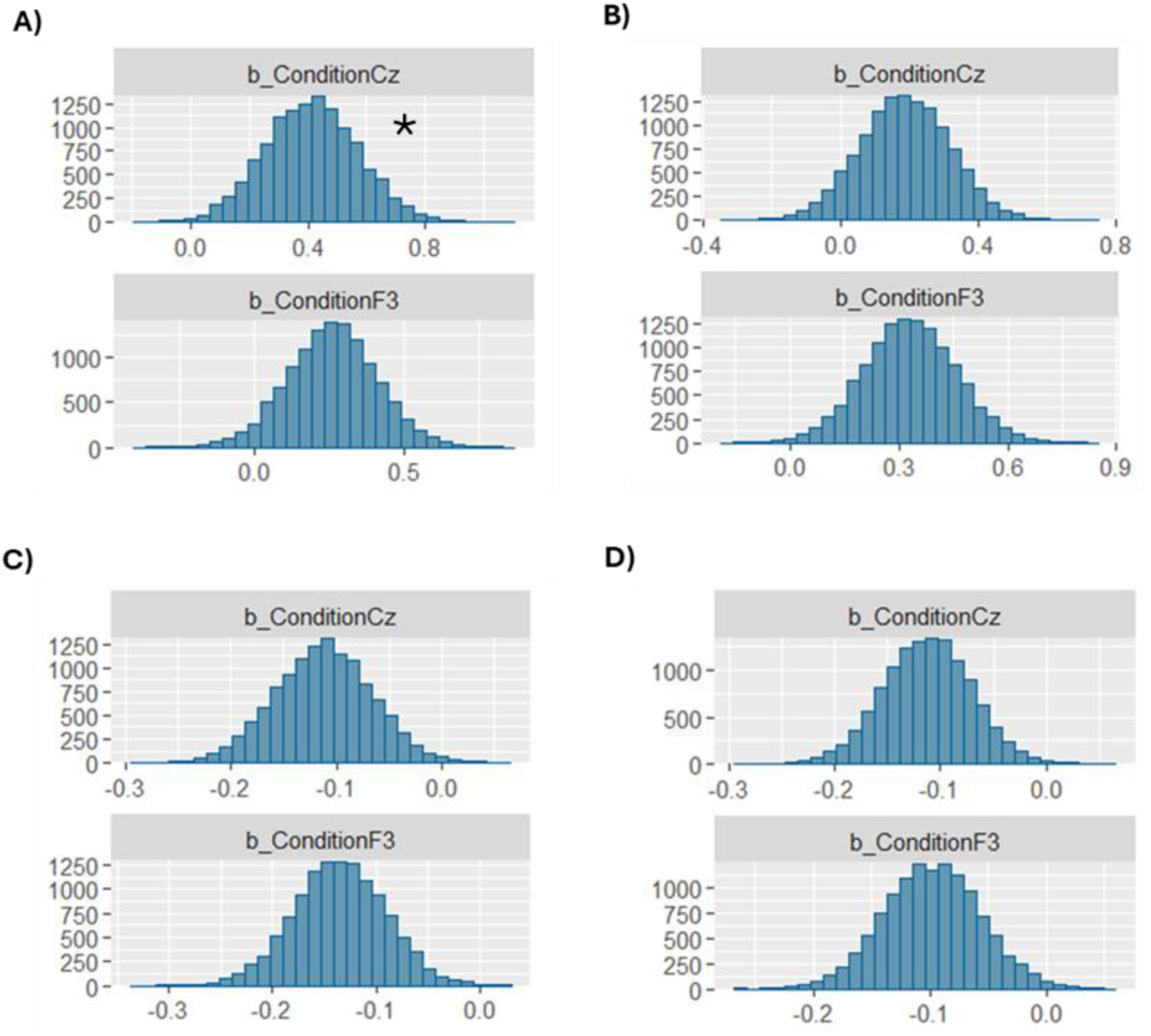
Posterior distributions of regression coefficients (β) from BMEMs assessing stimulation effects on behavioural performance. Cz refers to the cTBS-vertex condition and F3 to the cTBS-DLPFC condition (10–10 electrode system). (A) Verbal N-back accuracy. (B) Visuospatial N-back accuracy. (C) Verbal N-back RTs. (D) Visuospatial N-back RTs. Only the verbal N-back accuracy distribution under cTBS-vertex excluded zero within its 95% credible interval, indicating credible learning effects (marked with a *); all others overlapped with zero, showing no robust stimulation effects.

Regarding the differences between the cTBS-vertex and the cTBS-DLPFC condition, evidence was also inconclusive (B = 0.15, SE = 0.17, 95% CI = [-0.18, 0.48], PP = 22%), indicating that the data do not support a reliable difference between these two stimulation conditions. Finally, no evidence for effects of interactions were noted.

### Verbal N-back – RTs

The Bayesian analysis provided inconclusive evidence of learning effects in RTs, relative to baseline, for both the cTBS-vertex condition (B = -0.11, SE = 0.05, 95% CI = [-0.20, -0.02], PP = 61%) and the cTBS-DLPFC condition (B = -0.14, SE = 0.04, 95% CI = [-0.22, - 0.05], PP = 87%), indicating that the data were consistent with both the presence and absence of learning effects (see Figure 7C).

Regarding the differences between the cTBS-vertex and the cTBS-DLPFC condition, evidence was strong against any differences between the two stimulation conditions (B = 0.02, SE = 0.05, 95% CI = [-0.07, 0.12], PP = 6%), suggesting that the data strongly support a null difference between these two stimulation conditions. Finally, no evidence for effects of interactions were noted.

### Visuospatial N-back – Accuracy

The Bayesian analysis provided inconclusive evidence of learning effects in accuracy, relative to baseline, for both the cTBS-vertex condition (B = 0.19, SE = 0.13, 95% CI = [-0.07, 0.44], PP = 38%) and the cTBS-DLPFC condition (B = 0.33, SE = 0.13, 95% CI = [0.08, 0.58], PP = 86%), indicating that the data were consistent with both the presence and absence of learning effects (see Figure 7B).

Regarding the differences between the cTBS-vertex and the cTBS-DLPFC condition, evidence was also inconclusive (B = -0.14, SE = 0.13, 95% CI = [-0.04, 0.11], PP = 22%), suggesting that the data do not support a reliable difference between these two stimulation conditions. Finally, no evidence for effects of interactions were noted.

### Visuospatial N-back – RTs

The Bayesian analysis provided inconclusive evidence of learning effects in RTs, relative to baseline, for both the cTBS-vertex condition (B = -0.11, SE = 0.04, 95% CI = [-0.20, -0.03], PP = 73%) and the cTBS-DLPFC condition (B = -0.10, SE = 0.04, 95% CI = [-0.18, - 0.02], PP = 53%), indicating that the data were consistent with both the presence and absence of learning effects (see Figure 7D).

Regarding the differences between the cTBS-vertex and the cTBS-DLPFC condition, evidence was strong against any differences between the two stimulation conditions (B = -0.01, SE = 0.04, 95% CI = [-0.01, 0.07], PP = 5%), suggesting that the data strongly support a null difference between these two stimulation conditions. Finally, no evidence for effects of interactions were noted. Table 1 summarises the findings of the full Bayesian analysis and Figure 8 show the distribution of accuracy scores (d’ values) and RTs.

**Figure 8.**
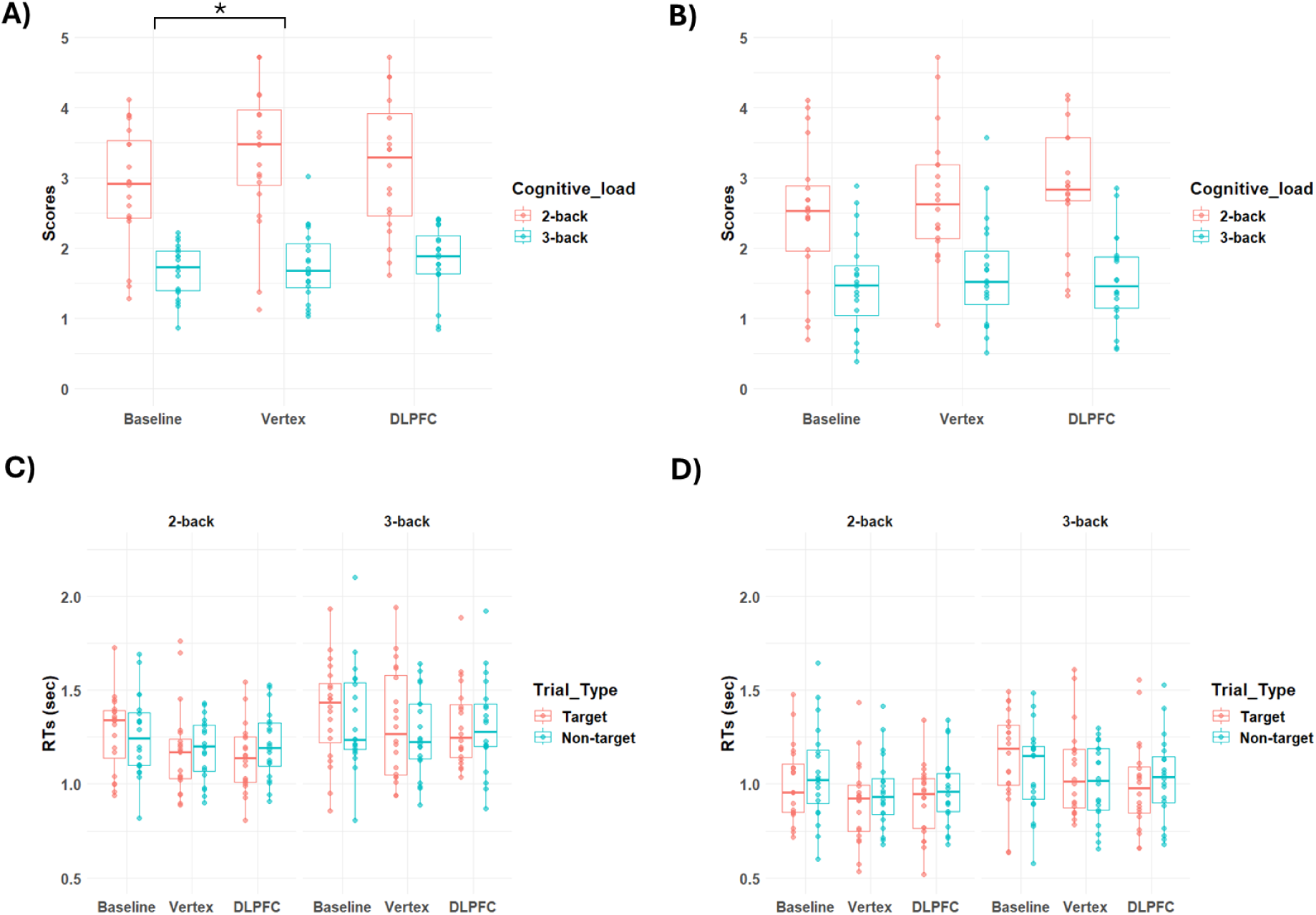
Behavioural results: Accuracy scores (d’ values) and RTs distributions. A) Verbal N-back accuracy. B) Visuospatial N-back accuracy. C) Verbal N-back RTs. D) Visuospatial N-back RTs. Moderate evidence supporting learning effects in accuracy scores was found in the cTBS-vertex condition of the verbal N-back (marked with a *).

**Table 1.** Bayesian analysis summary of findings. Moderate evidence supporting learning effects in accuracy scores was found in the cTBS-vertex condition of the verbal N-back. Evidence was inconclusive in all the other cases.

|  | Verbal N-back |  | Visuospatial N-back |  |
| --- | --- | --- | --- | --- |
|  | Vertex | DLPFC | Vertex | DLPFC |
| Accuracy | <u>Moderate</u> | Inconclusive | Inconclusive | Inconclusive |
| RTs | Inconclusive | Inconclusive | Inconclusive | Inconclusive |

### ERP amplitudes

Across both verbal and visuospatial N-back tasks, no significant main effects or interactions involving stimulation condition were found for any ERP component, indicating no amplitude differences across stimulation groups. A marginal trend for the visuospatial P200 (*F*(1,2033) = 2.95, *p* = .05, *η2* = 0.003) did not survive post hoc testing. Amplitude distributions are shown in Figures 9 and 10, and full statistics are provided in Tables S1 and S2 of the Supplementary Materials.

**Figure 9.**
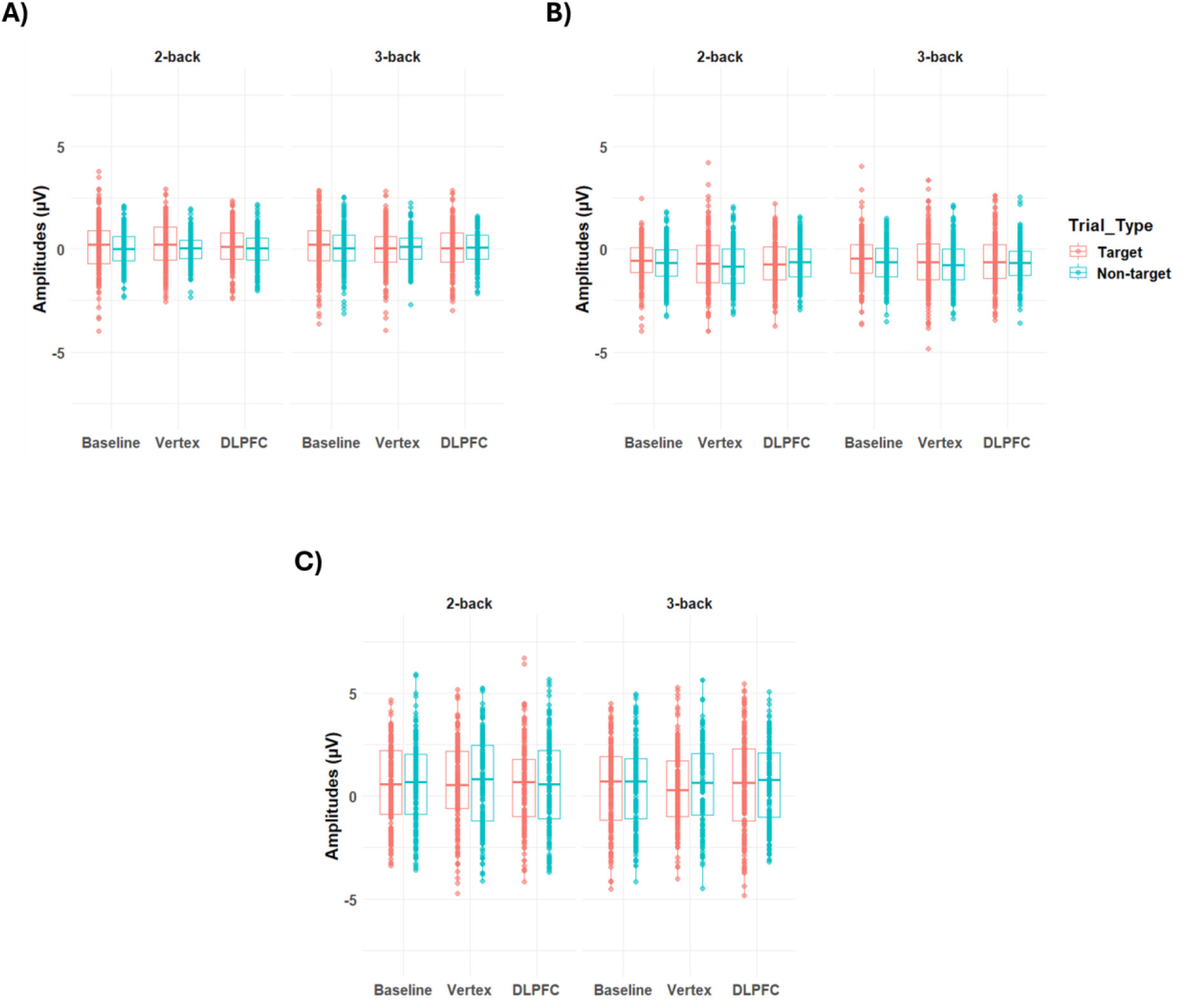
ERP amplitude distributions in the verbal N-back across baseline, cTBS-vertex, and cTBS-DLPFC conditions. A) P200 amplitudes. B) N200 amplitudes. C) P300 amplitudes. No significant differences were observed between stimulation conditions in any of the ERP components.

**Figure 10.**
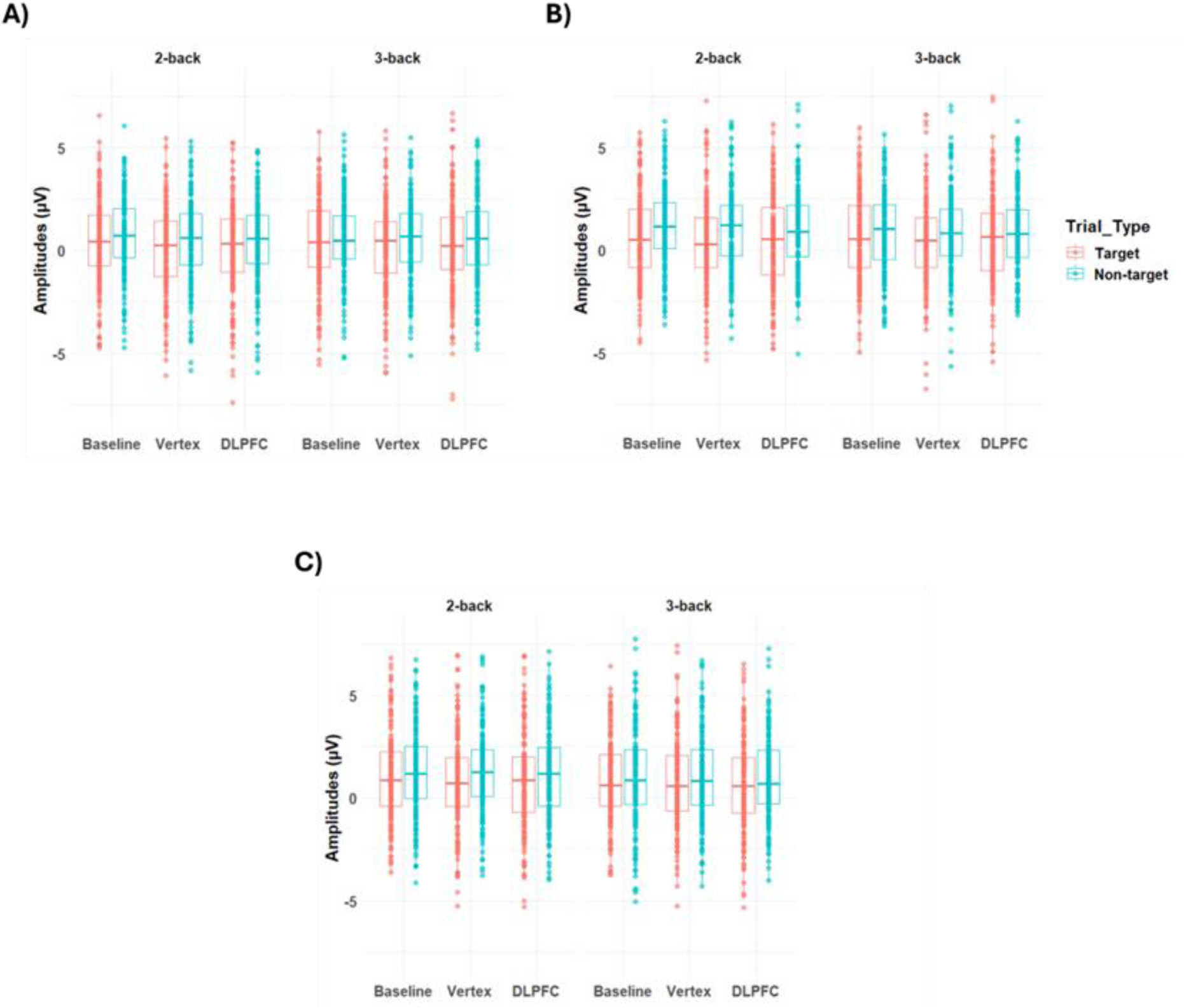
ERP amplitude distributions in the visuospatial N-back across baseline, cTBS-vertex, and cTBS-DLPFC conditions. A) P200 amplitudes. B) N200 amplitudes. C) P300 amplitudes. No significant differences were observed between stimulation conditions in any of the ERP components.

### ERP latencies

Several latency effects emerged in the verbal N-back. cTBS to the DLPFC generally prolonged P200 latencies in target trials across cognitive loads compared to both baseline and the cTBS-vertex condition (all ps < .001; *d* = 0.57 to 2.64), whereas cTBS-vertex shortened latencies relative to baseline (2-back: *t*(2033) = -7.32, *p* < .001, *d* = -0.78; 3-back: *t*(2033) = - 20.89, *p* < .001, *d* = -2.13). A similar pattern was observed for P300 latencies, which were longer under DLPFC stimulation particularly in target trials of higher load (3-back; vs baseline: *t*(2033) = 4.32, *p* < .001, *d* = 0.43; vs cTBS-vertex: *t*(2033) = 5.20, *p* < .001, *d* = 0.46), but in this case both stimulation conditions produced longer latencies than baseline overall (all ps < .01; *d* = 0.41 to 0.43). In contrast, N200 latencies showed the opposite trend, with slightly shorter responses following DLPFC stimulation at lower load (2-back; vs cTBS-vertex: *t*(2033) = -3.62, *p* < .01, *d* = -0.27) and in non-target trials (vs baseline: *t*(2033) = -3.47, *p* < .01, *d* = - 0.25; vs cTBS-vertex: *t*(2033) = -5.21, *p* < .001, *d* = -0.40) (see Figure 11). Overall, these findings suggest that DLPFC stimulation slowed neural processing during P200 and P300, particularly in target trials, but speeded processing during N200 at lower load and in non-target trials.

**Figure 11.**
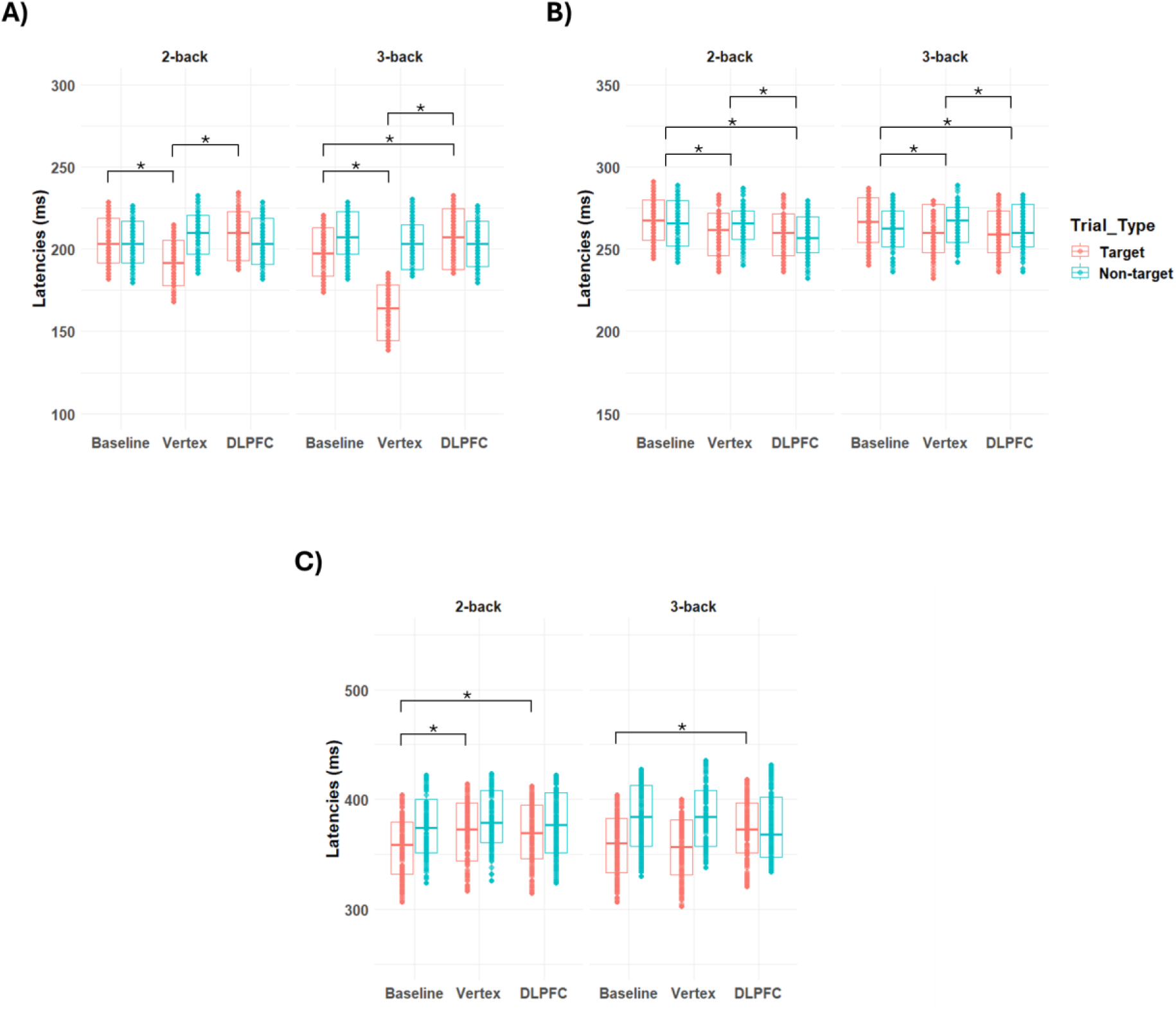
ERP latency distributions in the verbal N-back across baseline, cTBS-vertex, and cTBS-DLPFC conditions for target and non-target trials. A) P200 latency. B) N200 latency. C) P300 latency. Significant reductions in latency were observed after stimulation, with patterns varying by component, load, and stimulation site (see text for statistics).

In the visuospatial N-back, cTBS to the DLPFC also delayed P200 latencies, especially under higher cognitive load (vs cTBS-vertex; target: *t*(2033) = 3.53, *p* < .05, *d* = 0.37; non-target: (*t*(2033) = 3.64, *p* < .05, *d* = 0.39). However, across both stimulation sites, P200 latencies were generally shorter than at baseline across cognitive loads and stimulus types (all ps < .001; *d* = -2.34 to -0.79). DLPFC stimulation also prolonged P300 latencies, particularly in non-target trials across both cognitive loads (vs cTBS-vertex; 2-back: *t*(2033) = 10.33, *p* < .001, *d* = 1.02; 3-back: *t*(2033) = 5.52, *p* < .001, *d* = 0.55), while both stimulation conditions produced longer latencies than baseline overall (all ps < .001; *d* = 0.58 to 1.09). For the N200, effects were mixed: DLPFC stimulation delayed latencies in 2-back target trials (vs cTBS-vertex: *t*(2033) = 4.69, *p* < .001, *d* = 0.47) and in 3-back non-target trials (vs cTBS-vertex: *t*(2033) = 9.44, *p* < .001, *d* = 0.97), but shortened latencies in 3-back target trials (vs cTBS-vertex: *t*(2033) = -16.73, *p* < .001, *d* = -1.77). Notably, both stimulation conditions increased latencies relative to baseline (ps < .001; *d* = 0.38 to 1.39) (see Figure 12). Together, these findings indicate that DLPFC stimulation slowed neural processing during P200, particularly under high cognitive load, and prolonged P300 latency during non-target processing. In contrast, N200 latency effects were load- and stimulus-dependent, with slowing at low-load target and high-load non-target trials but speeding at high-load target trials.

**Figure 12.**
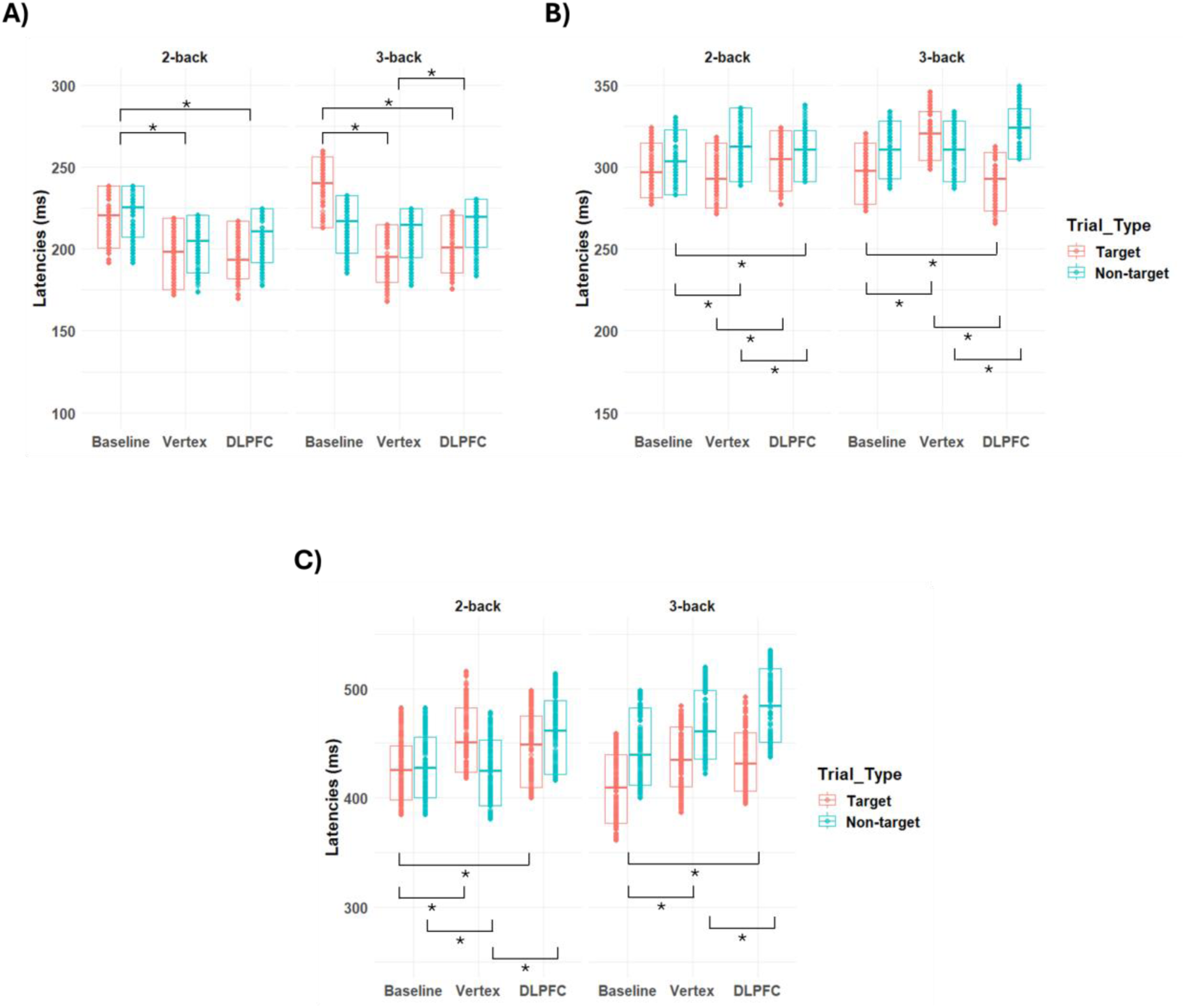
ERP latency distributions in the visuospatial N-back across baseline, cTBS-vertex, and cTBS-DLPFC conditions for target and non-target trials. A) P200 latency. B) N200 latency. C) P300 latency. Significant reductions in latency were observed after stimulation, with patterns varying by component, load, and stimulation site (see text for statistics).

Tables 2 and 3 summarise the main findings for the verbal and visuospatial N-back ERP latency analyses, respectively. Full statistics are provided in Tables S3 and S4 of the Supplementary Materials.

**Table 2.** Significant differences in ERP latencies between stimulation conditions in the verbal N-back. Only shorter latencies are shown; blank cells indicate no significant differences. In general, cTBS affected latency patterns differently across components, with effects modulated by cognitive load and stimulus type (see text for details).

| ERP | Cognitive Load | Trial Type | Vertex vs Baseline | DLPFC vs Baseline | Vertex vs DLPFC |
| --- | --- | --- | --- | --- | --- |
| P200 | 2-back | Target | **Vertex** | - | Vertex |
|  | 3-back | Target | **Vertex** | Baseline | Vertex |
| N200 | 2-back | - | **Vertex** | **DLPFC** | DLPFC |
|  | - | Target | **Vertex** | **DLPFC** | - |
|  | - | Non-target | - | **DLPFC** | DLPFC |
| P300 | 2-back | Target | Baseline | Baseline | - |
|  | 3-back | Target | - | Baseline | Vertex |

**Table 3.**
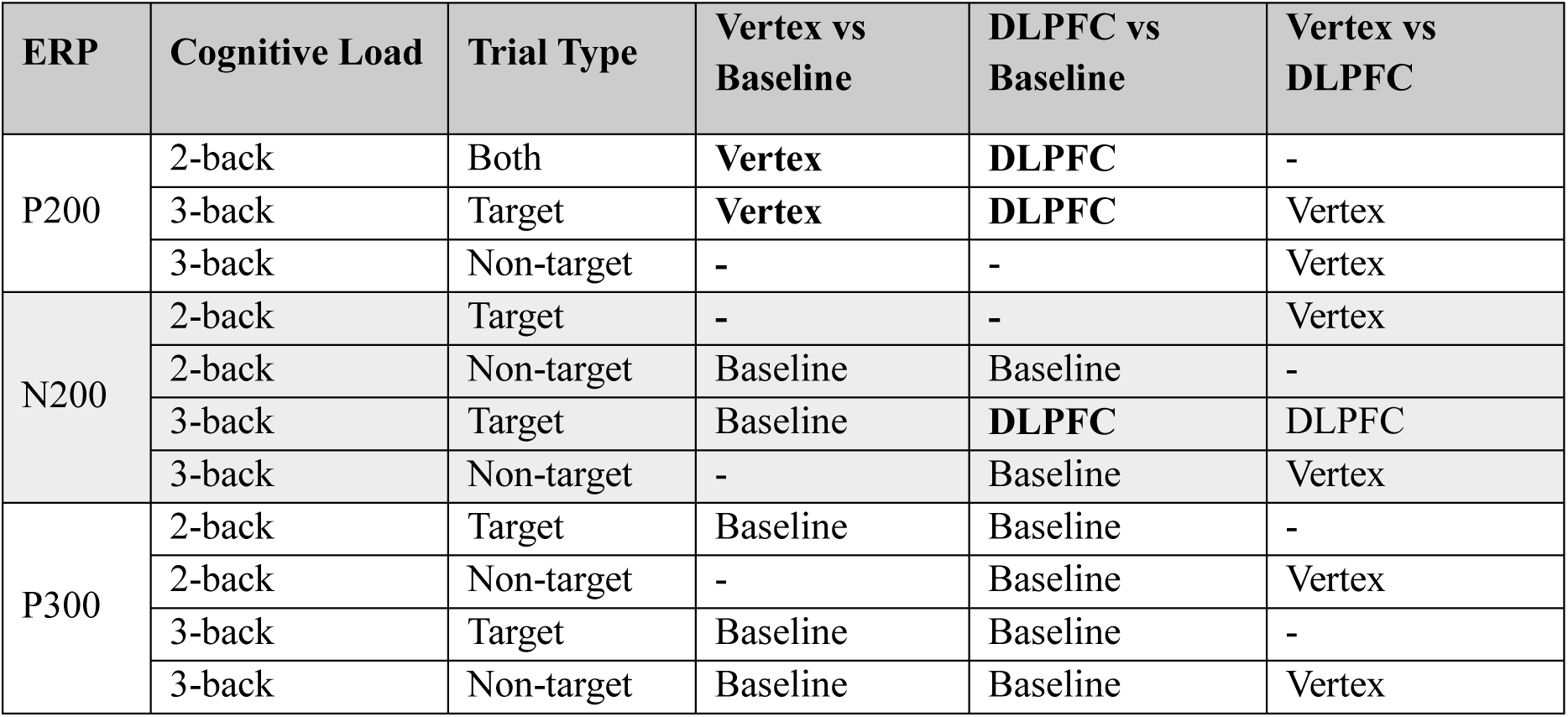
Significant differences in ERP latencies between stimulation conditions in the visuospatial N-back. Only shorter latencies are shown; blank cells indicate no significant differences. Overall, cTBS modulated ERP latencies across components, with distinct patterns for each and effects varying by cognitive load and stimulus type (see text for details).

| ERP | Cognitive Load | Trial Type | Vertex vs Baseline | DLPFC vs Baseline | Vertex vs DLPFC |
| --- | --- | --- | --- | --- | --- |
| P200 | 2-back | Both | **Vertex** | **DLPFC** | - |
|  | 3-back | Target | **Vertex** | **DLPFC** | Vertex |
|  | 3-back | Non-target | - | - | Vertex |
| N200 | 2-back | Target | - | - | Vertex |
|  | 2-back | Non-target | Baseline | Baseline | - |
|  | 3-back | Target | Baseline | **DLPFC** | DLPFC |
|  | 3-back | Non-target | - | Baseline | Vertex |
| P300 | 2-back | Target | Baseline | Baseline | - |
|  | 2-back | Non-target | - | Baseline | Vertex |
|  | 3-back | Target | Baseline | Baseline | - |
|  | 3-back | Non-target | Baseline | Baseline | Vertex |

### Regression analysis between behavioural results and ERP latencies

Given that latency effects appeared more sensitive to stimulation than amplitudes, regression analyses were conducted to examine whether ERP latencies predicted accuracy scores.

### Verbal N-back – cTBS-Vertex Model

The best model included frontal ERP latency changes (P200, N200, P300) without interaction terms. N200 latency negatively predicted performance (*t*(36) = -2.32, *p* < .05, *η2* = 0.11), suggesting that shorter N200 latencies relative to baseline were associated with better accuracy in the cTBS-vertex condition (see Figure 13). The model explained 12% of the variance (*F*(3,36) = 2.77, *p* = .06, *Adj* R² = 0.12). A model including latency interaction terms improved variance explained (*F*(7,32) = 2.46, *p* < .05, *Adj* R² = 0.21) but did not significantly improve fit over the simpler model (*F*(4,32) = 2.00, *p* = .12), which was therefore retained.

**Figure 13.**
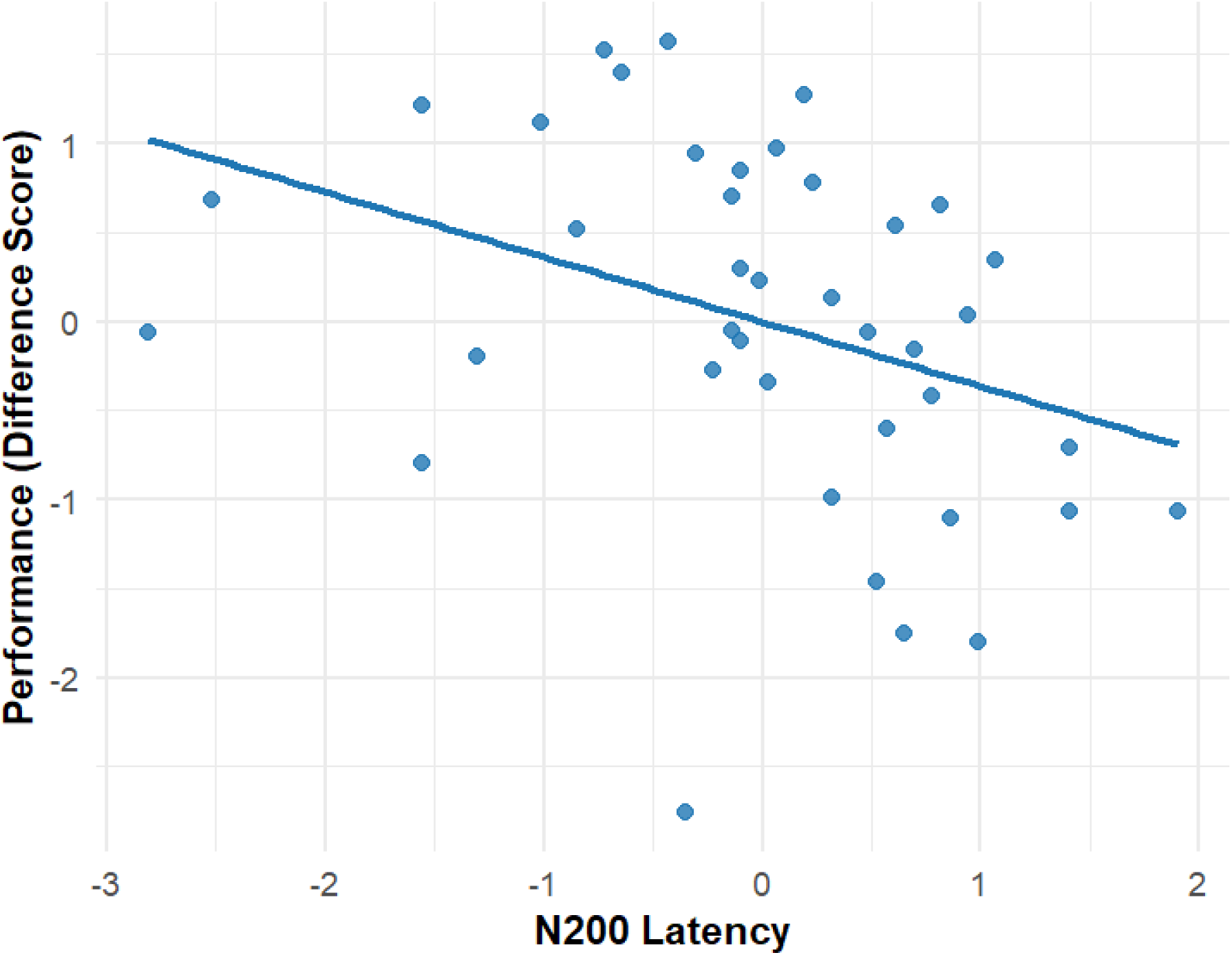
N200 latency main effect in the cTBS-vertex accuracy model of the verbal N-back. Shorter N200 latencies relative to baseline were associated with an increase in performance.

### Verbal N-back – cTBS-DLPFC Model

The best model included frontal ERP latency changes (P200, N200, P300) and cognitive load as predictors. P200 latency negatively predicted performance (*t*(32) = -3.41, *p* < .01, *η2* = 0.14), and significant interactions were found between cognitive load and both P200 latency (*t*(32) = 2.96, *p* < .01, *η2* = 0.15) and N200 latency (*t*(2033) = -2.48, *p* < .05, *η2* = 0.14). Simple slopes showed shorter P200 latencies predicted better performance in the 2-back (B = -0.76, 95% CI [-1.21, -0.30], *p* < .01), but not in the 3-back (B = 0.24, 95% CI [-0.28, 0.75], *p* = .36) (see Figure 14). Although the interaction with N200 latency was significant, neither slope reached significance (*ps* > .07). (see Figure 15). These results suggest that shorter P200 latencies relative to baseline were associated with better accuracy in the cTBS-DLPFC condition only under lower cognitive load. The model explained 27% of the variance (*F*(7,32) = 3.04, *p* = .01, *Adj* R² = 0.27).

**Figure 14.**
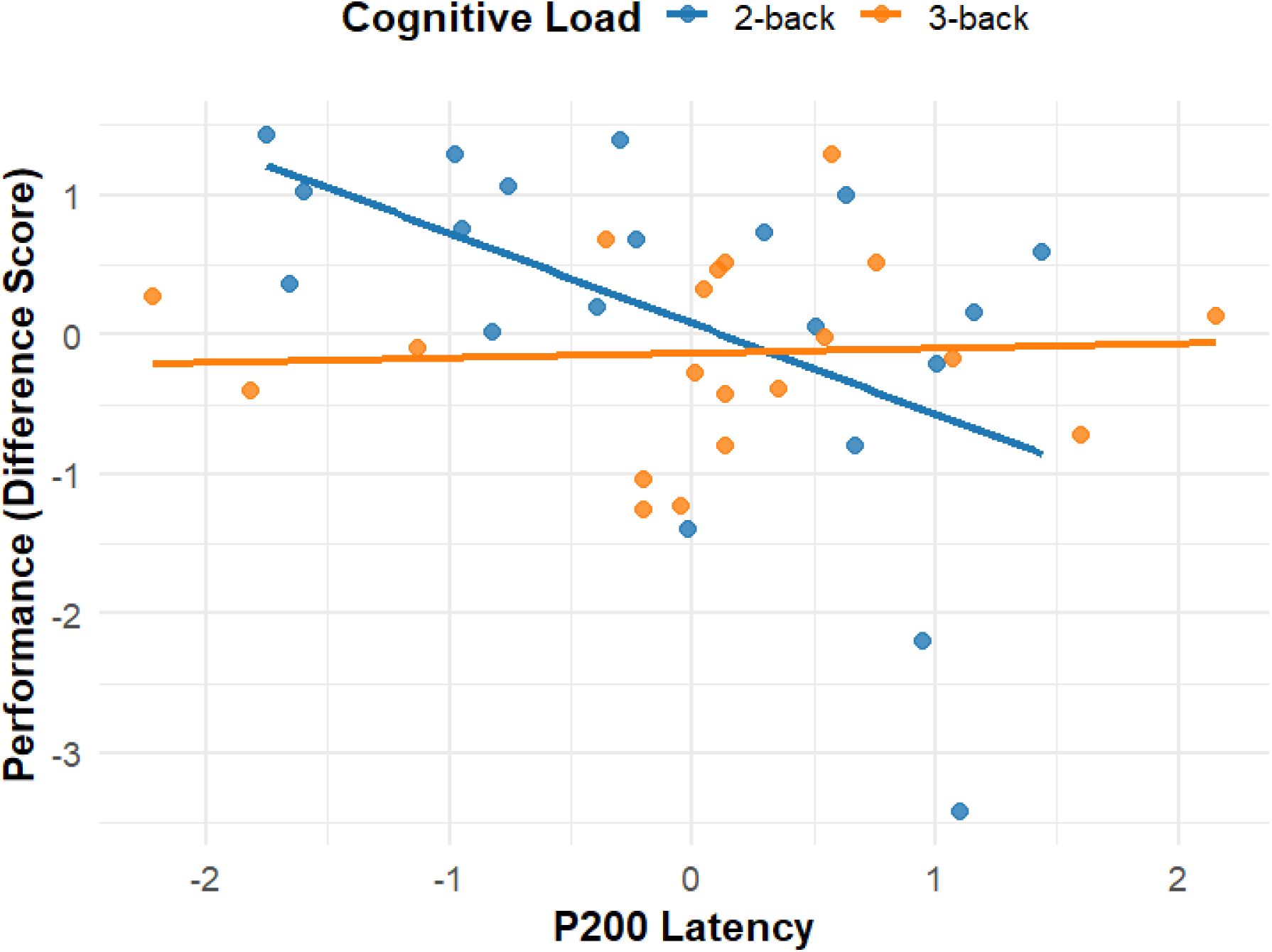
P200 latency interaction effects in the cTBS-DLPFC accuracy model of the verbal N-back. Shorter P200 latencies relative to baseline were associated with an increase in performance only in the 2-back.

**Figure 15.**
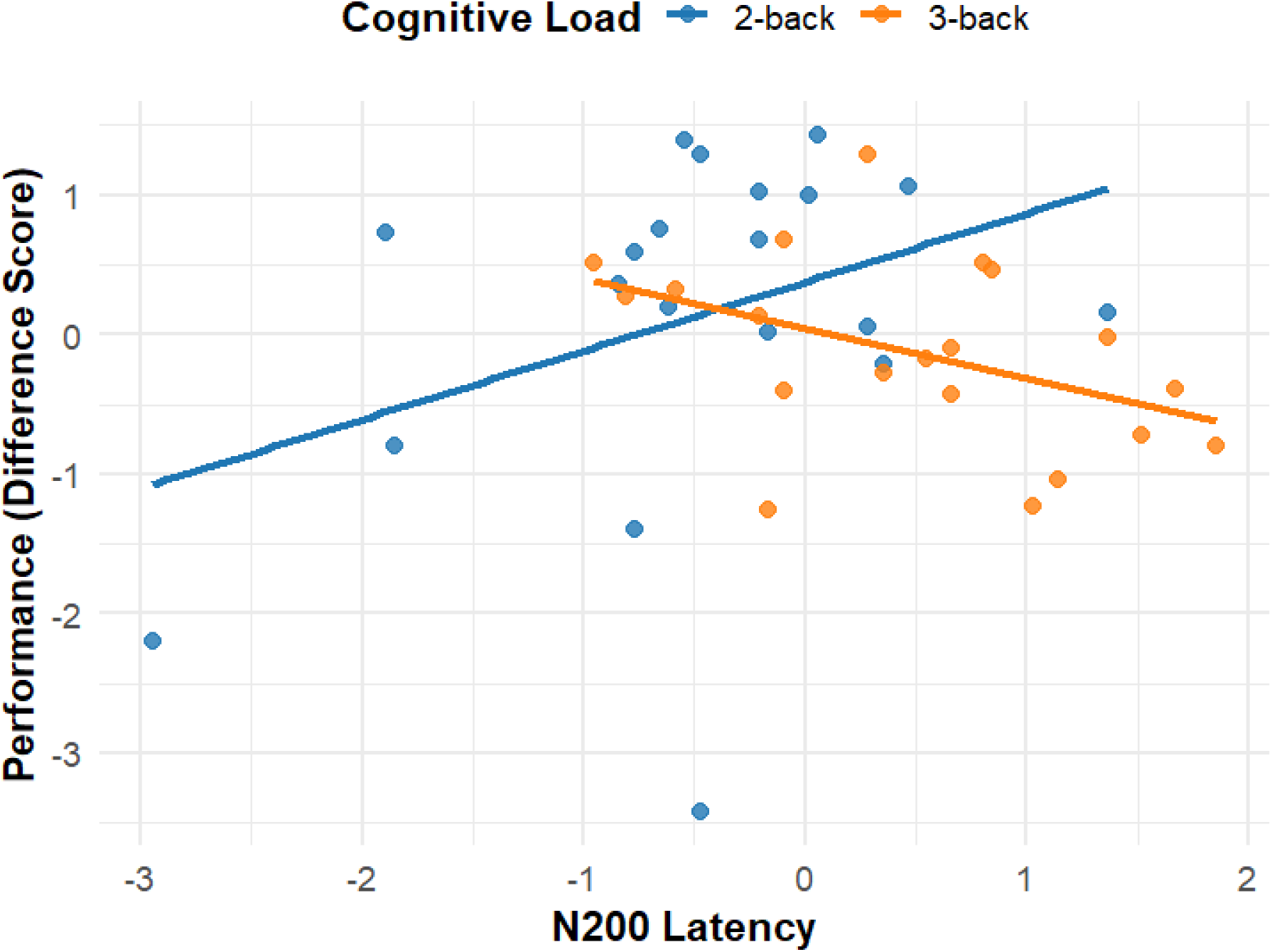
N200 latency interaction effects in the cTBS-DLPFC accuracy model of the verbal N-back. The effect of N200 latency on performance was load-specific, showing a positive correlation in the 2-back and a negative correlation in the 3-back, although neither effect reached statistical significance.

### Visuospatial N-back – cTBS-Vertex Model

The cTBS-vertex model did not perform well regardless of the predictors chosen (all *ps* > .05 and all *Adj* R < 0.07).

### Visuospatial N-back – cTBS-DLPFC Model

The best cTBS-DLPFC model included frontal P200 and N200, parietal P200, N200 and P300 parietal latency changes, and cognitive load and predictors. N200 latency negatively predicted performance (*t*(32) = -3.32, *p* < .01, *η2* = 0.21), whereas P300 latency was positively associated with performance (*t*(32) = 2.49, *p* < .01, *η2* = 0.21). A significant interaction was found between cognitive load and parietal P300 latency (*t*(32) = -3.18, *p* < .01, *η2* = 0.21). Simple slopes showed that longer P300 latencies predicted better performance in the 2-back (95% CI [0.16, 1.05], *p* < .01), but not in the 3-back (95% CI [-0.77, 0.07], *p* = .10). These results suggest that shorter fronto-parietal N200 and longer parietal P300 latencies relative to baseline were associated with better accuracy in the cTBS-DLPFC condition, with the P300 effect specific to lower cognitive load (see Figures 16 and 17). The model explained 33% of the variance (*F*(7,32) = 3.69, *p* < .01, *Adj* R² = 0.33).

**Figure 16.**
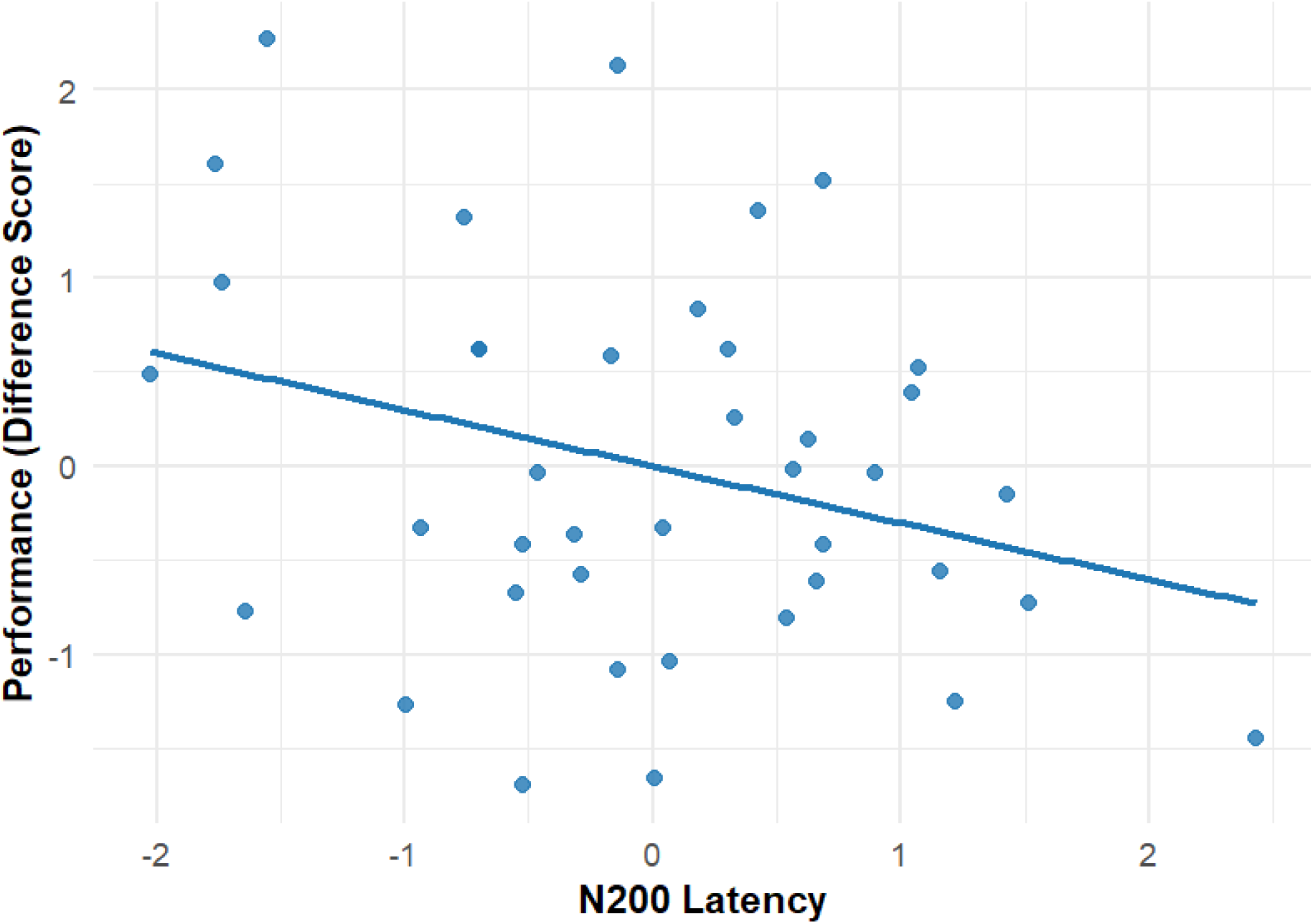
N200 latency main effect in the cTBS-DLPFC accuracy model of the visuospatial N-back. Shorter N200 latencies relative to baseline were associated with an increase in performance.

**Figure 17.**
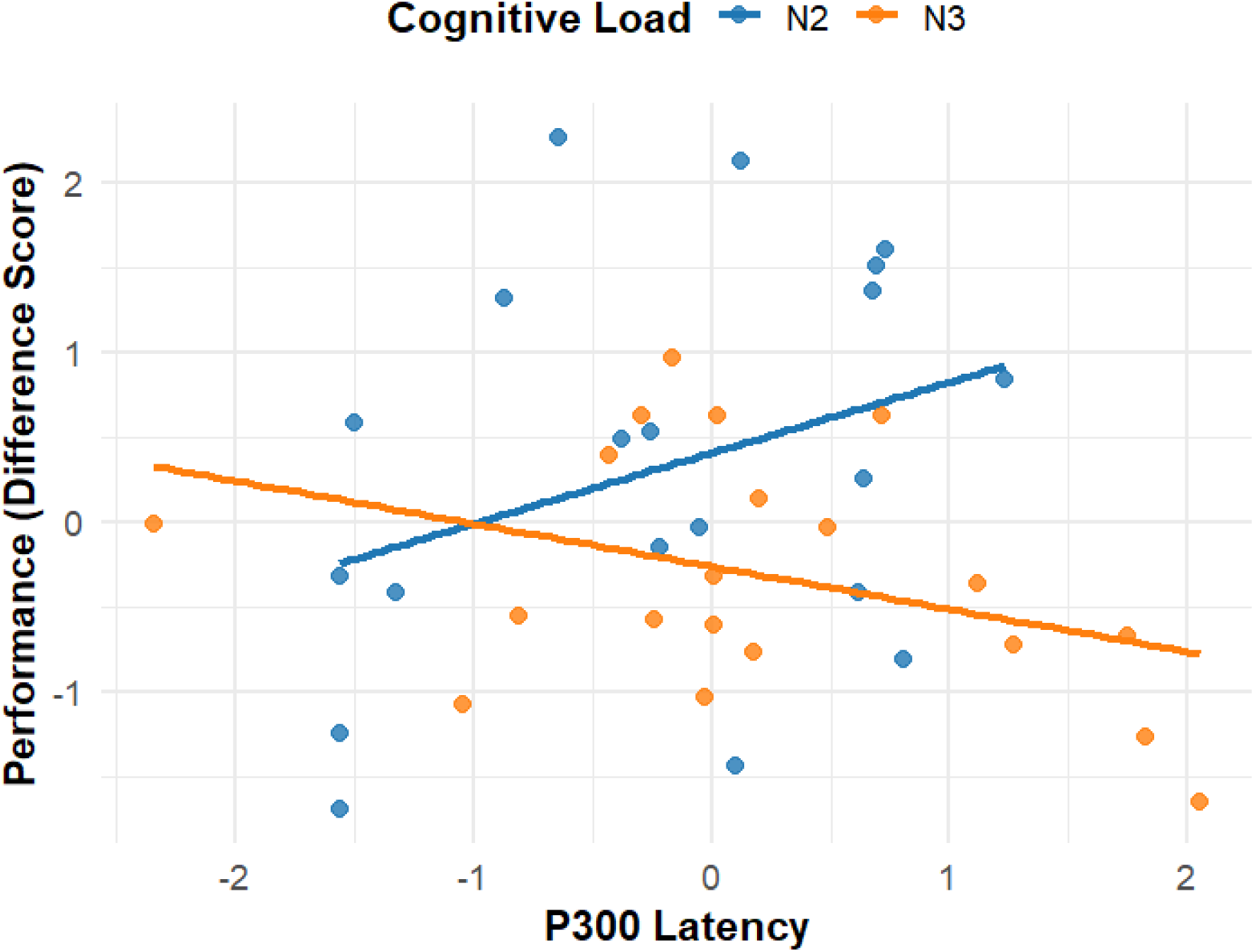
P300 latency interaction effects in the cTBS-DLPFC accuracy model of the visuospatial N-back. Longer P300 latencies relative to baseline were associated with an increase in performance only in the 2-back.

Table 4 summarises the main findings of the regression analysis. A summary of all models and full statistics is provided in Tables S5–S11 of the Supplementary Materials.

**Table 4.** Summary of correlations between ERP latencies and accuracy. A negative correlation direction reflects greater accuracy improvements with shorter latencies; and a positive correlation direction reflects the opposite pattern.

| Task Type | ERP | Stimulation Condition | Correlation Direction | Cognitive Load | Channels |
| --- | --- | --- | --- | --- | --- |
| Verbal N-back | P200 | DLPFC | Negative | 2-back | Frontal |
|  | N200 | Vertex | Negative | Both | Frontal |
| Visuospatial N-back | N200 | DLPFC | Negative | Both | Frontal / Parietal |
|  | P300 | DLPFC | Positive | 2-back | Parietal |

## DISCUSSION

We hypothesised that cTBS to the DLPFC would disrupt learning in the verbal N-back, particularly in the 2-back, while learning would be preserved following vertex stimulation. For the visuospatial N-back, we expected learning in both stimulation conditions, irrespective of load.

Bayesian analyses partially supported these predictions. In the verbal task, moderate evidence for learning emerged following cTBS to the vertex, whereas evidence remained inconclusive for DLPFC stimulation. Although the results do not conclusively demonstrate that DLPFC stimulation abolished learning, the absence of robust improvement (which was evident in the cTBS-vertex condition) suggests reduced or more variable performance. Contrary to expectations, this pattern was consistent across both cognitive loads rather than being restricted to the 2-back, echoing previous findings that DLPFC inhibition disrupts verbal WM learning under both moderate and high load (2-back and 3-back) (Ngetich et al., 2021; Schicktanz et al., 2015; Viejo-Sobera et al., 2017; Vékony et al., 2018).

In contrast, the visuospatial N-back yielded inconclusive evidence for learning regardless of stimulation site. This may reflect greater task difficulty and increased interindividual variability, consistent with research showing that visuospatial WM declines more steeply with age due to impairments in spatial and executive processes (Borella et al., 2014; Brown, 2016; Cansino et al., 2013; Holmes et al., 2019; Myerson et al., 1999; Salthouse, 2010). If participants struggled to improve across sessions, this may have masked stimulation-related effects. An alternative explanation is that stimulation may have induced inhibition across both sites; however, this is unlikely given the preserved learning following vertex stimulation in the verbal task and the domain-specific role of the left DLPFC. Evidence for learning in RTs was generally inconclusive.

ERP results help explain these behavioural patterns by revealing how DLPFC inhibition affected the temporal dynamics of cognitive processing. In the verbal N-back, vertex stimulation preserved learning, and shorter N200 latencies predicted higher accuracy, consistent with the N200’s involvement in conflict monitoring and inhibitory control (Daffner et al., 2011; Folstein & Van Petten, 2008; Gajewski & Falkenstein, 2014; Patel & Azzam, 2005; Xiao et al., 2019). Conversely, cTBS to the DLPFC produced even shorter N200 latencies—particularly in non-target trials of the 2-back—without corresponding accuracy gains. Such premature processing likely reflects dysregulated or inefficient control, consistent with reports of impaired cognitive control following DLPFC inhibition (Fried et al., 2014; Postle et al., 2006; Rogasch et al., 2015; Webler et al., 2022). The fact that this pattern was strongest during the more frequent non-target trials (70%) further supports the interpretation of increased reliance on habitual responding (Brevet-Aeby et al., 2016; Loftus et al., 2015; Yang et al., 2018).

Regression analyses revealed possible load-dependent compensatory responses following DLPFC disruption: in the 2-back, longer N200 latencies tended to predict better performance, whereas in the 3-back, shorter latencies showed a similar tendency. Although neither association reached significance, this pattern may reflect attempts by participants to adapt their strategies, slowing responses when feasible to enable more deliberate processing (Festini et al., 2018; Reuter-Lorenz & Cappell, 2008; Tucker & Stern, 2011). Such compensatory slowing is commonly reported in ageing, where older adults deliberately decelerate processing to preserve accuracy under higher cognitive demands, although not always successfully (Bugg et al., 2007; Wascher et al., 2012; Wascher & Beste, 2010).

This interpretation is supported by latency effects in both early and late ERP components. DLPFC stimulation delayed P200 and P300 latencies—especially in target trials—whereas vertex stimulation led to latency shortening. These differences suggest disrupted neural efficiency under DLPFC inhibition and enhanced learning-related processing in the vertex condition (Bourisly & Shuaib, 2018; Lijffijt et al., 2009). The frontal P200, associated with early attentional gating and the inhibition of irrelevant information (Blanchet et al., 2007; Chen & Mitra, 2009; Lijffijt et al., 2009; Luck & Hillyard, 1994), was the only component that significantly predicted performance: shorter P200 latencies relative to baseline were linked to higher accuracy. This finding suggests that DLPFC inhibition impaired top-down modulation of sensory–attentional processes, delaying stimulus evaluation and constraining learning. Similar P200 delays have been identified as early neurophysiological markers of cognitive decline in ageing (Gu & Zhang, 2017; López Zunini et al., 2016; Oakley et al., 2025), underscoring the role of DLPFC function in early attentional control.

### Why latencies and not amplitudes?

The observation that stimulation influenced ERP latencies but not amplitudes suggests that cTBS primarily disrupted processing speed and neural efficiency rather than the magnitude of cognitive resource allocation. This interpretation is supported by two arguments. First, older adults frequently show reduced or absent DLPFC activation during WM tasks compared with younger adults (Mattay et al., 2006; Yaple et al., 2019; Zając-Lamparska et al., 2024). Second, they often compensate by engaging broader, more distributed neural networks (Cabeza et al., 2018; Festini et al., 2018; Jannati et al., 2023; Korsch et al., 2014; Kropotov et al., 2016; Morcom & Johnson, 2015). Although the DLPFC continues to play an important role, its reduced baseline activation in older adults may make it less directly susceptible to stimulation. Instead, cTBS may have disrupted functional connectivity and coordination between the DLPFC and other regions, affecting the timing of cognitive operations without substantially changing response magnitude.

This interpretation aligns with the processing-speed hypothesis, which posits that age-related limitations in processing speed hinder cognitive function by altering the timing of information transfer (Kropotov et al., 2016; Salthouse, 1996). Older adults often struggle with executive components of WM—such as cognitive control and updating—and compensate by relying more heavily on other systems such as short-term and long-term memory (Gajewski et al., 2018). This also fits with the inhibitory-deficit hypothesis, which proposes that older adults have greater difficulty suppressing irrelevant information (Hasher & Zacks, 1988; Kane et al., 1994; Kang et al., 2022). Given these baseline limitations in inhibition and control, cTBS-induced disruption of the DLPFC—critical for attention allocation and inhibition—would be expected to impair performance, as reflected here in delayed P200 and N200 processing.

In addition to these executive deficits, our ERP findings also revealed more general task-related impairments that appeared independent of stimulation and were driven primarily by task difficulty. These included effects on task switching, updating, and decision making, indexed by the P300 component.

The persistence of hemispheric specialisation in ageing was also evident. The selective disruption in the verbal task supports the central role of the left DLPFC in verbal WM, whereas visuospatial WM relies more heavily on right-lateralised networks (Chen et al., 2008; Owen et al., 2005; Wischnewski et al., 2024). Studies targeting the right DLPFC similarly report selective disruption of visuospatial but not verbal WM (Fried et al., 2014), even without overt behavioural impairments.

Overall, inhibitory cTBS to the DLPFC appears to impair the temporal coordination of neural activity, with behavioural consequences depending on the alignment between stimulation site and task domain. In verbal WM, where the left DLPFC is central, this manifests as impaired learning and maladaptive timing across early (P200–N200) and late (P300) processing stages.

In contrast, visuospatial WM—supported by more distributed or right-dominant networks—shows timing alterations that are less behaviourally consequential, reflecting compensatory recruitment within the fronto-parietal system.

### Limitations, strengths and future directions

Several limitations warrant discussion. First, although a baseline session was included, it was not repeated across stimulation conditions, limiting the ability to separate stimulation-specific effects from session-related influences (e.g., fatigue, habituation). Replicating baseline across conditions would, however, have substantially increased participant burden for this older sample; to mitigate this, stimulation order was counterbalanced and fatigue monitored. Second, stimulation sites were defined using the 10–10 EEG system rather than MRI-guided neuronavigation. While F3 reliably targets the left DLPFC at the group level, individual neuroanatomical variability may have reduced precision. Third, the focus on ERP latencies provided important insight into temporal dynamics but did not capture network-level effects that might be revealed by connectivity or time–frequency analyses.

Despite these limitations, the study has several strengths. It is, to our knowledge, the first to combine cTBS and EEG to examine the causal contribution of the DLPFC to WM in older adults. The inclusion of both verbal and visuospatial tasks across two cognitive loads enabled detailed examination of domain-specific and lateralised effects. The integration of behavioural, electrophysiological, and Bayesian approaches further strengthened interpretability and reliability.

Future research should replicate these findings in larger and more diverse samples, given individual variability in responsiveness to NIBS. Direct comparisons with younger adults and studies targeting the right DLPFC would clarify age- and hemisphere-specific effects.

Incorporating MRI-guided neuronavigation and multimodal imaging (e.g., fMRI, MEG) would improve target precision and illuminate changes in connectivity following cTBS. Longitudinal and multi-session designs could explore the persistence and plasticity of stimulation-induced changes in ageing.

## CONCLUSION

This study provides novel causal evidence for the role of the DLPFC in WM in older adults. Although cTBS did not induce robust behavioural impairments, ERP analyses revealed latency modulations across components associated with early attention (P200), conflict monitoring (N200), and decision making (P300), particularly in the verbal N-back. P200 and N200 latencies predicted performance, highlighting the relevance of temporal precision for behavioural outcomes. In contrast, visuospatial WM was largely unaffected by left DLPFC stimulation, supporting domain-specific hemispheric organisation. Together, these findings demonstrate the value of ERP latency markers in detecting subtle cognitive and neuroplastic changes and underscore the importance of processing speed and network timing for cognitive performance in ageing.

## DECLARATIONS

### Ethics approval

The study was approved by the Edge Hill University Ethics Research Committee (Reference: ETH2122-0155).

### Consent to participate

Informed consent was obtained from all individual participants included in the study.

### Consent for publication

All participants provided consent for anonymised data to be published in academic journals and presentations.

### Availability of data and materials

The data and materials used in this study are openly available on the Open Science Framework (OSF) at: https://osf.io/pfyj2/?view_only=b1f7f3adc2d546cb870e174d944d61e2

### Code availability

The code used for data collection and analysis is available at the same OSF repository: https://osf.io/pfyj2/?view_only=b1f7f3adc2d546cb870e174d944d61e2

## SUPPLEMENTARY MATERIAL

**Table S1.** Summary of statistical analyses and results for the Verbal N-back – ERP amplitudes.

| Analysis | Model / Effect | Test statistics | *p*-value | Effect size |
| --- | --- | --- | --- | --- |
| P200 | LMEM Model | AIC = 5499.63 | – | R<sup>2</sup>m = 0.03; |
|  | 3 (Condition) × 2 (Cognitive load) |  |  | R<sup>2</sup>c = 0.41 |
|  | × 2 (Stimulus type) × 8 (Channel) |  |  |  |
|  | with random intercepts for |  |  |  |
|  | Participants |  |  |  |
| | Condition | $F(2, 2033) = 0.48$ | .62 | – |
| | Cognitive load | $F(1, 2033) = 0.64$ | .42 | – |
| | Stimulus type | $F(1, 2033) = 3.07$ | .08 | – |
| | Channel | $F(8, 2033) = 5.13$ | < .001* | $\eta^2 = 0.02$ |
| | Condition × Cognitive load | $F(2, 2033) = 0.40$ | .67 | – |
| | Condition × Stimulus type | $F(2, 2033) = 0.35$ | .71 | – |
| | Condition × Channel | $F(16, 2033) = 0.35$ | .99 | – |
| | Cognitive load × Stimulus type | $F(1, 2033) = 5.07$ | < .05* | $\eta^2 = 0.002$ |
| | Condition × Cognitive load ×<br>Stimulus type | $F(2, 2033) = 0.65$ | .52 | – |
| Post-hoc contrasts<br>(Tukey) | Target – Non-target (2-back)<br>(coefficients) | $B = 0.14, SE = 0.05,$<br>$t(2033) = 2.83$ | < .05* | $d = 0.14$ |
| | Target – Non-target (3-back)<br>(coefficients) | $B = -0.02, SE = 0.05,$<br>$t(2033) = -0.35$ | .99 | – |
| N200 | LMEM Model<br>3 (Condition) × 2 (Cognitive load)<br>× 2 (Stimulus type) × 8 (Channel) | AIC = 6519.78 | – | R<sup>2</sup>m = 0.03;<br>R<sup>2</sup>c = 0.22 |
|  | with random intercepts for<br>Participants |  |  |  |
| | Condition | $F(2, 2033) = 1.23$ | .29 | — |
| | Cognitive load | $F(1, 2033) = 2.79$ | .09 | — |
| | Stimulus type | $F(1, 2033) = 3.74$ | .05* | $\eta^2 = 0.002$ |
| | Channel | $F(8, 2033) = 2.07$ | < .05* | $\eta^2 = 0.01$ |
| | Condition × Cognitive load | $F(2, 2033) = 0.02$ | .98 | — |
| | Condition × Stimulus type | $F(2, 2033) = 0.86$ | .42 | — |
| | Condition × Channel | $F(16, 2033) = 0.57$ | .91 | — |
| | Cognitive load × Stimulus type | $F(1, 2033) = 0.96$ | .33 | — |
| | Condition × Cognitive load ×<br>Stimulus type | $F(2, 2033) = 0.13$ | .87 | – |
| Post-hoc contrasts<br>(Tukey) | Target – Non-target<br>(coefficients) | $B = -0.09, SE = 0.04,$<br>$t(2033) = -1.94$ | .05* | $d = -0.06$ |
| P300 | LMEM Model<br>3 (Condition) × 2 (Cognitive load)<br>× 2 (Stimulus type) × 8 (Channel)<br>with random intercepts for<br>Participants | $AIC = 7210.44$ | – | $R^2m = 0.02;$<br>$R^2c = 0.67$ |
| | Condition | $F(2, 2033) = 0.60$ | .55 | – |
| | Cognitive load | $F(1, 2033) = 1.73$ | .19 | – |
| | Stimulus type | $F(1, 2033) = 2.93$ | .09 | – |
| | Channel | $F(8, 2033) = 8.23$ | < .001* | $\eta^2 = 0.03$ |
| | Condition × Cognitive load | $F(2, 2033) = 1.01$ | .36 | — |
| | Condition × Stimulus type | $F(2, 2033) = 0.69$ | .50 | — |
| | Condition × Channel | $F(16, 2033) = 1.30$ | .19 | — |
| | Cognitive load × Stimulus type | $F(1, 2033) = 0.30$ | .59 | — |
| | Condition × Cognitive load ×<br>Stimulus type | $F(2, 2033) = 0.25$ | .78 | — |
| Post-hoc contrasts<br>(Tukey) | — | — | — | — |

*Note.* LMEM = linear mixed-effects model; AIC = Akaike Information Criterion; $R^2_m$ = marginal $R^2$ ; $R^2_c$ = conditional $R^2$ . Significant effects are marked with an asterisk (\*). Effect sizes ( $d$ , $\eta^2$ , $R^2$ , $Adj. R^2$ ) are reported only for significant effects. Effect size benchmarks for $\eta^2$ : small = .01, medium = .06, large = .14 (Lakens, 2013). Effect size benchmarks for Cohen's $d$ : small = .02, medium = .05, large = .08 (Cohen, 1988).

**Table S2.**
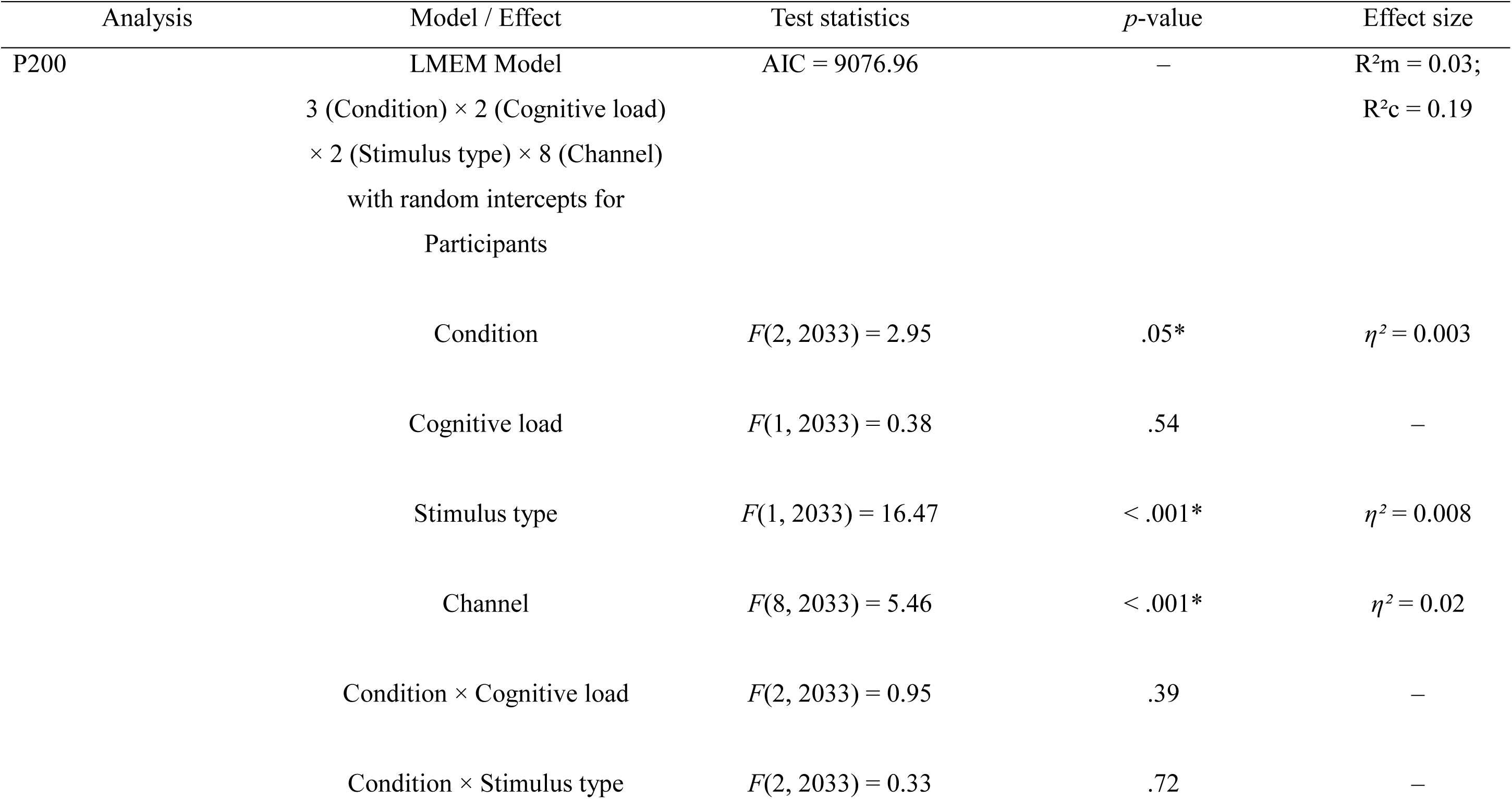
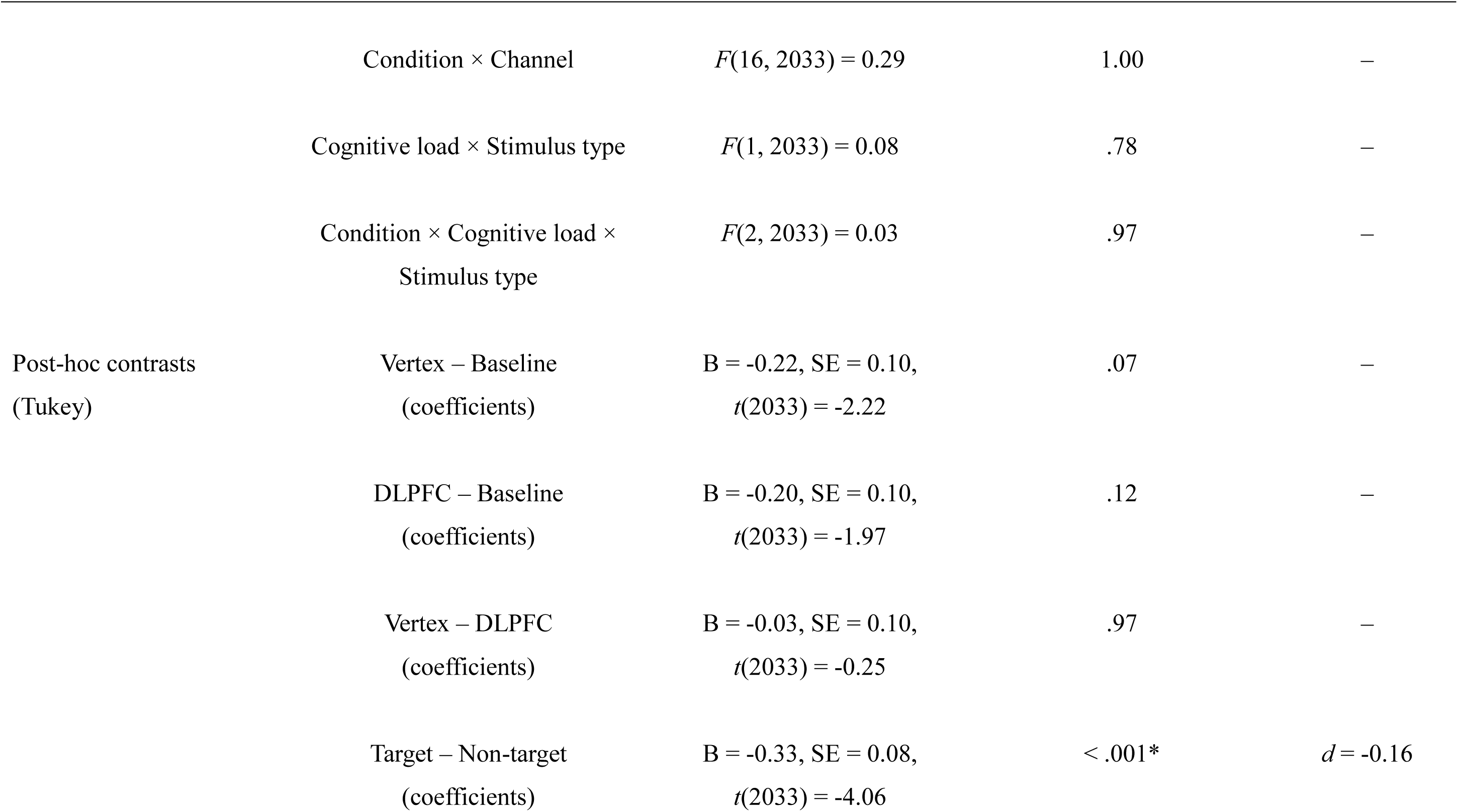
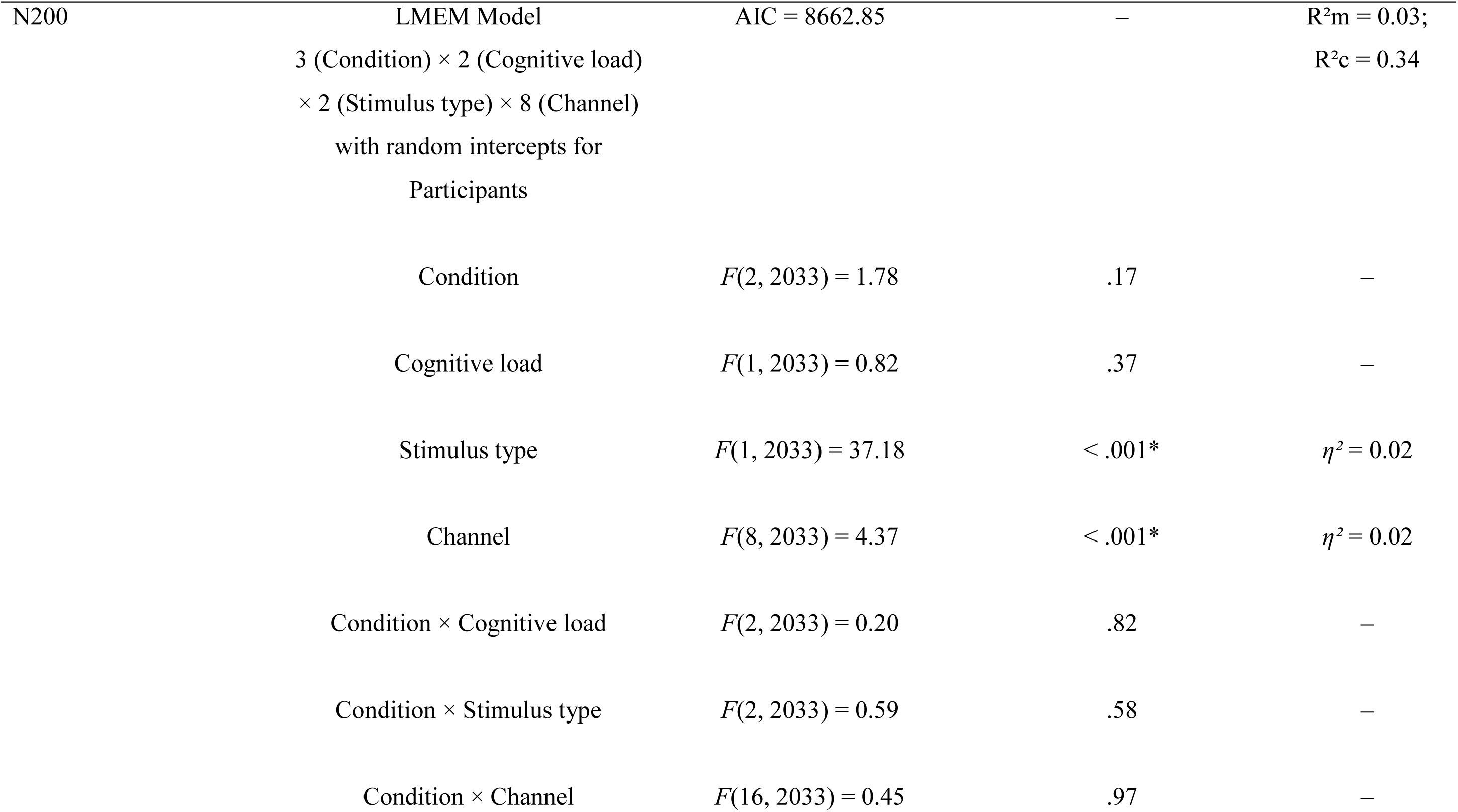
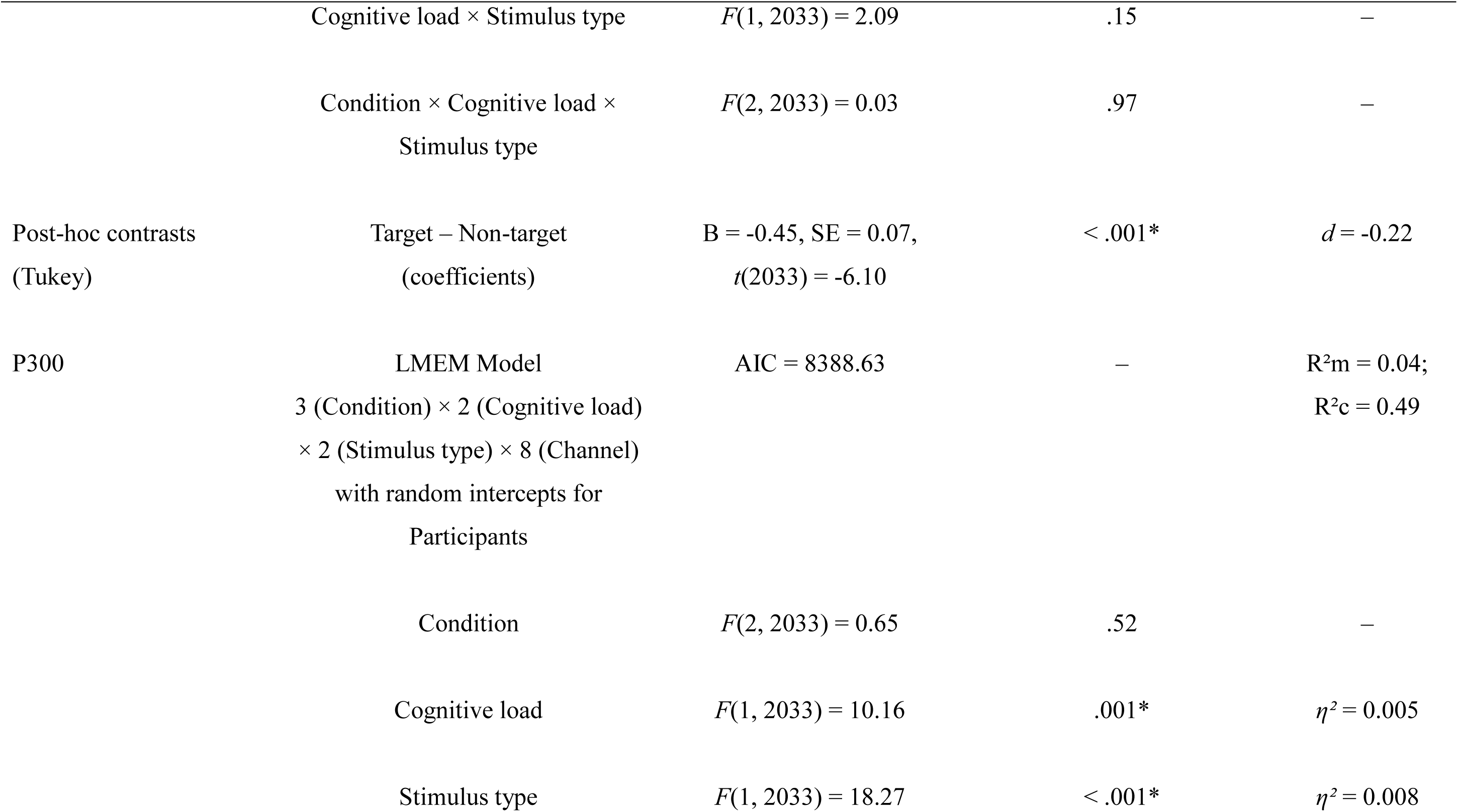
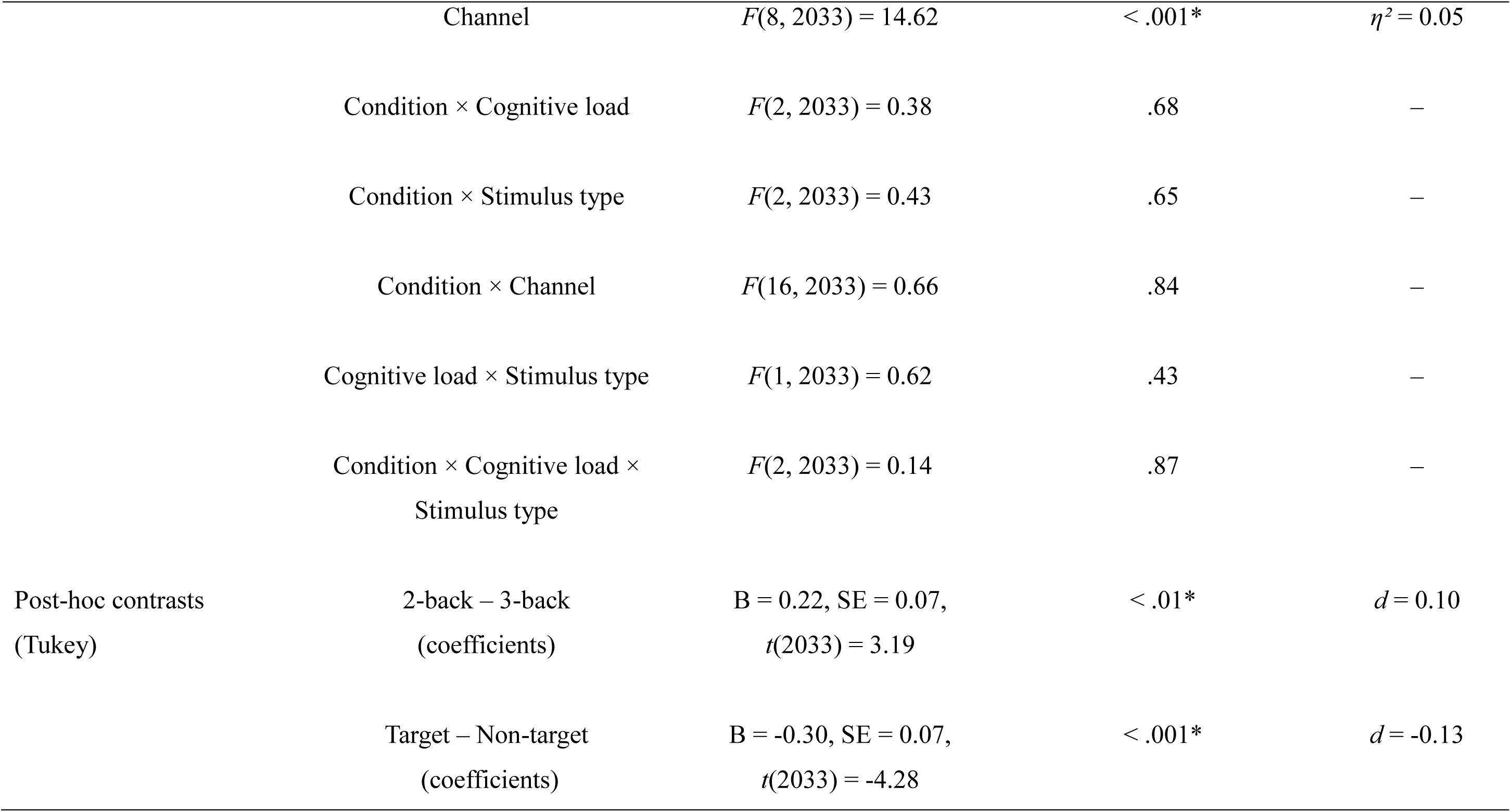
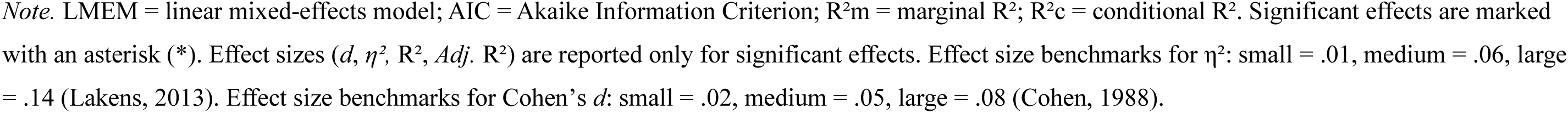
Summary of statistical analyses and results for the Visuospatial N-back – ERP amplitudes.

| Analysis | Model / Effect | Test statistics | *p*-value | Effect size |
| --- | --- | --- | --- | --- |
| P200 | LMEM Model | AIC = 9076.96 | – | R<sup>2</sup><sub>m</sub> = 0.03; |
|  | 3 (Condition) × 2 (Cognitive load) |  |  | R<sup>2</sup><sub>c</sub> = 0.19 |
|  | × 2 (Stimulus type) × 8 (Channel) |  |  |  |
|  | with random intercepts for |  |  |  |
|  | Participants |  |  |  |
| | Condition | $F(2, 2033) = 2.95$ | .05* | $\eta^2 = 0.003$ |
| | Cognitive load | $F(1, 2033) = 0.38$ | .54 | – |
| | Stimulus type | $F(1, 2033) = 16.47$ | < .001* | $\eta^2 = 0.008$ |
| | Channel | $F(8, 2033) = 5.46$ | < .001* | $\eta^2 = 0.02$ |
| | Condition × Cognitive load | $F(2, 2033) = 0.95$ | .39 | – |
| | Condition × Stimulus type | $F(2, 2033) = 0.33$ | .72 | – |
| | Condition × Channel | $F(16, 2033) = 0.29$ | 1.00 | — |
| | Cognitive load × Stimulus type | $F(1, 2033) = 0.08$ | .78 | — |
| | Condition × Cognitive load ×<br>Stimulus type | $F(2, 2033) = 0.03$ | .97 | — |
| Post-hoc contrasts<br>(Tukey) | Vertex – Baseline<br>(coefficients) | B = -0.22, SE = 0.10,<br>$t(2033) = -2.22$ | .07 | — |
| | DLPFC – Baseline<br>(coefficients) | B = -0.20, SE = 0.10,<br>$t(2033) = -1.97$ | .12 | — |
| | Vertex – DLPFC<br>(coefficients) | B = -0.03, SE = 0.10,<br>$t(2033) = -0.25$ | .97 | — |
| | Target – Non-target<br>(coefficients) | B = -0.33, SE = 0.08,<br>$t(2033) = -4.06$ | < .001* | $d = -0.16$ |
| N200 | LMEM Model | AIC = 8662.85 | – | R<sup>2</sup>m = 0.03; |
|  | 3 (Condition) × 2 (Cognitive load) |  |  | R<sup>2</sup>c = 0.34 |
|  | × 2 (Stimulus type) × 8 (Channel) |  |  |  |
|  | with random intercepts for |  |  |  |
|  | Participants |  |  |  |
|  | Condition | *F*(2, 2033) = 1.78 | .17 | – |
|  | Cognitive load | *F*(1, 2033) = 0.82 | .37 | – |
| | Stimulus type | *F*(1, 2033) = 37.18 | < .001* | $\eta^2 = 0.02$ |
| | Channel | *F*(8, 2033) = 4.37 | < .001* | $\eta^2 = 0.02$ |
|  | Condition × Cognitive load | *F*(2, 2033) = 0.20 | .82 | – |
|  | Condition × Stimulus type | *F*(2, 2033) = 0.59 | .58 | – |
|  | Condition × Channel | *F*(16, 2033) = 0.45 | .97 | – |
| | Cognitive load × Stimulus type | $F(1, 2033) = 2.09$ | .15 | – |
| | Condition × Cognitive load ×<br>Stimulus type | $F(2, 2033) = 0.03$ | .97 | – |
| Post-hoc contrasts<br>(Tukey) | Target – Non-target<br>(coefficients) | $B = -0.45, SE = 0.07,$<br>$t(2033) = -6.10$ | < .001* | $d = -0.22$ |
| P300 | LMEM Model<br>3 (Condition) × 2 (Cognitive load)<br>× 2 (Stimulus type) × 8 (Channel)<br>with random intercepts for<br>Participants | AIC = 8388.63 | – | R<sup>2</sup>m = 0.04;<br>R<sup>2</sup>c = 0.49 |
| | Condition | $F(2, 2033) = 0.65$ | .52 | – |
| | Cognitive load | $F(1, 2033) = 10.16$ | .001* | $\eta^2 = 0.005$ |
| | Stimulus type | $F(1, 2033) = 18.27$ | < .001* | $\eta^2 = 0.008$ |
| | Channel | $F(8, 2033) = 14.62$ | < .001* | $\eta^2 = 0.05$ |
| | Condition × Cognitive load | $F(2, 2033) = 0.38$ | .68 | – |
| | Condition × Stimulus type | $F(2, 2033) = 0.43$ | .65 | – |
| | Condition × Channel | $F(16, 2033) = 0.66$ | .84 | – |
| | Cognitive load × Stimulus type | $F(1, 2033) = 0.62$ | .43 | – |
| | Condition × Cognitive load ×<br>Stimulus type | $F(2, 2033) = 0.14$ | .87 | – |
| Post-hoc contrasts<br>(Tukey) | 2-back – 3-back<br>(coefficients) | B = 0.22, SE = 0.07,<br>$t(2033) = 3.19$ | < .01* | $d = 0.10$ |
| | Target – Non-target<br>(coefficients) | B = -0.30, SE = 0.07,<br>$t(2033) = -4.28$ | < .001* | $d = -0.13$ |

*Note.* LMEM = linear mixed-effects model; AIC = Akaike Information Criterion; R<sup>2</sup>m = marginal R<sup>2</sup>; R<sup>2</sup>c = conditional R<sup>2</sup>. Significant effects are marked with an asterisk (\*). Effect sizes (*d*, $\eta^2$ , R<sup>2</sup>, *Adj.* R<sup>2</sup>) are reported only for significant effects. Effect size benchmarks for $\eta^2$ : small = .01, medium = .06, large = .14 (Lakens, 2013). Effect size benchmarks for Cohen's *d*: small = .02, medium = .05, large = .08 (Cohen, 1988).

**Table S3.**
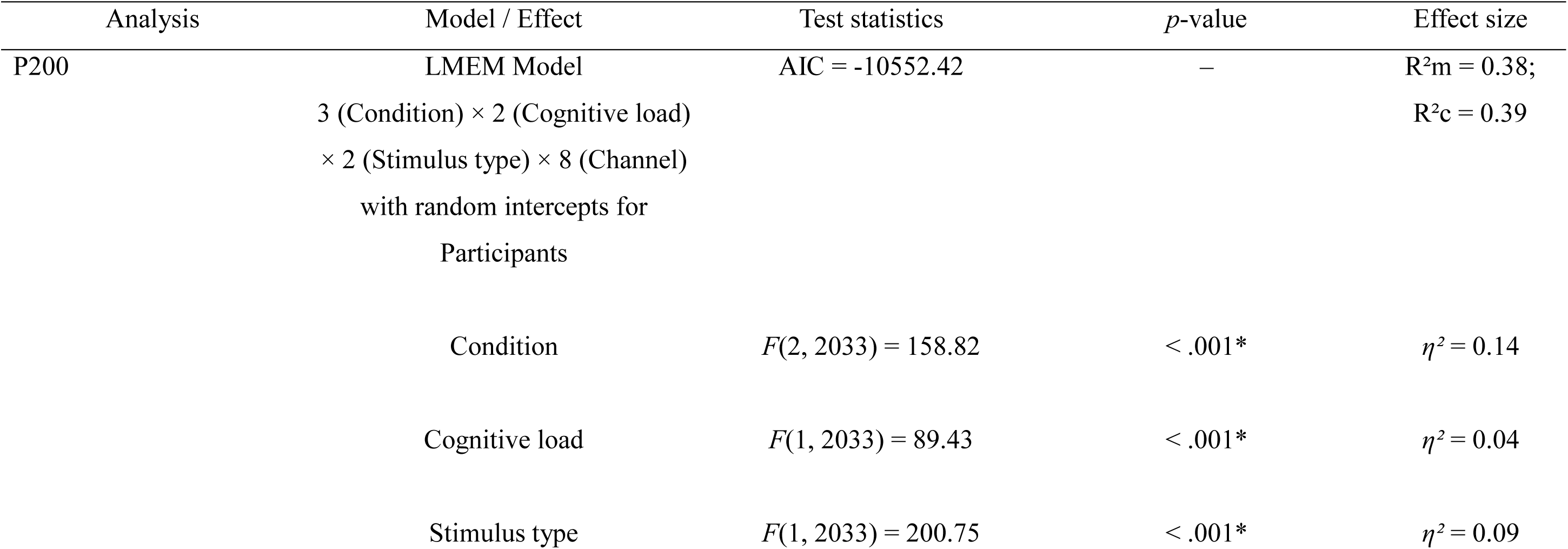
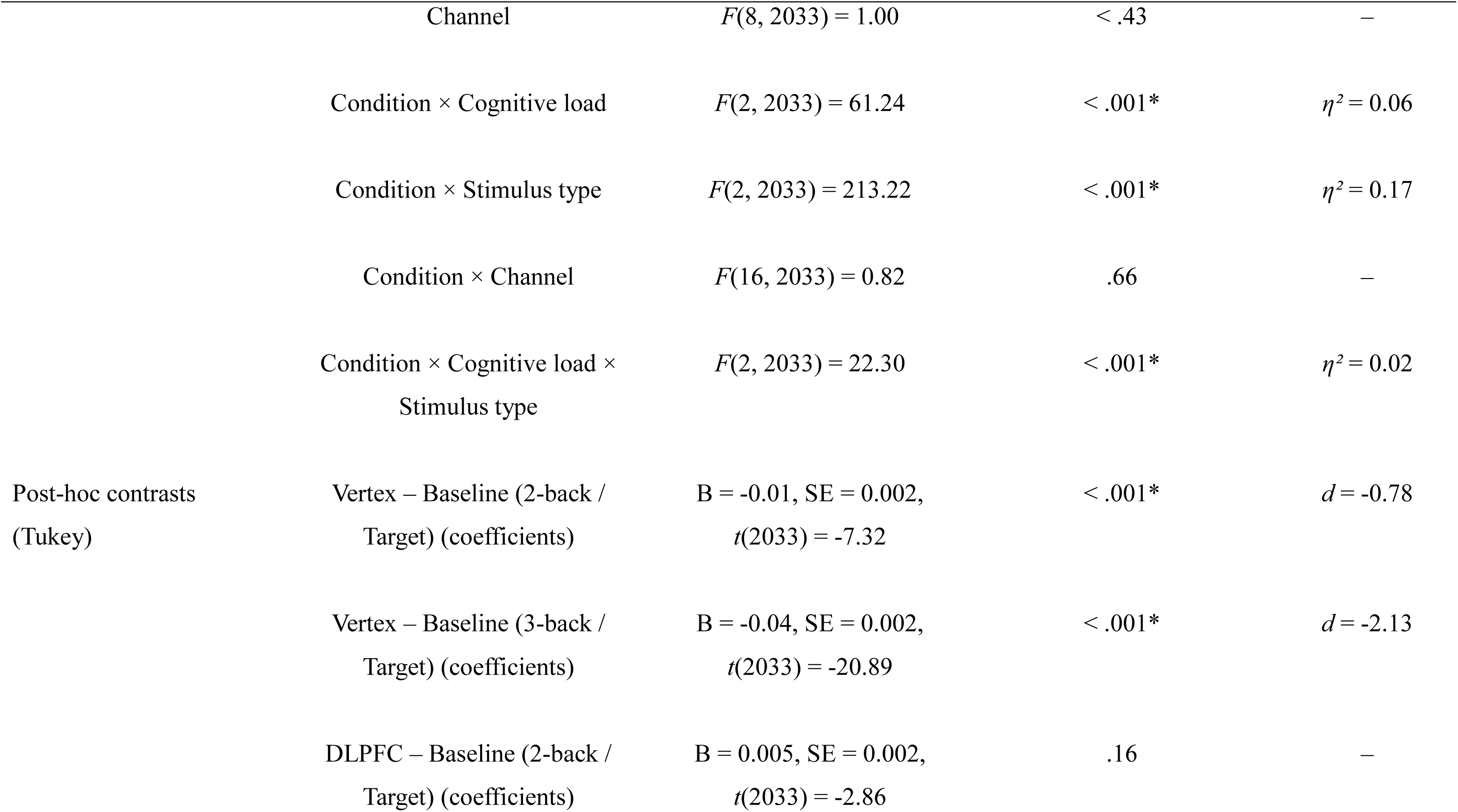
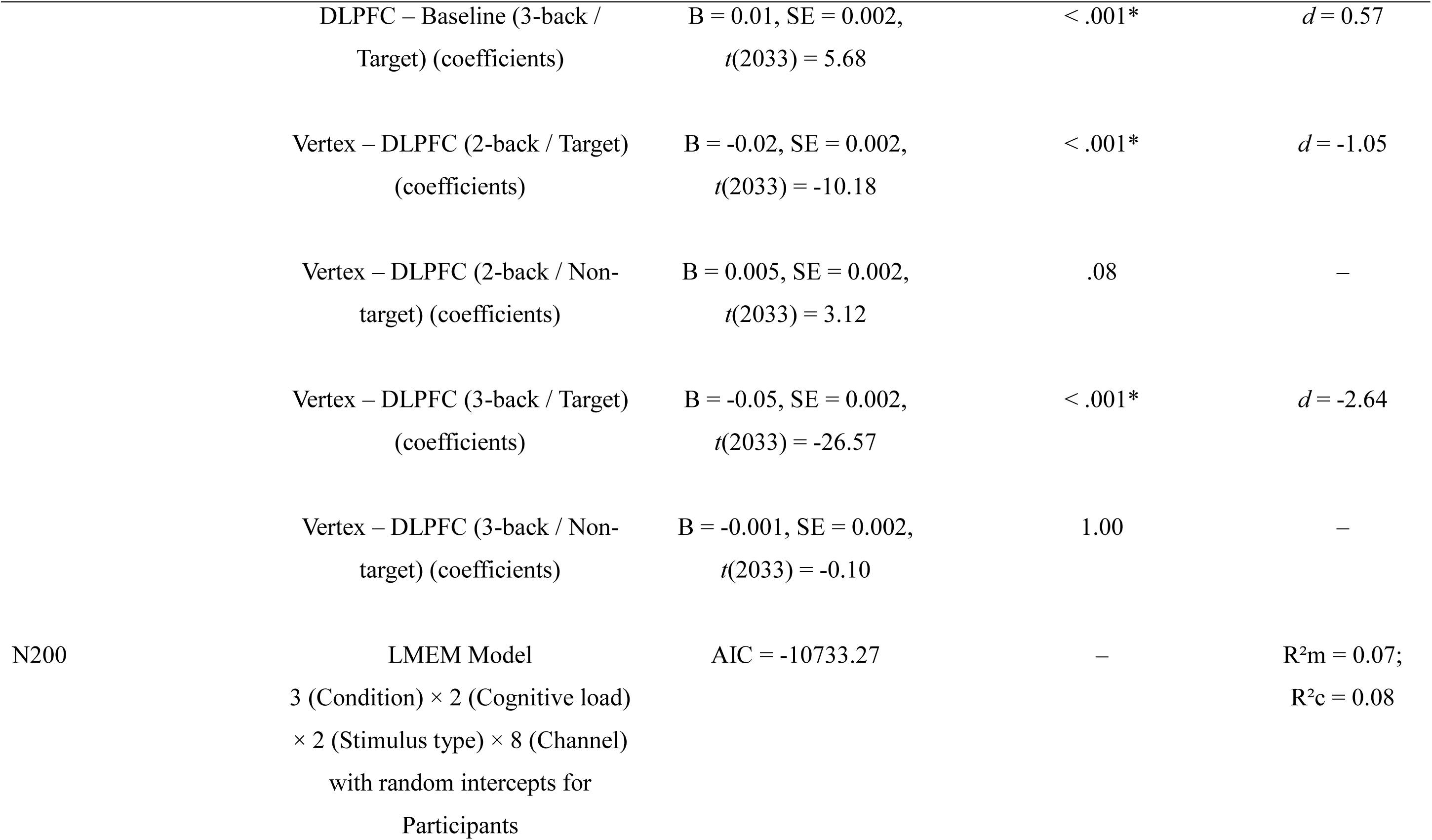
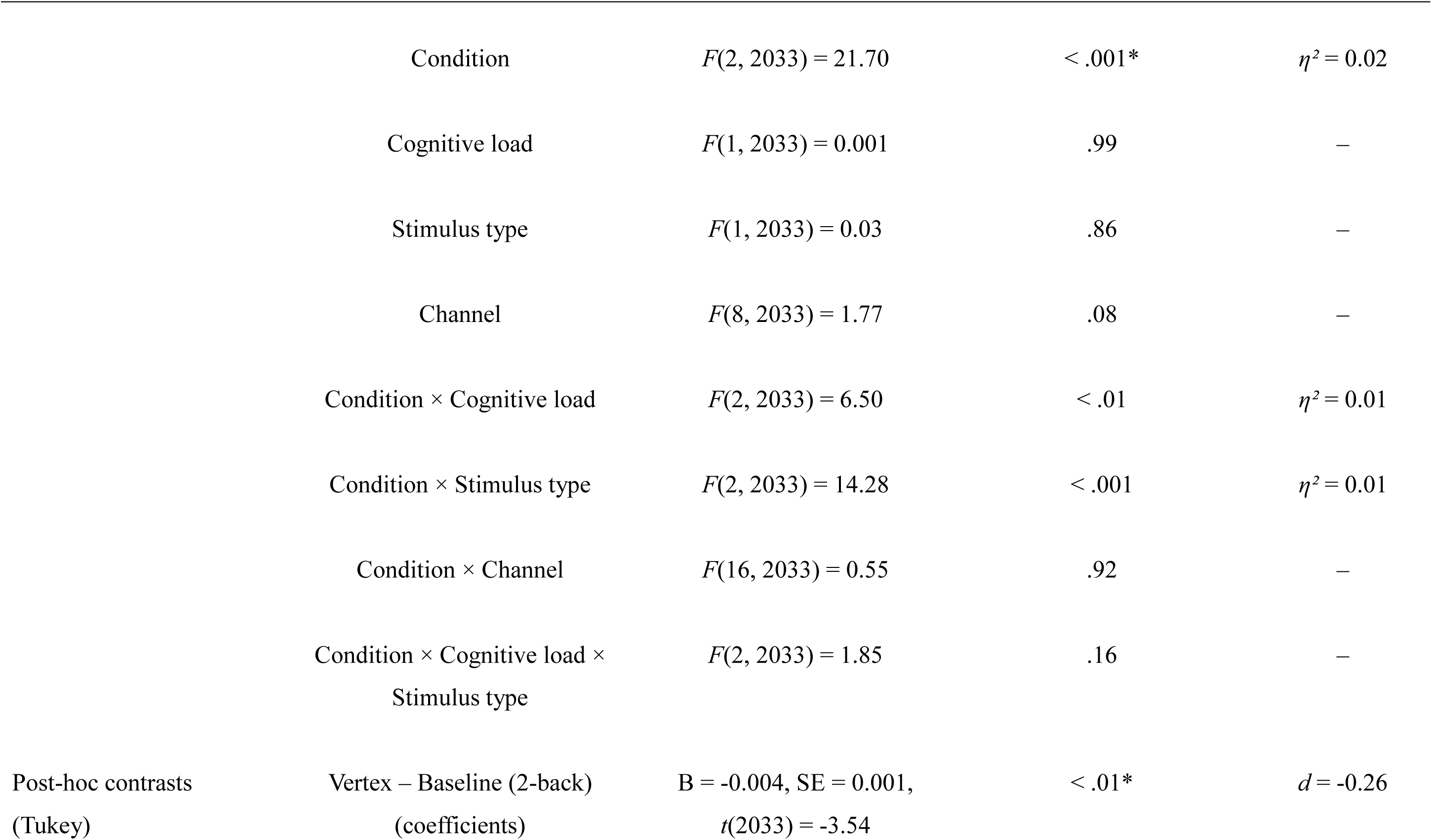
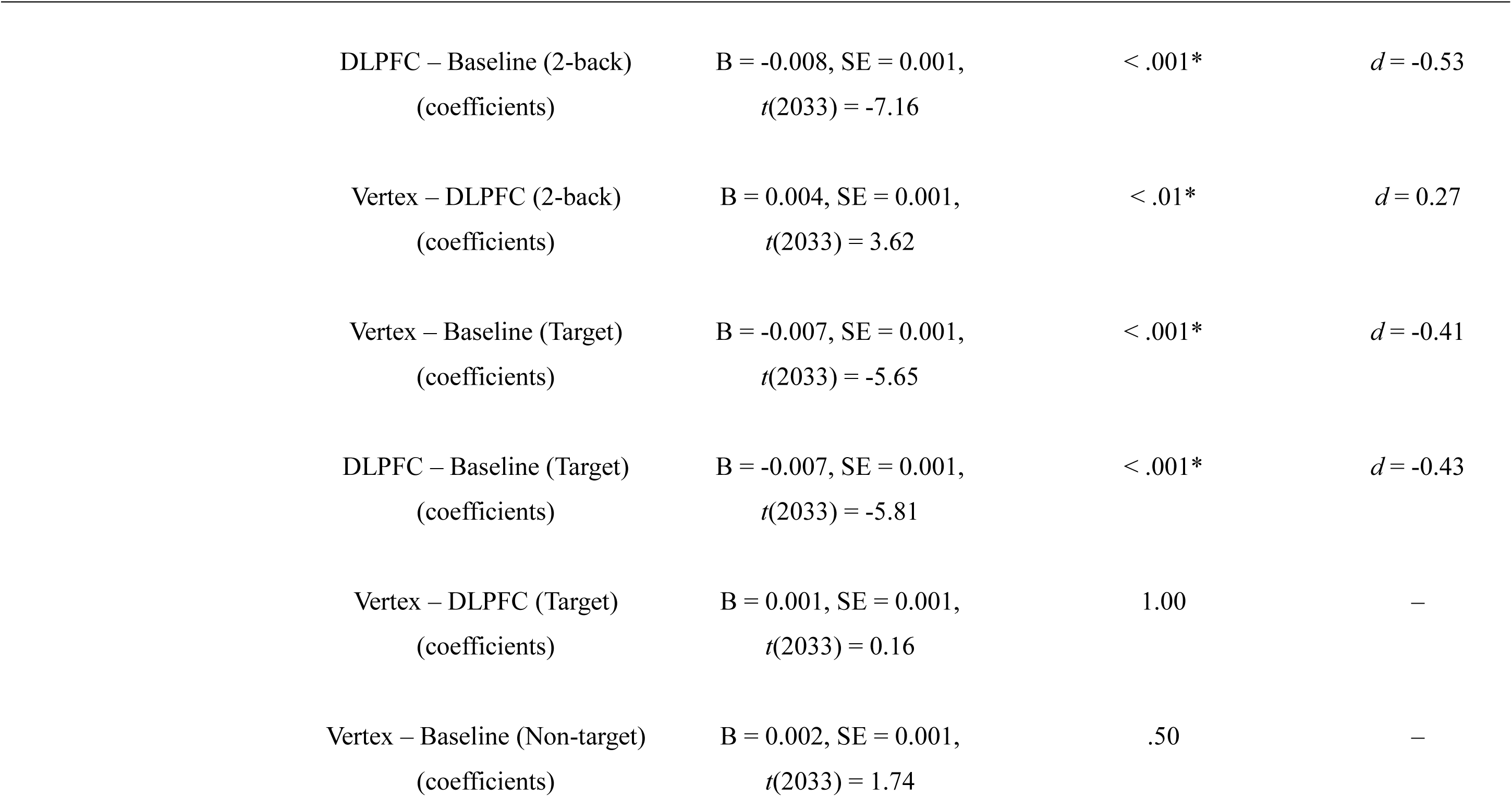
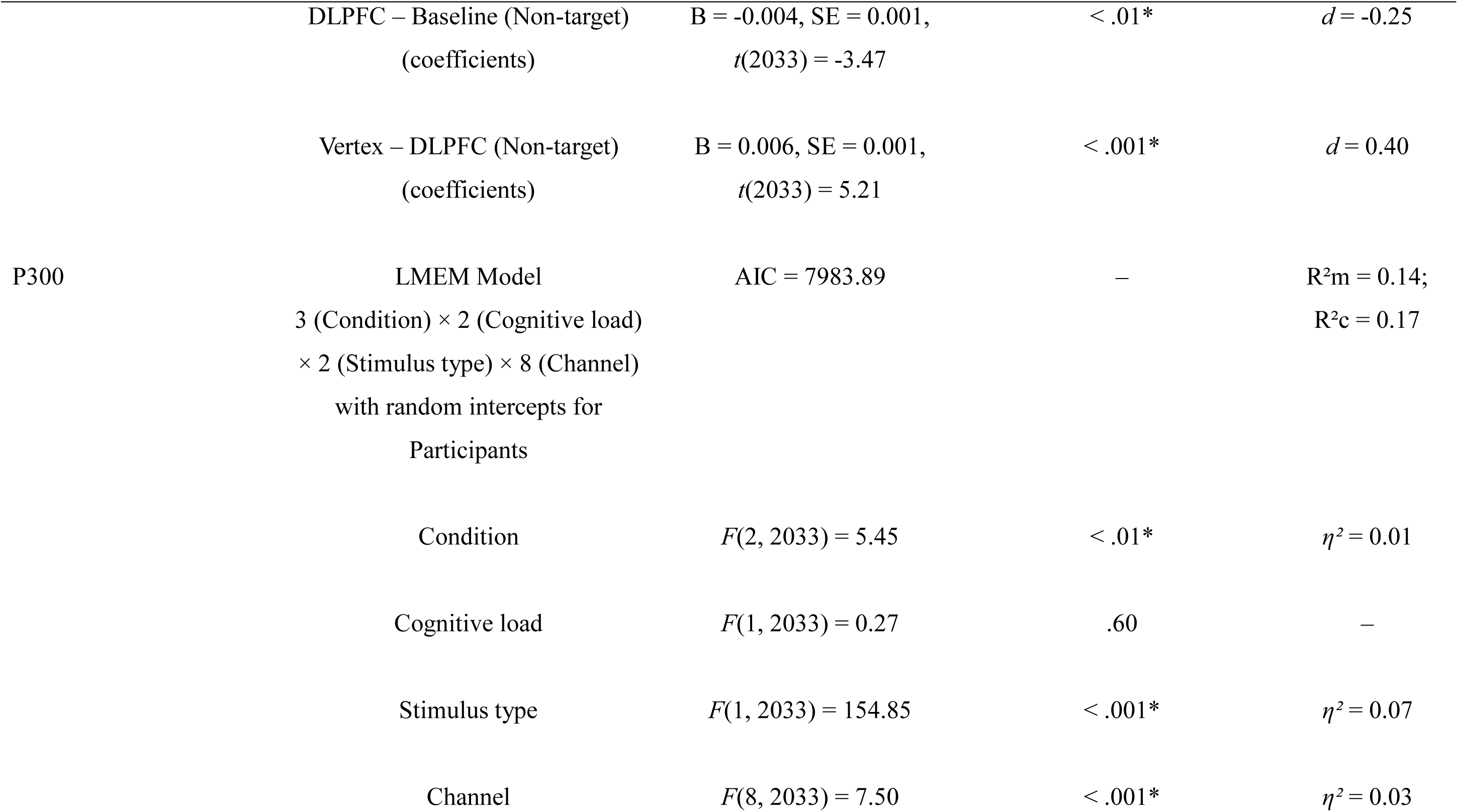
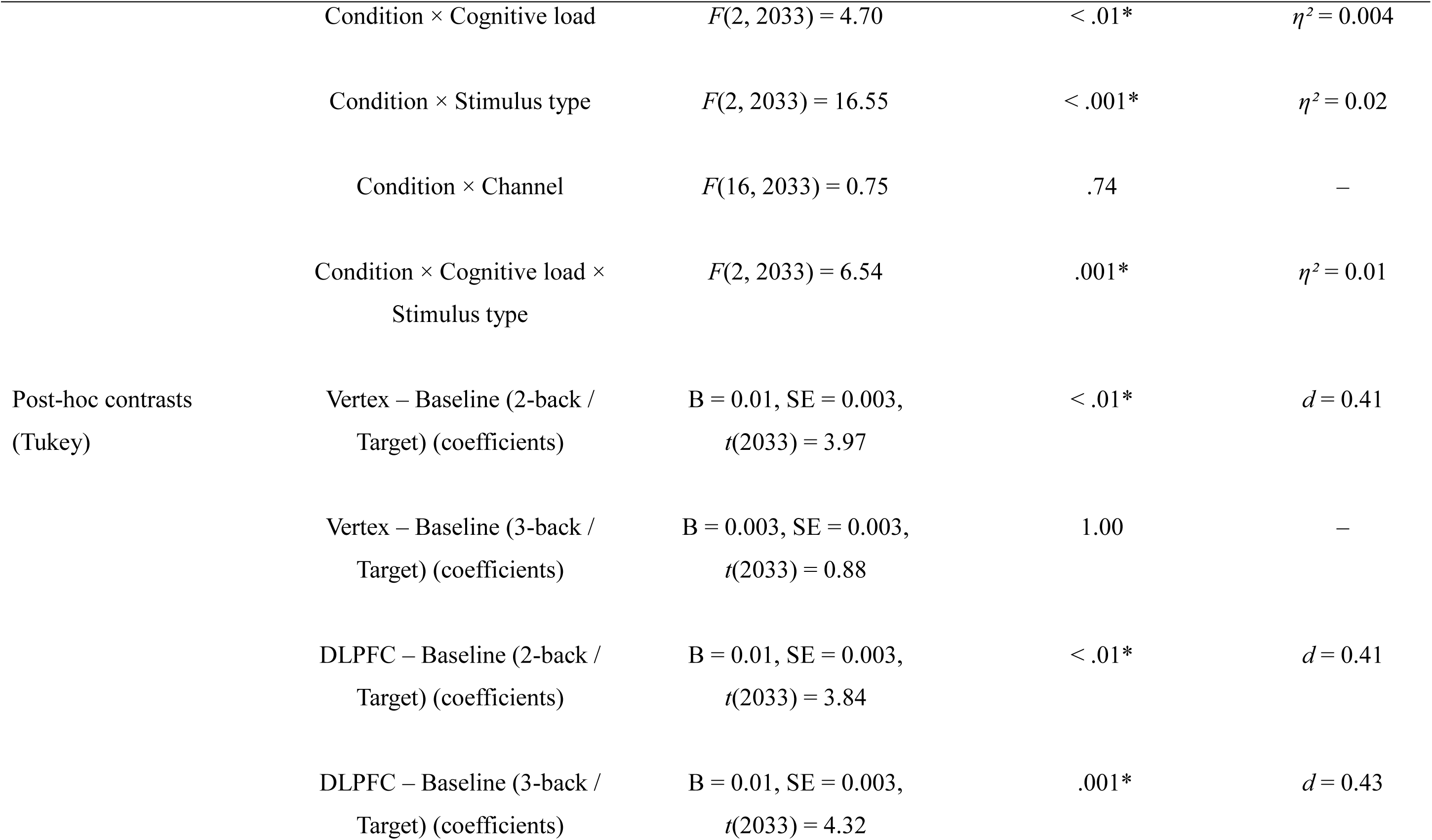
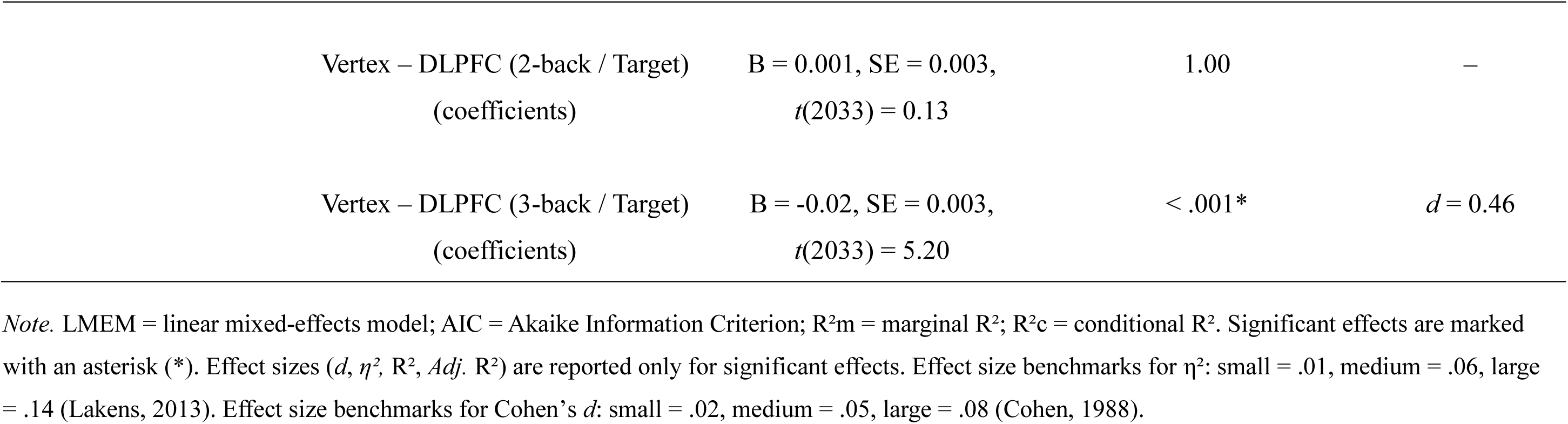
Summary of statistical analyses and results for the Verbal N-back – ERP latencies.

**Table S4.**
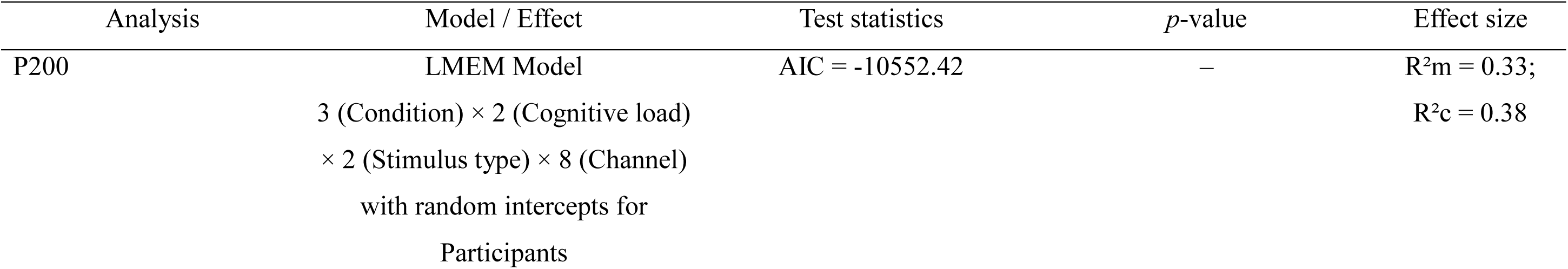
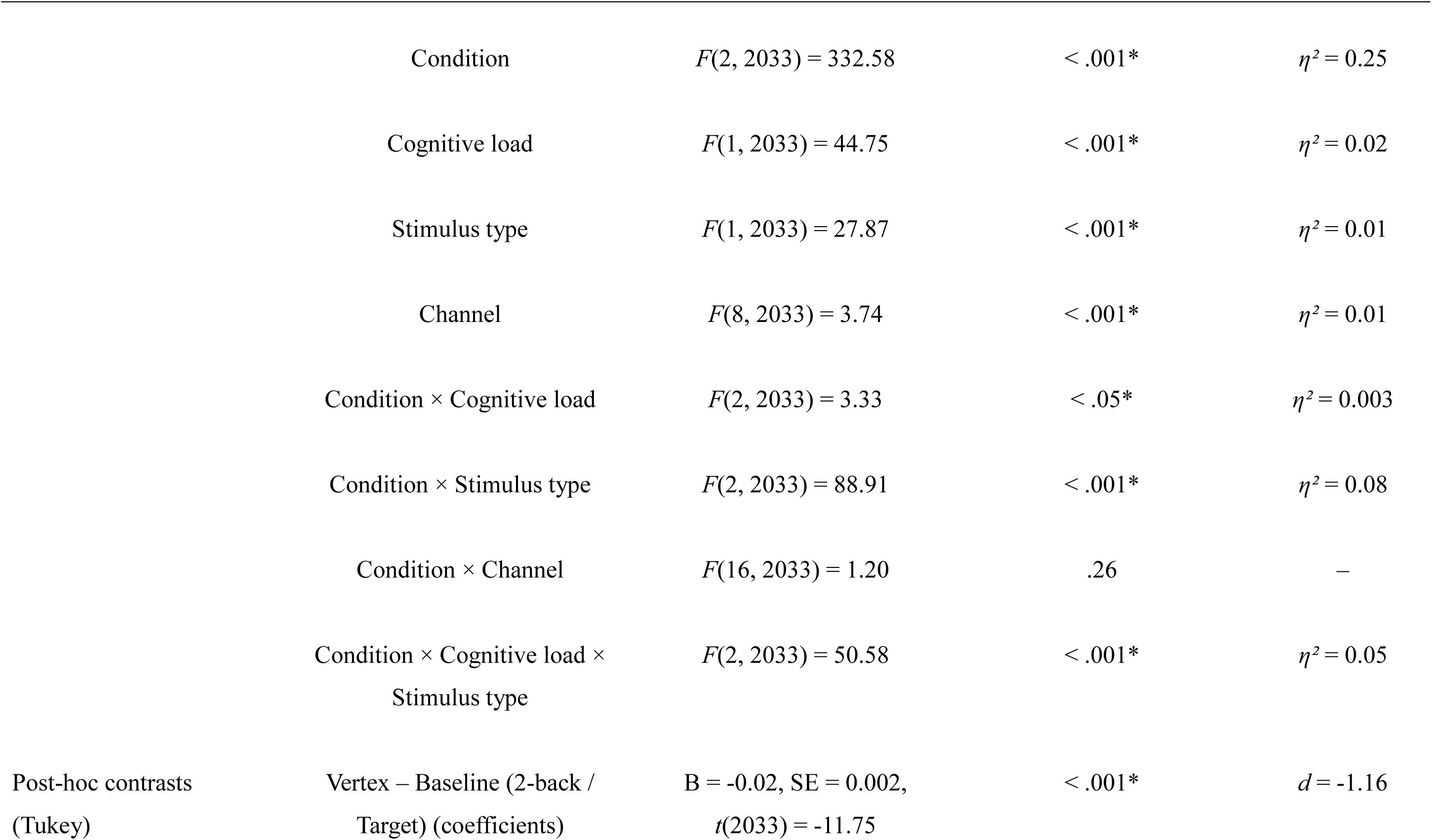
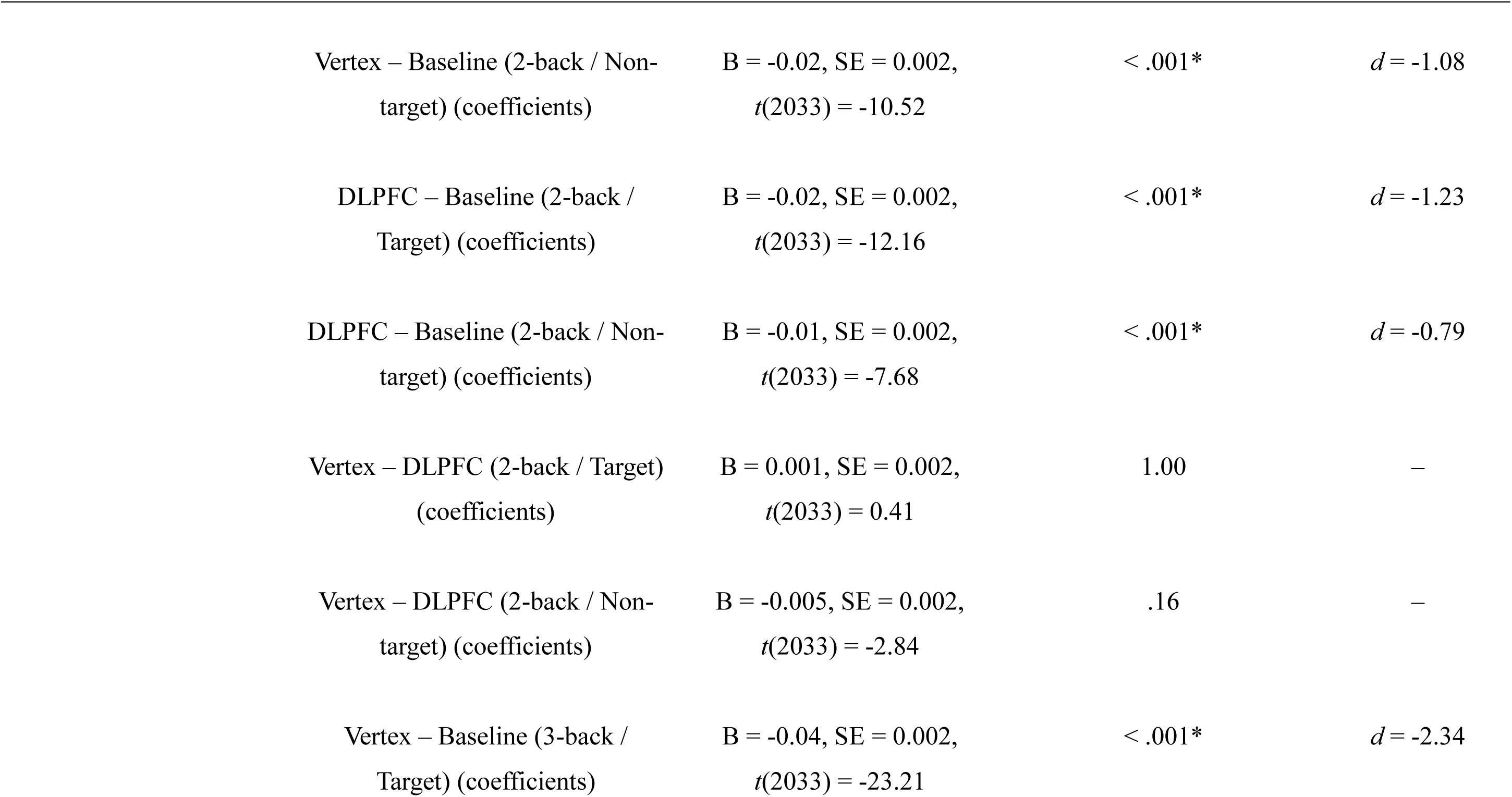
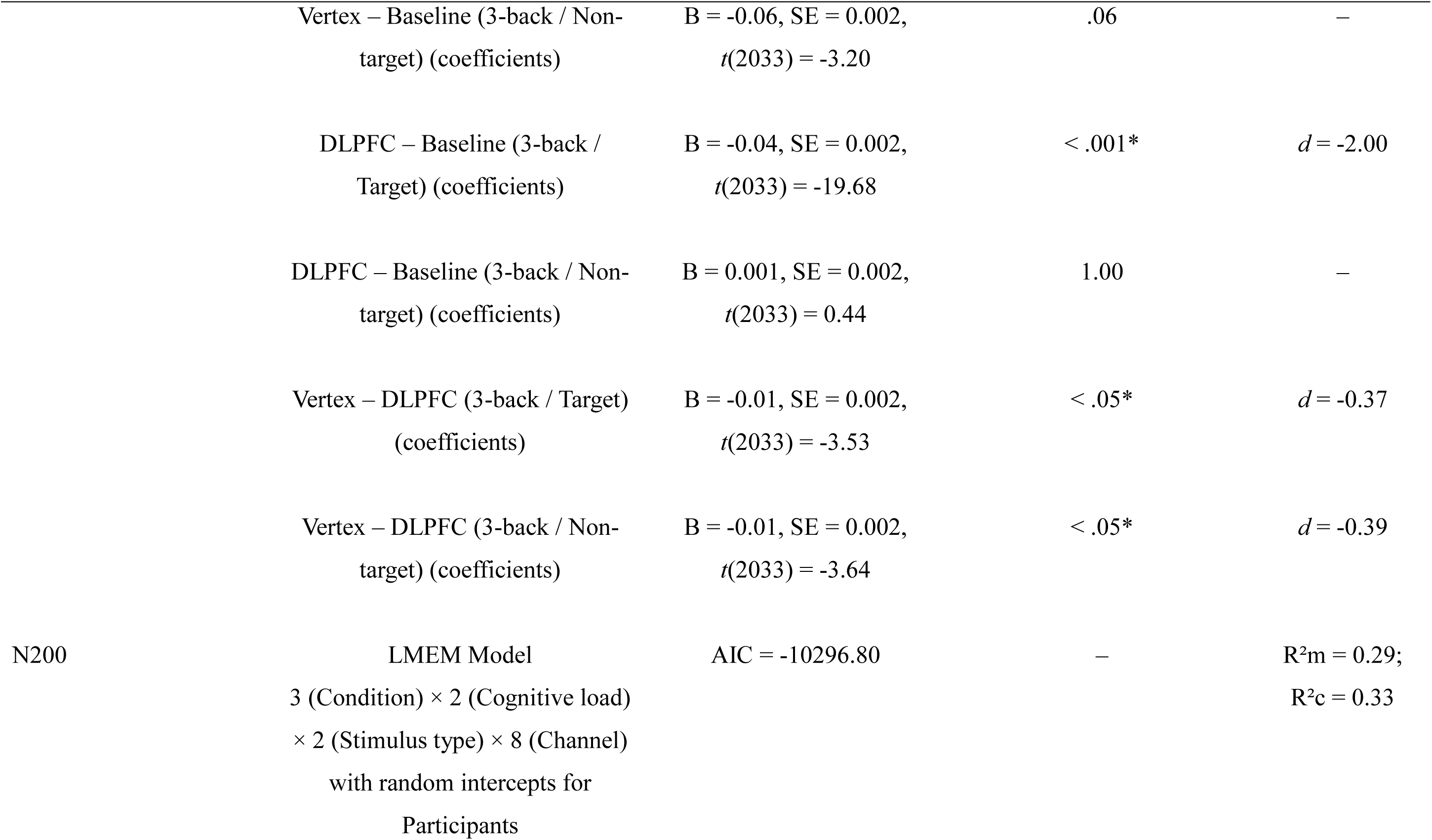
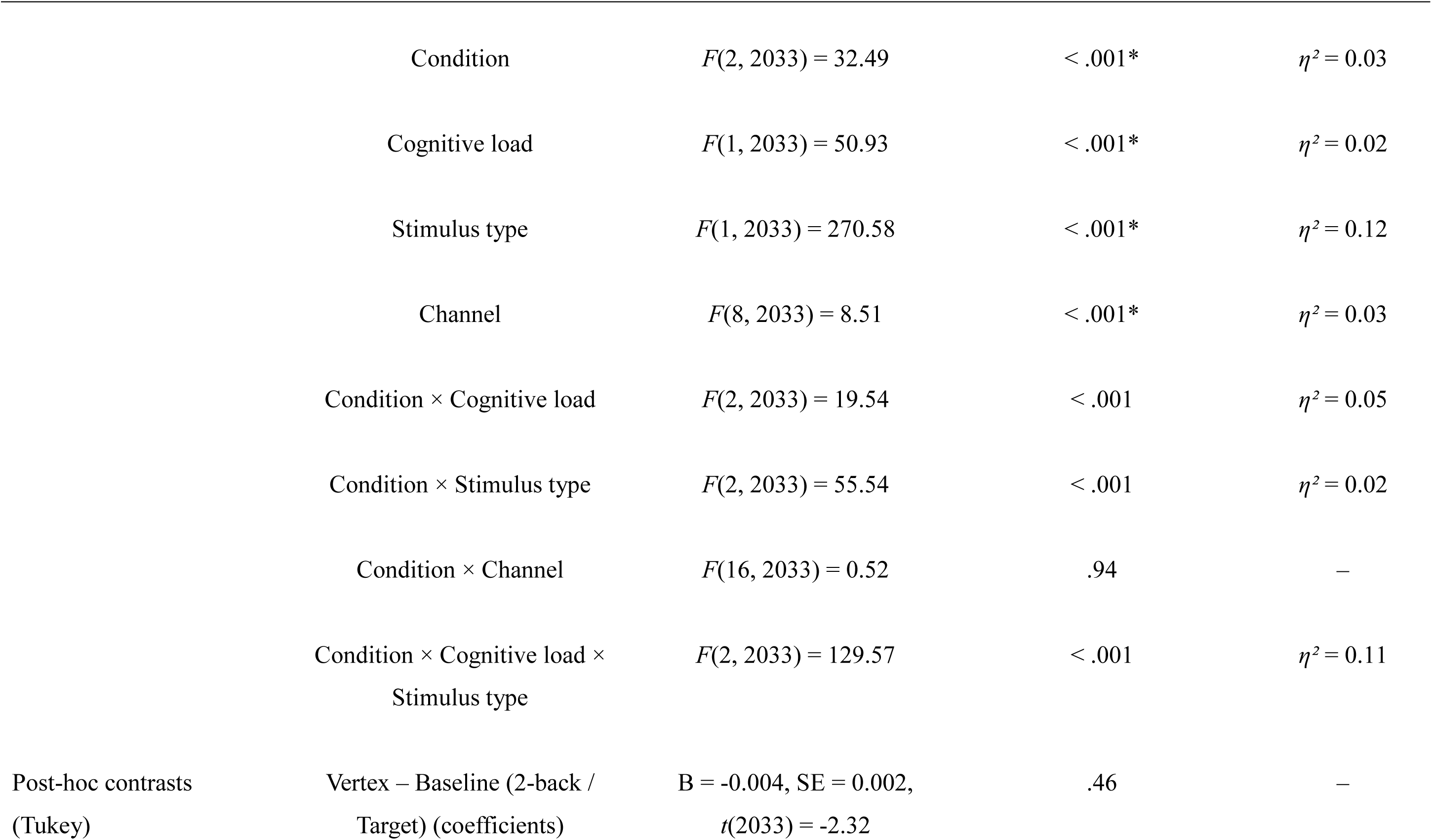
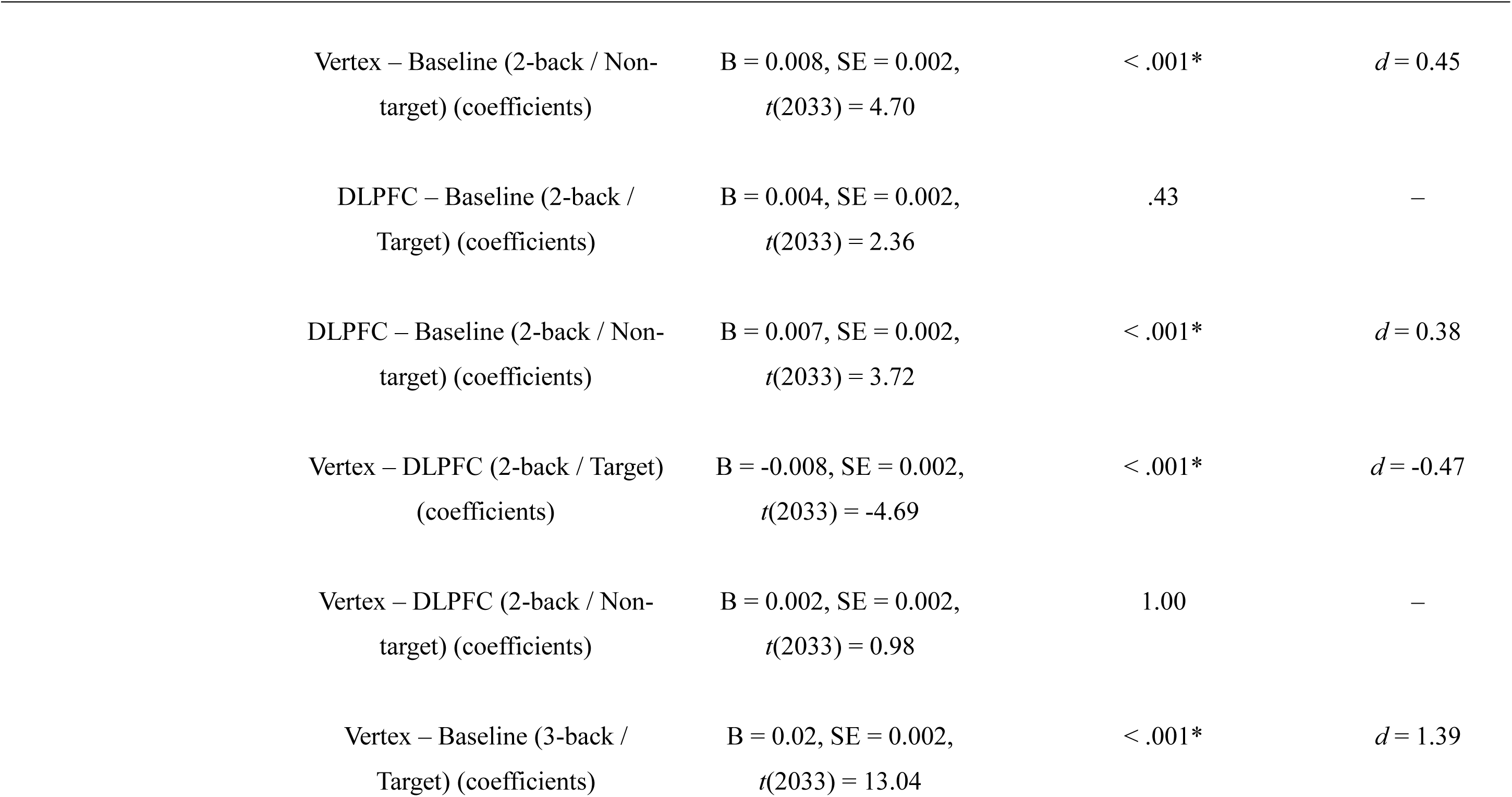
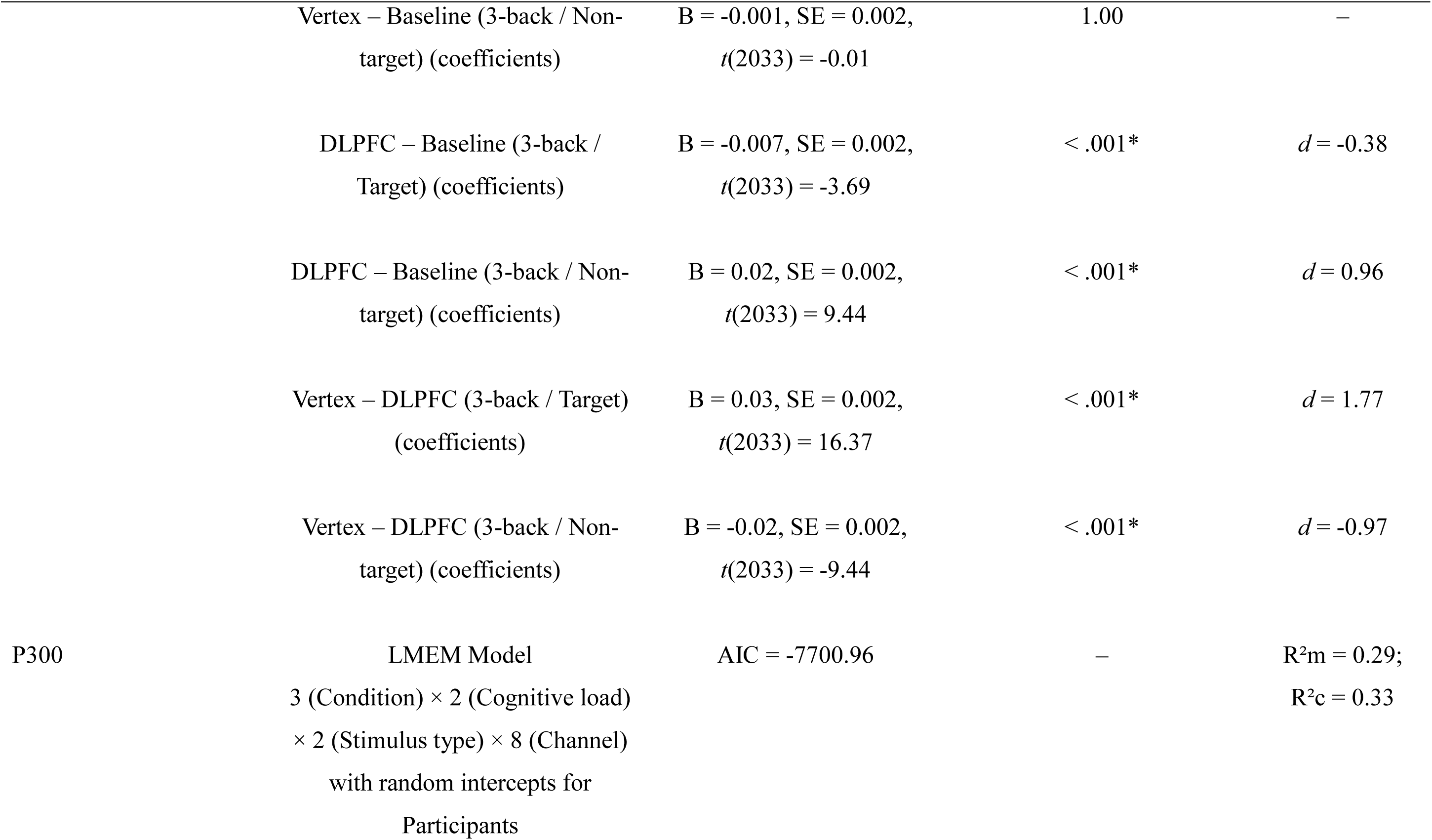
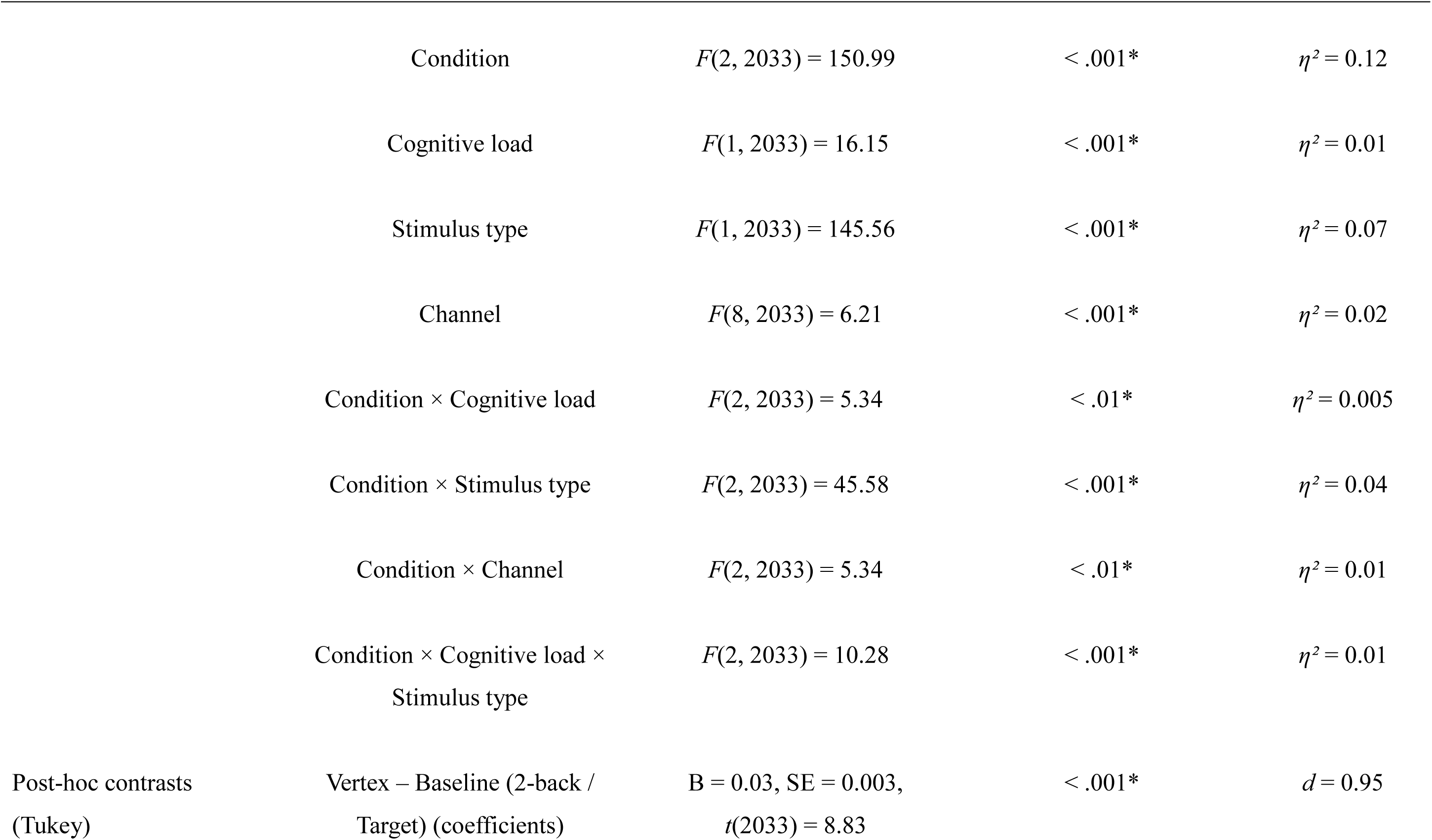
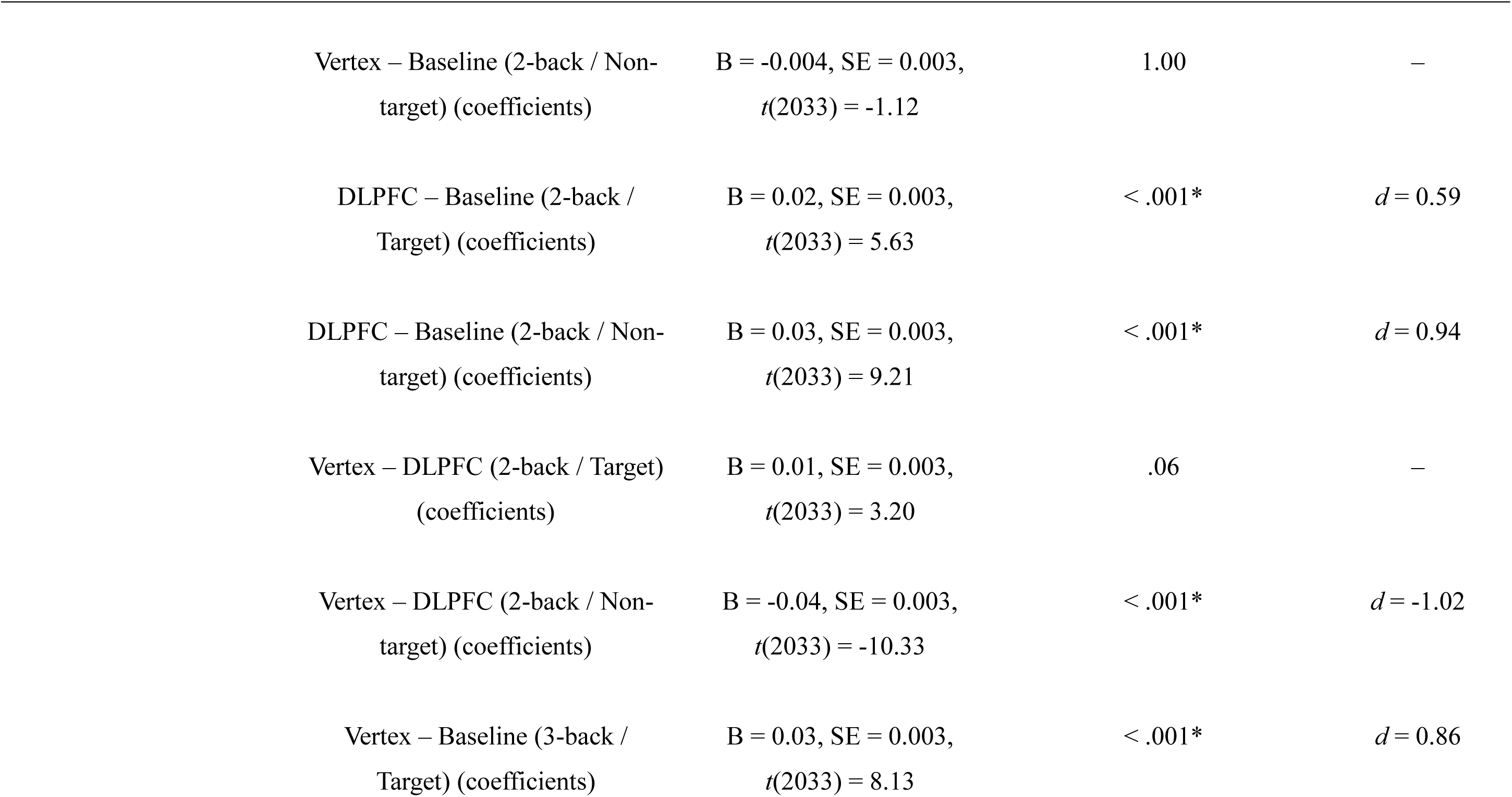
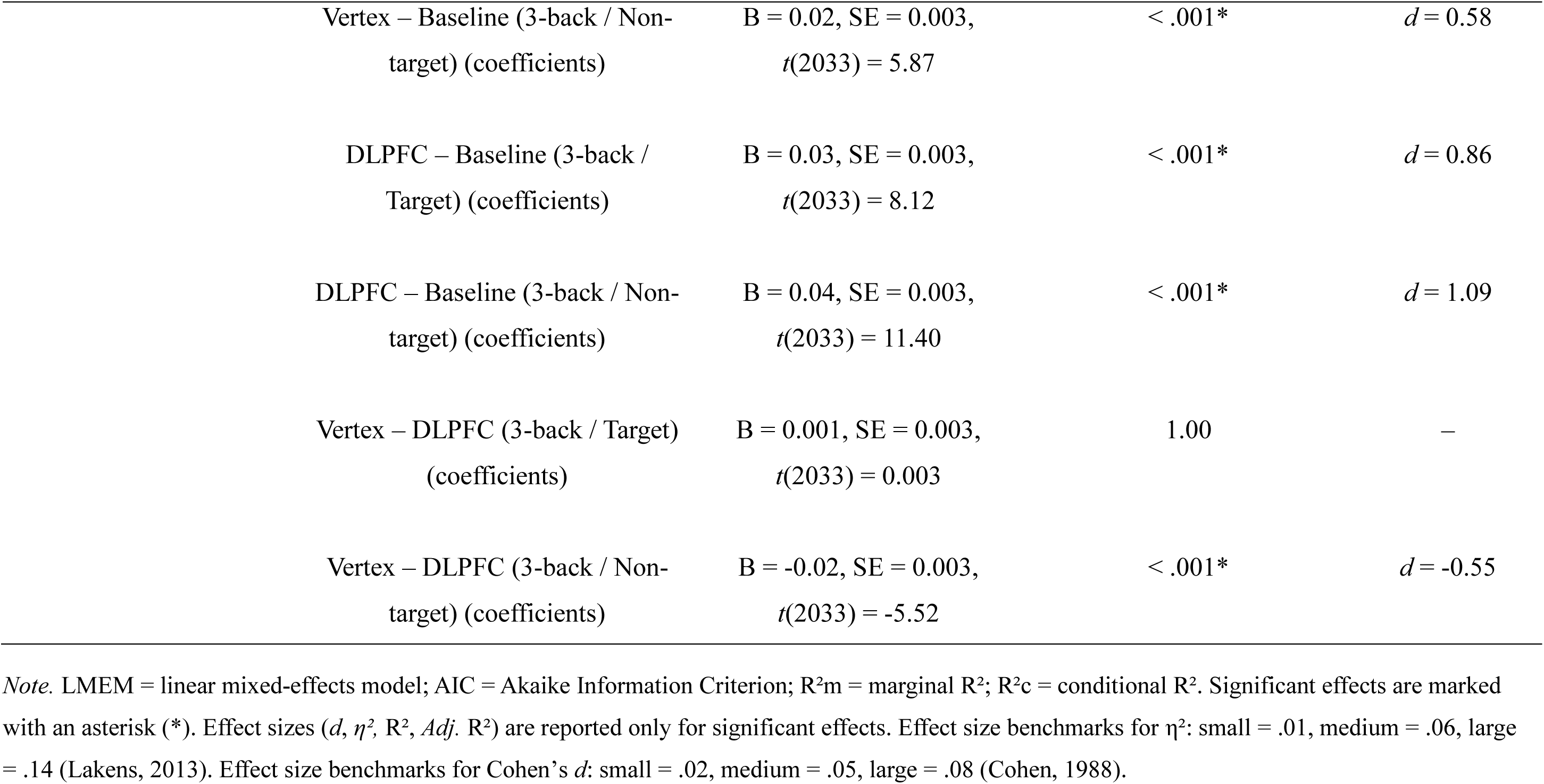
Summary of statistical analyses and results for the Visuospatial N-back – ERP latencies.

**Table S5.** Summary of cTBS-Vertex regression modelling results – Verbal N-back (accuracy)

| Model | Channels | Statistics |  | *p* | *Adj* R<sup>2</sup> |
| --- | --- | --- | --- | --- | --- |
|  |  | df | *F* |  |  |
| Scores ~ Cognitive load * (P200 + N200 + P300) | Frontal | (7, 32) | 1.19 | .34 | 0.03 |
| Scores ~ P200 * N200 * P300 | Frontal | (7, 32) | 2.46 | < .05* | 0.21 |
| Scores ~ P200 + N200 + P300 | Frontal | (3, 36) | 2.77 | .06 | 0.12** |
| ANOVA comparison of nested linear models | Frontal | (4, 32) | 2.00 | 0.12 | – |

*Note.* The best regression model was defined by a *p* value $\leq 0.05$ and more variance explained (*Adj.* R<sup>2</sup>). Only models based exclusively on frontal channels (Fz, F3, F4) are reported in the main text, given the established role of frontal regions in WM processes and their correspondence with the stimulation site (left DLPFC) used in the study. The best frontal model is marked with double asterisks (\*\*). A model comparison was performed to test whether including higher-order interactions among ERP components improved model fit. As comparison was not significant ( $p > .05$ ), a simpler additive model was preferred, following the principle of parsimony. Significant effects are marked with an asterisk (\*).

**Table S6.**
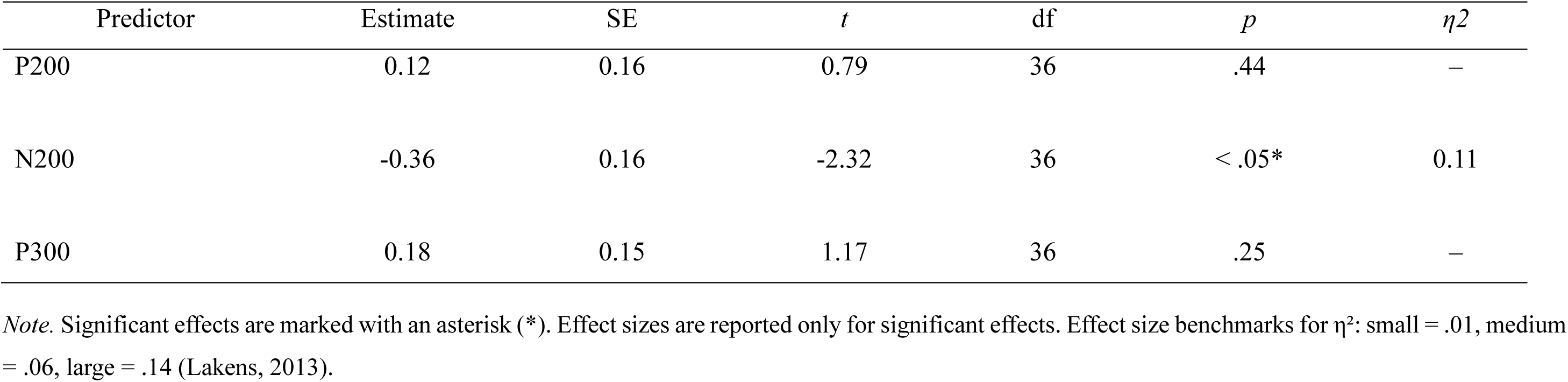
Statistics of the best regression model for the cTBS-Vertex condition – Verbal N-back (accuracy)

**Table S7.** Summary of cTBS-DLPFC regression modelling results – Verbal N-back (accuracy)

| Model | Channels | Statistics |  | *p* | *Adj R*<sup>2</sup> |
| --- | --- | --- | --- | --- | --- |
|  |  | df | *F* |  |  |
| Scores ~ Cognitive load * (P200 + N200 + P300) | Frontal | (7, 32) | 3.04 | .01* | 0.27** |

| Model | Channels | Statistics |  | *p* | *Adj* R<sup>2</sup> |
| --- | --- | --- | --- | --- | --- |
|  |  | df | *F* |  |  |
| Scores ~ P200 * N200 * P300 | Frontal | (7, 32) | 0.93 | .10 | 0.14 |
| Scores ~ P200 + N200 + P300 | Frontal | (3, 36) | 1.70 | .19 | 0.05 |

*Note.* The best regression model was defined by a *p* value $\leq 0.05$ and more variance explained (*Adj.* R<sup>2</sup>). Only models based exclusively on frontal channels (Fz, F3, F4) are reported in the main text, given the established role of frontal regions in WM processes and their correspondence with the stimulation site (left DLPFC) used in the study. The best frontal model is marked with double asterisks (\*\*). Significant effects are marked with an asterisk (\*).

**Table S8.** Statistics of the best regression model for the cTBS-DLPFC condition – Verbal N-back (accuracy)

| Predictor | Estimate | SE | *t* | df | *p* | $\eta^2$ |
| --- | --- | --- | --- | --- | --- | --- |
| Cognitive load | -0.20 | 0.33 | -0.59 | 32 | .56 | – |
| P200 | -0.76 | 0.22 | -3.41 | 32 | < .01* | 0.14 |
| N200 | 0.38 | 0.21 | 1.79 | 32 | .08 | – |
| P300 | -0.28 | 0.20 | -1.44 | 32 | .16 | – |
| Cognitive load x P200 | 0.99 | 0.33 | 2.96 | 32 | < .01* | 0.15 |
| Cognitive load x N200 | -0.86 | 0.35 | -2.48 | 32 | < .05* | 0.14 |
| Cognitive load x P300 | 0.47 | 0.35 | 1.34 | 32 | .19 | – |

*Note.* Significant effects are marked with an asterisk (\*). Effect sizes are reported only for significant effects. Effect size benchmarks for $\eta^2$ : small = .01, medium = .06, large = .14 (Lakens, 2013).

**Table S9.**
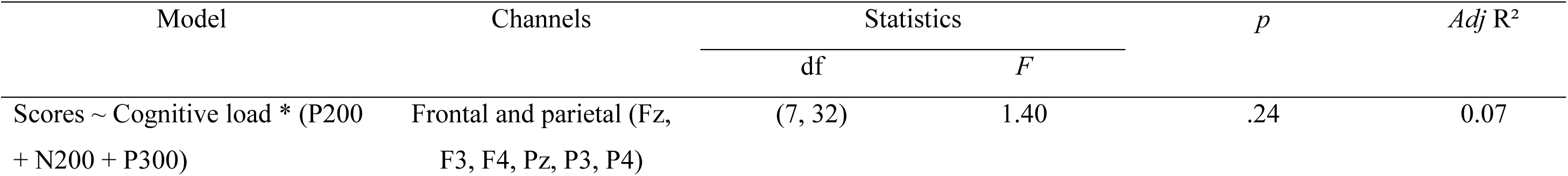
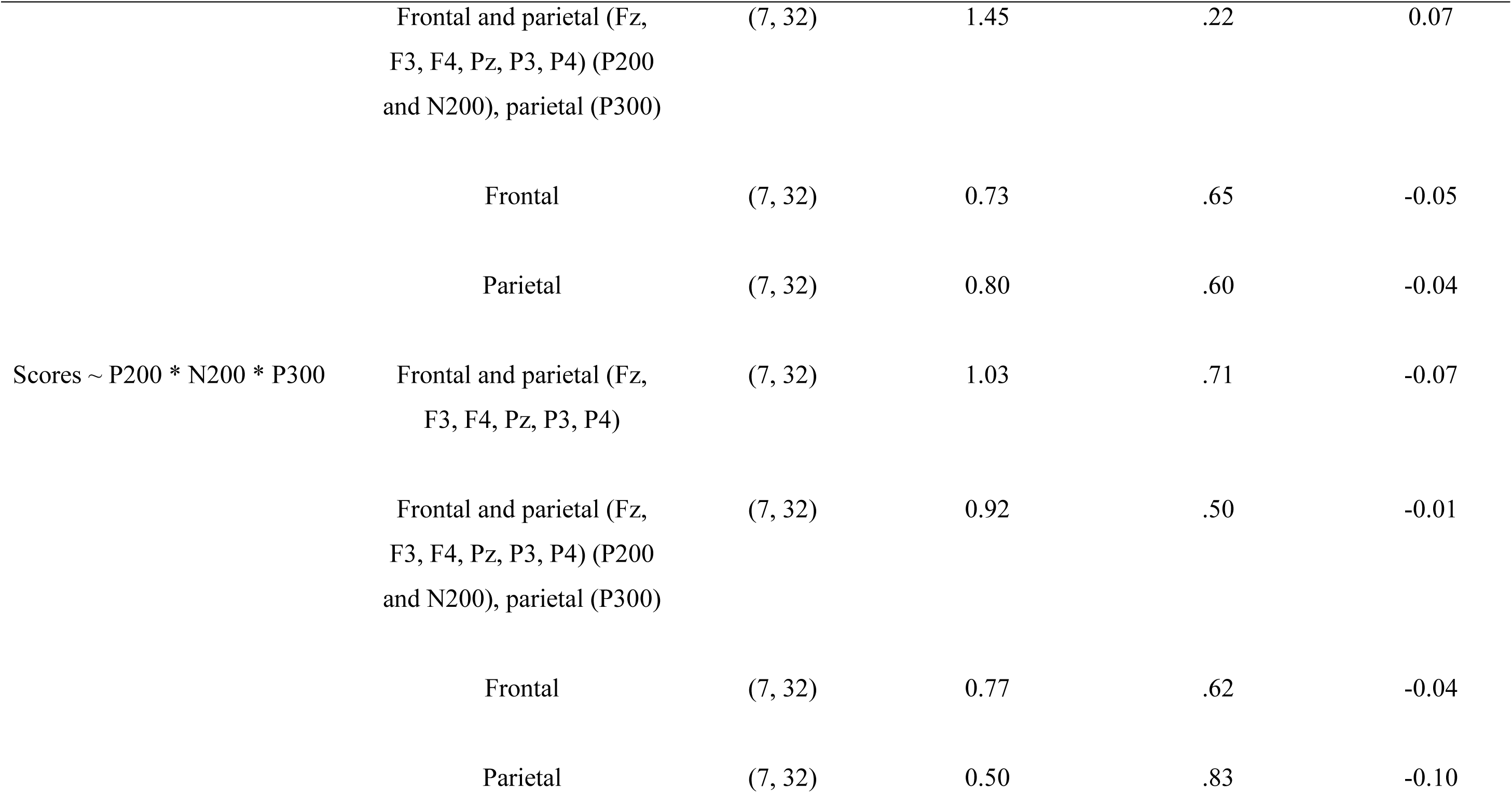
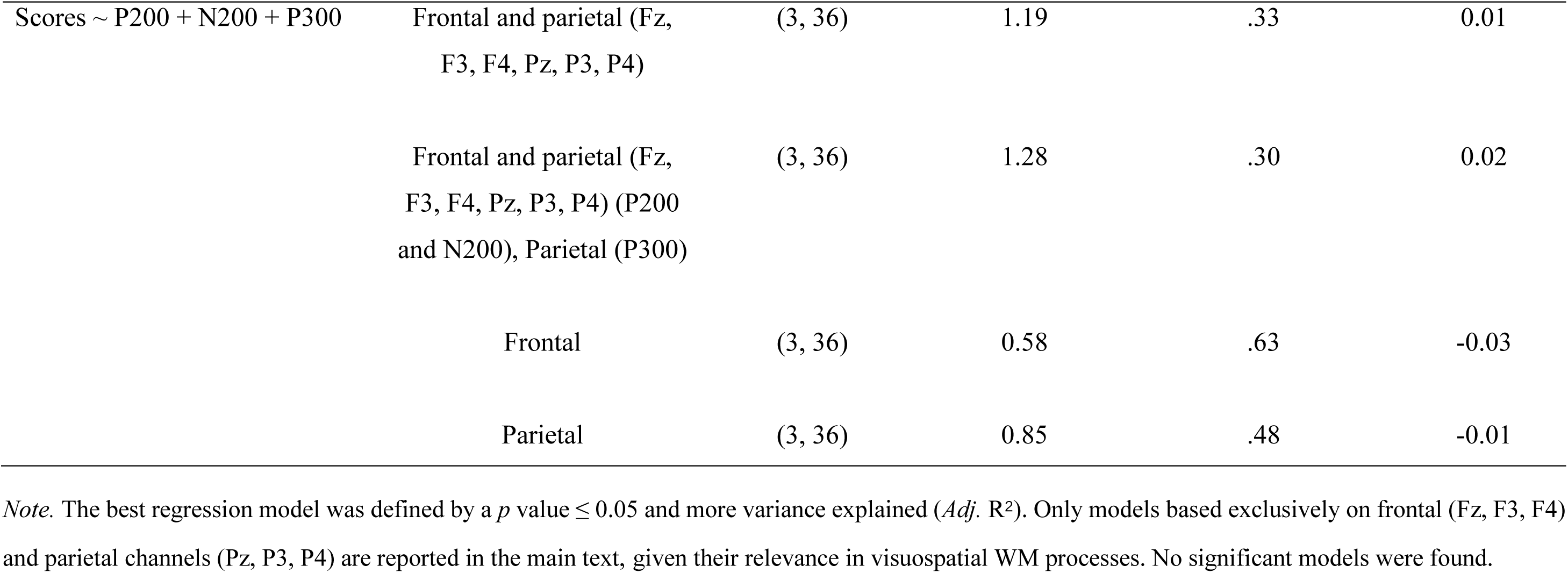
Summary of cTBS-Vertex regression modelling results – Visuospatial N-back (accuracy)

**Table S10.**
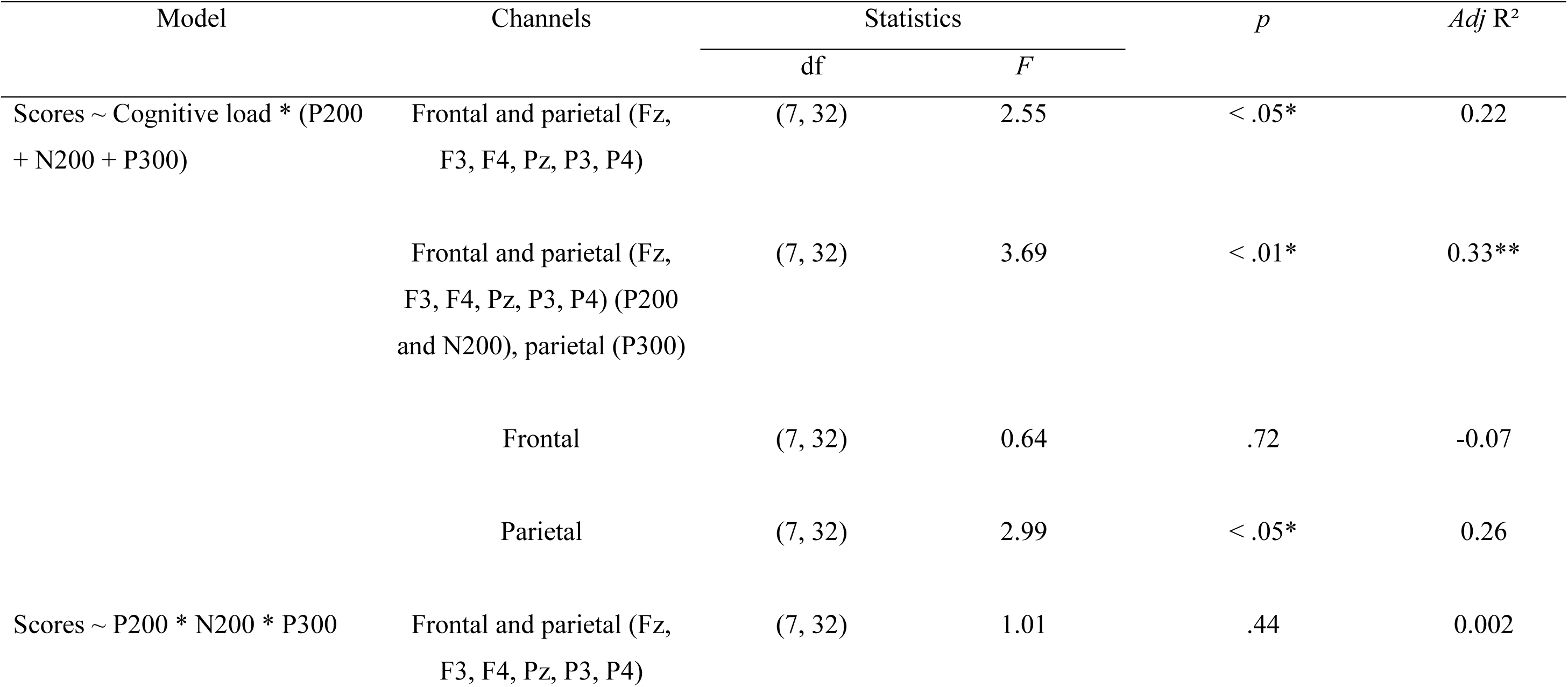
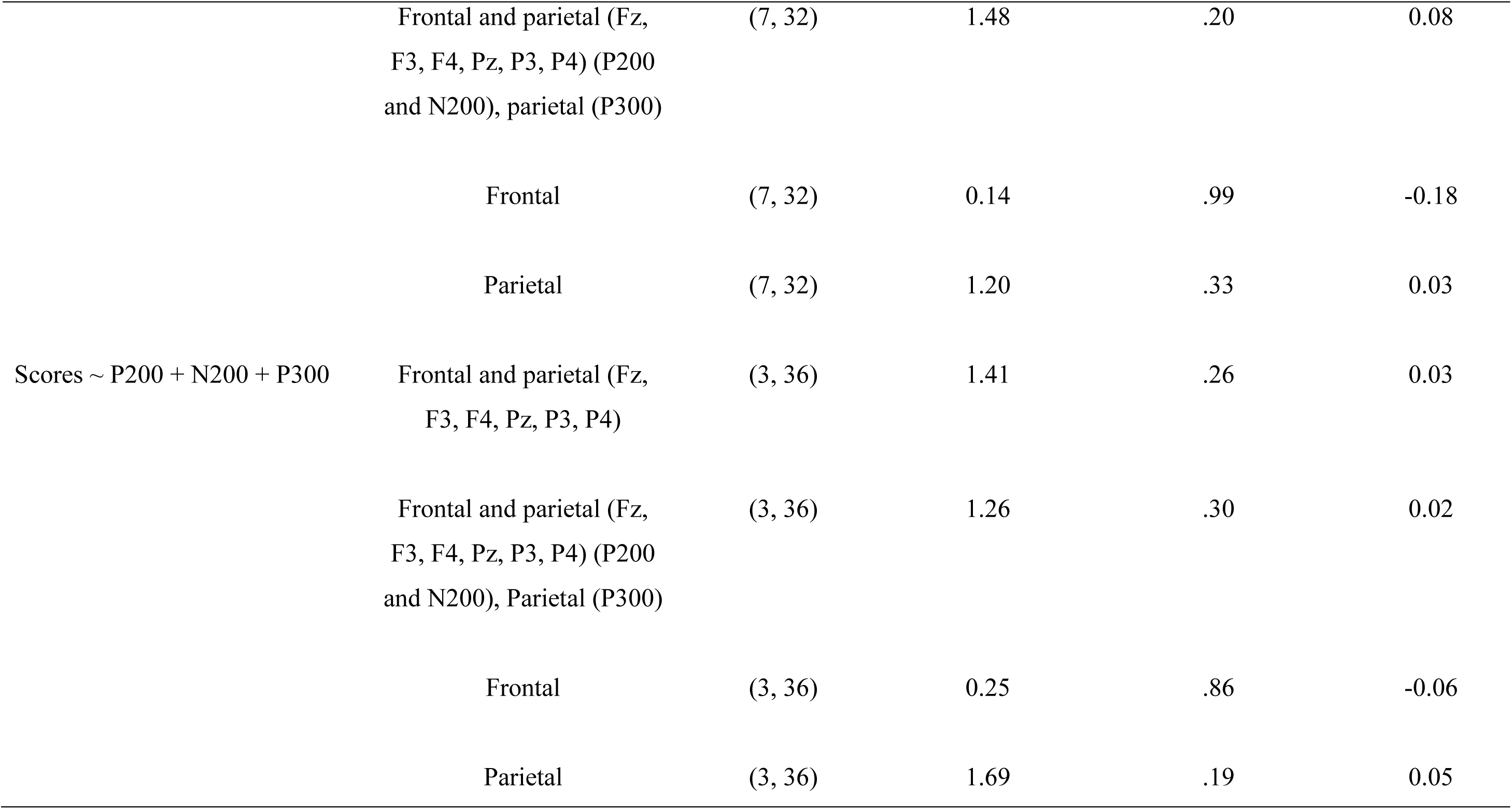
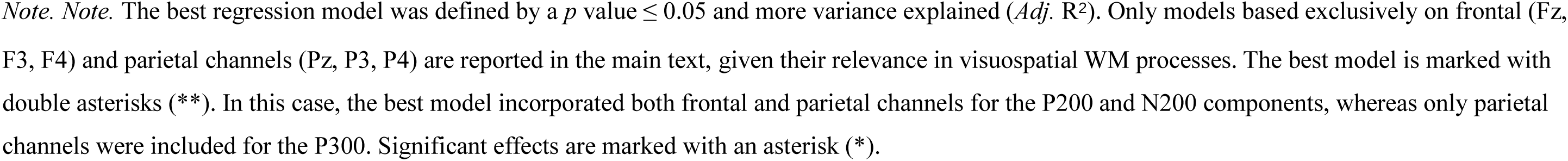
Summary of cTBS-DLPFC regression modelling results – Visuospatial N-back (accuracy)

**Table S11.** Statistics of the best regression model for the cTBS-DLPFC condition – Verbal N-back (accuracy)

| Predictor | Estimate | SE | $t$ | df | $p$ | $\eta^2$ |
| --- | --- | --- | --- | --- | --- | --- |
| Cognitive load | -0.73 | 0.27 | -2.64 | 32 | .01* | 0.16 |
| P200 | -0.19 | 0.17 | -1.12 | 32 | .27 | – |
| N200 | -0.57 | 0.17 | -3.32 | 32 | < .01* | 0.17 |
| P300 | 0.61 | 0.22 | 2.79 | 32 | < .01* | 0.003 |
| Cognitive load x P200 | -0.02 | 0.32 | -0.07 | 32 | .94 | – |
| Cognitive load x N200 | 0.47 | 0.28 | 1.68 | 32 | .10 | – |

| Predictor | Estimate | SE | *t* | df | *p* | $\eta^2$ |
| --- | --- | --- | --- | --- | --- | --- |
| Cognitive load x P300 | -0.95 | 0.30 | -3.18 | 32 | < .01* | 0.24 |

*Note.* Significant effects are marked with an asterisk (\*). Effect sizes are reported only for significant effects. Effect size benchmarks for $\eta^2$ : small = .01, medium = .06, large = .14 (Lakens, 2013).

